# A Wild Yeast Laboratory Activity: From Isolation to Brewing

**DOI:** 10.1101/698589

**Authors:** Amanda N. Scholes, Erik D. Pollock, Jeffrey A. Lewis

**Affiliations:** Cell and Molecular Biology Program, University of Arkansas, Fayetteville, AR; University of Arkansas Stable Isotope Laboratory, Fayetteville, AR; Department of Biological Sciences, University of Arkansas, Fayetteville, AR

**Author notes:** Corresponding author: Jeffrey A. Lewis, University of Arkansas, Department of Biological Sciences, 850 W. Dickson St., SCEN 601, Fayetteville, AR 72701.

## Abstract

Microbial fermentation is a common form of metabolism that has been exploited by humans to great benefit. Industrial fermentation currently produces a myriad of products ranging from biofuels to pharmaceuticals. About one third of the world’s food is fermented, and the brewing of fermented beverages in particular has an ancient and storied history. Because fermentation is so intertwined with our daily lives, the topic is easily relatable to students interested in real-world applications for microbiology. Here, we describe the curriculum for an inquiry-based laboratory course that combines yeast molecular ecology and brewing. The rationale for the course is to compare commercial *Saccharomyces cerevisiae* yeast strains, which have been domesticated through thousands of generations of selection, with wild yeast, where there is growing interest in their potentially unique brewing characteristics. Because wild yeast are so easy to isolate, identify, and characterize, this is a great opportunity to present key concepts in molecular ecology and genetics in a way that is relevant and accessible to students. We organized the course around three main modules: isolation and identification of wild yeast, phenotypic characterization of wild and commercial ale yeast strains, and scientific design of a brewing recipe and head-to-head comparison of the performance of a commercial and wild yeast strain in the brewing process. Pre and post assessment showed that students made significant gains in the learning objectives for the course, and students enjoyed connecting microbiology to a real-world application.

## Introduction

Microbial fermentation is a ubiquitous form of metabolism that has been exploited by humans for thousands of years (1-4). About one third of the world’s food is fermented (5), which of course has massive effects on global and local economies. Fermentation has a particularly rich history in the baking and brewing of alcoholic beverages, with the yeast *Saccharomyces cerevisiae* being among the oldest domesticated organisms (3, 6). While the first beer may have been brewed as long as 13,000 years ago (7), what we would now recognize as modern beer took shape in the Middle Ages, where malted barley was used as a source of fermentable sugars and hops were used as a bittering agent (8). During this time span, continuous selection of yeast in the brewing environment selected for a number of traits, including better utilization of wort carbon sources and increased fermentation efficiency. Modern brewing styles emerged from regional differences in brewing, and early brewers selected for yeast strains that complemented their brewing ingredients. For example, while the primary products of yeast fermentation are ethanol and carbon dioxide, a number of secondary products including esters and fusel alcohols are also produced that have unique flavor and aroma profiles (9). Certain beer styles (e.g. Belgian Lambic and German-style Hefeweizen) favor high levels of secondary fermentation products, while other styles favor little to none and consider these compounds to be “off flavors” (e.g. many Stouts and Amber Ales). The choice of yeast strain became a critical parameter for brewing design.

While brewers have most frequently used domesticated yeast strains, it’s becoming increasingly clear that wild yeast strains are important reservoir for traits important to industrial fermentations including brewing (10). This can include novel metabolic capabilities, such as the ability to ferment complex carbohydrates in wort, or the ability to produce novel flavor compounds (11). Because wild yeast are so easy to isolate, phenotype, and genotype, this provides a unique opportunity for undergraduates in laboratory courses to engage in inquiry-based research. As such, we designed a course around the microbiology of brewing a fermentation to provide a real-life application. We organized the course around three main modules: isolation and identification of wild yeast, phenotypic characterization of commercial and wild ale yeast strains, and scientific design of a brewing recipe and head-to-head comparison of the performance of a commercial and wild yeast strain in the brewing process.

### Intended Audience and prerequisite student knowledge

This course was designed to provide our senior Biology majors with an upper-level Microbiology laboratory course. This course also provides an opportunity for students to write a research paper that can satisfy our university’s writing requirement for graduation. Students should have some knowledge of molecular biology and biochemistry, particularly central metabolism and regulation of gene expression. As an upper-level course, students were required to have taken our sophomore-level Cell Biology and General Genetics courses and one of the associated introductory lab courses as prerequisites. While not required, we also suggested that our junior-level Prokaryote Biology course would be helpful.

### Learning time

The laboratory was structured as a three credit-hour full-semester course (16 weeks). The class was scheduled to meet twice a week for three hours, and the approximate length of each lab can be found in the instructor’s manual (Appendix 2). The majority of learning time and experiments took place in the laboratory. Some time outside of class was spent collecting wild yeast samples, reading relevant scientific literature, and completing assignments (laboratory notebooks, homework, oral presentation, and final research paper).

### Leaving objectives

The overall goal of the course is to provide both conceptual learning and hands-on laboratory skills. Upon completion of the course, students should be able to:

1. Summarize and discuss primary research literature.
2. Predict where wild yeast can be isolated based on the natural ecology of yeast, and explain how one can enrich for yeast from environmental samples.
3. Explain why and how ITS sequencing is used to determine fungal species, and analyze ITS sequencing data to assign the species of an unknown isolate.
4. Describe the primary and secondary products of yeast fermentation, and how differences in fermentative metabolism across yeast strains impact brewing.
5. Analyze yeast phenotypic data for traits relevant to brewing, and then use those data to predict brewing outcomes.
6. Explain the role of each ingredient and step in the brewing process, and scientifically design and implement a brewing protocol.

## Procedure

While we provide detailed student and instructor instructions in the Appendices, here we will briefly describe the main modules of the course (Figure 1):

**Figure 1:**
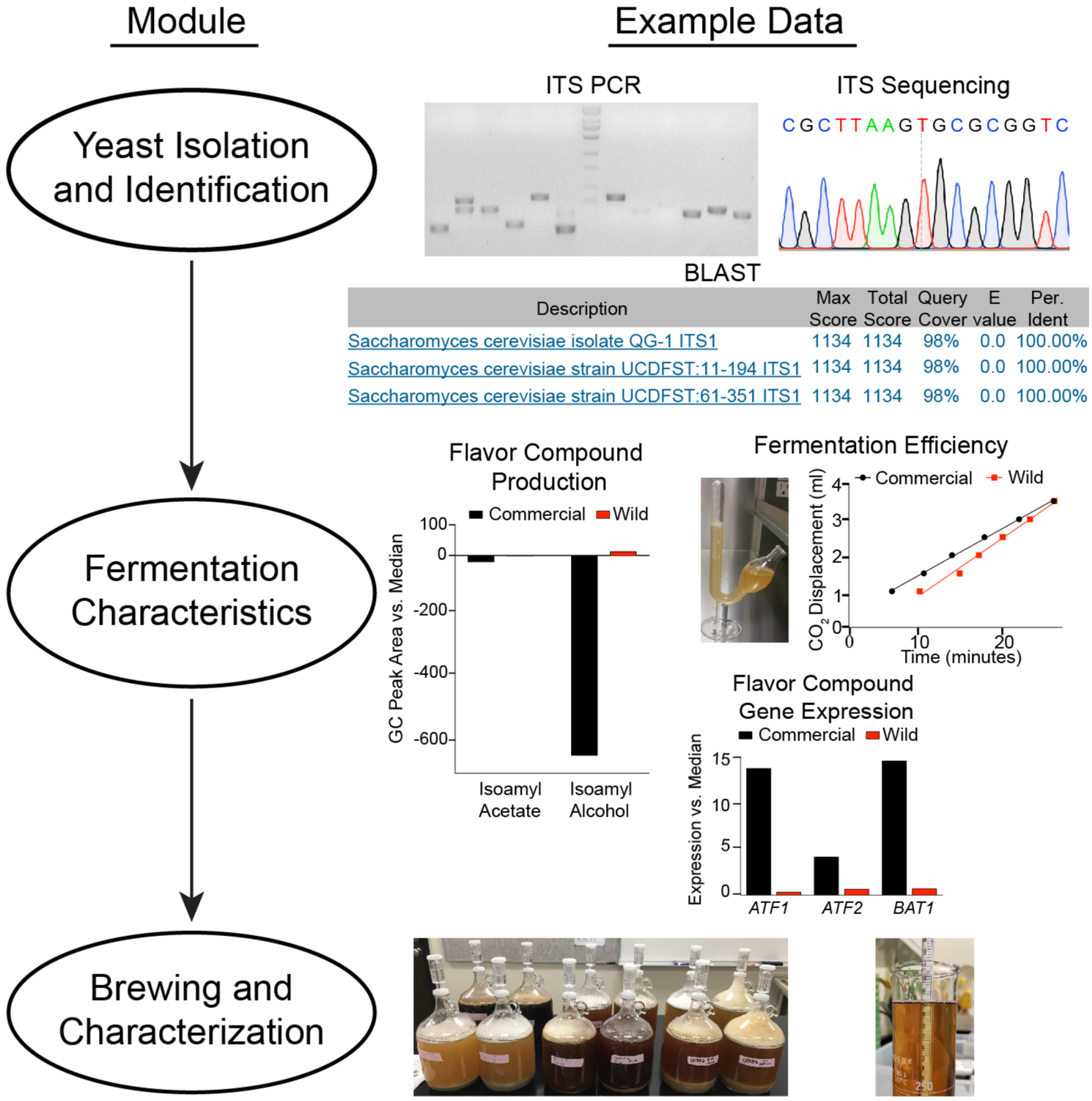
Flowchart displaying the different course modules and example data. ITS: internal transcribed spacer; PCR: polymerase chain reaction; BLAST: Basic Local Alignment Search Tool. qPCR: quantitative PCR; GC: gas chromatography.

### Wild Yeast Isolation and Identification

Yeast are ubiquitous in the environment and can be found on a number of substrates ranging from rotting fruit to soil to tree bark (12). For the first part of this course, students are given materials to sample from nature to isolate wild yeast. Students then place the samples in liquid media that enriches for budding yeast, and samples showing evidence of fermentation (gas bubbles) are plated to identify colonies consistent with those of yeast, which can be confirmed for the presence of budding yeast via microscopy. Following successful yeast isolation, students then perform DNA extractions, PCR and sequence the ITS/5.8S ribosomal DNA locus that is frequently used to differentiate yeast species (13), and then perform BLAST analyses to determine the species of their isolated yeast.

### Wild and Commercial Yeast Phenotypic Characterization

Students are then paired, and half of the class is charged with phenotypically characterizing different wild *S. cerevisiae* strain, and the other half of the class will characterize different commercial brewing strains. First the entire group learns how to “mash” malted grains together (which they will need to understand for the following module). The resulting wort from each group is then pooled and autoclaved to generate a standardized “beer media” to characterize all of the strains. Phenotypes for characterization include fermentation rate, quantitative PCR of mRNA levels for genes known to be responsible for ester and fusel alcohol production, and gas chromatography-mass spectrometry analysis of fermented beer media to directly quantify secondary metabolite levels.

### Wild and Commercial Yeast Brewing and Beer Characterization

The final module has student pairs join to form a larger group to design a brewing recipe where they will compete a wild and commercial *S. cerevisiae* strain head-to-head. Student pairs share their data with each other and then design a brewing recipe that fits the characteristics of one or both of their yeast strains. Here students gain hands-on experience for all of the major steps of brewing: mashing, boiling, and fermentation. During mashing, the grains are mixed with water and heated to a temperature that activates the alpha and beta amylases that naturally occur in malted grains. This leads to conversion of the grain starches into sugars that can be fermented by yeast (with the added wrinkle that alpha and beta amylases are most active at different temperatures, leading to different sugar profiles in the final wort depending on mash temperature). “Roasting” of the malted grains at different temperatures and times leads to lighter or darker malts (with darker malts having fewer active amylases and more Maillard products that are not fermentable). Following mashing, hops are generally added to the resulting sweet wort, which is then boiled. Boiling partially sterilizes the wort, and isomerizes hop alpha-acids leading to characteristic bitterness (with different varieties of hops containing differing amounts of alpha acids and other flavor compounds). Hop iso-alpha-acids also are bacteriostatic against many Gram-positive bacteria (14, 15). Finally, the wort is chilled, the yeast are “pitched”, and fermentation converts the wort sugars to mainly ethanol and CO_2_ along with secondary esters and alcohols.

For this course, we used the “brew in a bag” method, where the grains are placed in a bag that is submerged during the mashing process. Following mashing, the bag is simply removed and squeezed to drain the residual sweet wort. Then the sweet wort is brought to a boil for sterilization and hop additions, cooled to allow for yeast pitching, fermented for 3 weeks (typical for many ales), and finally bottle conditioned for 2 weeks. Following brewing, students measured their beers’ final gravities (to determine percent attenuation and alcohol percentage), color, bitterness, and secondary flavor compounds. Students also had the option of participating in a voluntary taste test of the final beers. Below is an example of a student-designed recipe built around a low-ester producing and highly fermentative yeast strain:

#### Recipe: Blood Orange Ginger American Ale

#### Ingredients

##### Grain (target original gravity 1.068)

4.96 lb 2-row U.S. Pale Malt

1.3 lb Briess Aromatic Munich Malt

0.43 lb Flaked Wheat

##### Hops (target IBU 86)

0.58 oz Citra – boiled for 60 min

##### Additives

1 Whirfloc Tablet (Irish moss; clarifying agent)– boiled for last 5 min

0.5 oz Blood Orange Extract – boiled for last 5 min

oz Sliced Ginger Root – boiled for last 10 min

##### Mashing

1. Heat 3.5 gallons of ultra-pure water in stockpot to 67°C.
2. Add all grain to the “brew bag” within the stockpot and mash at 67°C for 60 minutes.
3. Pull out brew bag and squeeze to drain excess wort. Discard spent grain

##### Boiling

4. Raise mash to a rolling boil.
5. Add Citra hops to a hop bag and add to boiling wort.
6. Incubate for 60 minutes.
7. With 10 minutes left in the boil, add 0.4 oz sliced ginger root.
8. With 5 minutes left, add 0.5 oz blood orange extract and 1 Whirfloc tablet.

##### Fermentation

9. Cool wort to near room temperature using a wort chiller.
10. Add cooled wort and 65.5 billion yeast cells to a 1-gallon fermentation growler.
11. Add airlock and fill with water diluted Star San sanitizer.
12. Transfer 250mL of the remaining wort to graduated cylinder and measure initial gravity with a hydrometer
13. Place fermentation growlers in a dark area at room temperature for 3 weeks.

##### Bottle Conditioning

14. Add 6.6 ml 50% glucose (priming sugar for carbonation) to sterilized 16 oz amber swing-neck bottle.
15. Auto-siphon the beer into a sterile 16oz amber bottle.
16. Transfer 250 ml of the remaining beer to graduated cylinder and measure final gravity with a hydrometer.
17. Incubate at room temperature for 2 weeks in the dark to carbonate the beer.

### Materials

Materials are listed for a class of 24 students working individually for the initial yeast isolation, and then in pairs for the subsequent experiments. Materials (including media recipes) and equipment are listed in Appendix 2.

### Student instructions

The student manual can be found in Appendix 3. Students were required to maintain a lab notebook with detailed rationale, methods, results, and discussion sections. An example lab notebook entry can be given to students to serve as a guide (Appendix 4). The notebook was collected three times during the 16-week course. Students were also responsible for preparing a ten-minute fermentation-related oral presentation, along with a final term paper describing their scientifically-designed brewing recipe in journal article format.

### Faculty instructions

Detailed faculty instructions for lab activities can be found in the instructor manual (Appendix 2). Lab lectures and active learning activities (clicker questions and group discussions) can be found in Appendix 1. Instructor materials for all graded assignments including associated rubrics can be found in Appendix 4.

### Outcomes and issues for discussion with students

Because this is a research-based course, anticipated outcomes are not guaranteed. Not all students are guaranteed to isolate yeast for molecular characterization. Those students should be provided with a wild yeast isolate, either from another classmate who isolated more than one unique strain, or from the instructor. Likewise, there is no guarantee that the class will isolate enough wild *S. cerevisiae* strains for subsequent experiments, so the instructors should be prepared to supply wild *S. cerevisiae* strains as a backup. Wild yeast strains can be ordered from the ARS Culture Collection (https://nrrl.ncaur.usda.gov), but the corresponding author (Dr. Jeff Lewis) is happy to send wild *S. cerevisiae* strains upon request. It is helpful to cryopreserve all positively screened wild *S. cerevisiae* strains so that they can be used in future classes if necessary.

### Suggestions for determining student learning

A pre- and post-laboratory exam and survey (Appendix 5) were administered to students. The 15-question exam consisted of an equal number of multiple choice, true-false, and short answer questions. The 5-question survey measured student perceptions of proficiency using a Likert-like scale. We did not use quizzes, midterms, or a final exam to assess student learning, though those could certainly be implemented. The ability to summarize and discuss the primary literature was assessed via homework assignments and a short (10-12 min) oral presentation. For each module, laboratory notebooks were graded to assess student learning. A final paper in the form of a primary research article was used as an additional summative assessment of student learning.

### Safety issues

Students must demonstrate competency with BSL1 safety procedures before working with unknown samples that require BSL2 precautions. Because this in an upper-level course that requires prerequisite BSL1 level lab activities, students were mostly familiar with BSL1 precautions. Nonetheless, students received important safety training on proper BSL1 and BSL2 procedures, and were required to demonstrate proficiency with BSL1 procedures before performing BSL2 procedures including safe handling of potentially pathogenic unknown organisms (16). Students were required to wear personal protective equipment (gloves, lab coat, eye protection) at all times, and received instructions for how to minimize aerosolizing cultures. All bench surfaces and objects on the laboratory bench were disinfected after each class with 70% ethanol. The instructors were responsible for autoclaving all plates and contaminated materials after every class according to the minimal standards set by the ASM Biosafety Guidelines (16). All chemicals in this course are low risk biohazardous agents except for methylene blue, hydrochloric acid, iodine, and iso-octane, which were discarded according to the institutional biohazard waste disposal guidelines.

All ingredients used for brewing were food grade, and brewing was conducted in a space safe for food handling. While many different types of wild yeast can be used for brewing, we were cautious to only use wild *Saccharomyces cerevisiae*. This activity and the associated research was submitted to the University of Arkansas IRB Committee (Protocol No. 1807133914) and determined to be exempt. The course also included optional tours of a local craft brewery (Core Brewing in Springdale, AR) and a local homebrew store (Steve’s Brew Shop in Fayetteville, AR), as well as an optional taste test of the final beers. We recognized that tasting of alcohol beverages is a potentially sensitive subject, so we worked closely with the university administration to ensure that we complied with all university regulations and guidelines. We came up with the following guidelines for beer tasting: 1) Tasting is entirely optional. Any students who do not wish to participate do not have to, and the tasting will have no impact on student grades. 2) Only students 21 years of age or older may participate in tasting. A valid photo ID with birth date will be required. IDs will be checked by the trained staff at Core Brewing. 3) Tasting will only occur at Core Brewing. There will be no tasting of alcoholic beverages on campus. 4) Tasting will be through the sip and spit method only. There will be no drinking of the beer. 5) Students must sign a waiver that includes the above information, as well as a statement that they will act responsibly.

## Discussion

### Field testing

This class was developed, and field tested through two years as an upper-level research-based undergraduate course at the University of Arkansas (23 students in 2017, and 24 students in 2018). Students worked independently for yeast isolation, in pairs for yeast sequencing and characterization, and in groups of four (two pairs) for brewing. Discussions within and between groups were encouraged. For yeast isolation, 37 / 47 students successfully isolated wild budding yeast. Based on ITS sequencing, 6 / 37 isolates were *S. cerevisiae*. Several other species were identified including *S. paradoxus, S. cariocanus, Pichia* species (*P. kudriavzevii, P. kluyveri, P. fermentans, P. terricola*), *Meyerozyma caribbica, Lachancea fermentati, Wickerhamomyces anomalus, Kodamaea ohmeri*, and *Debaryomyces sp*. Further characterization only proceeded with wild *S. cerevisiae* strains. While all-grain brewing may seem intimidating for novices, the “brew in a bag” method dramatically simplifies the process, and works extremely well for the small volumes being brewed in the course. Neither the instructors nor most of the students had any experience brewing, but every group in both student cohorts was able to successfully brew beer.

In the second offering of the course, we changed the focus of the brewing module to be more “yeast centric.” We did this by having pairs of lab partners—each working with either a commercial brewing strain or a wild *S. cerevisiae* strain—scientifically design a single brewing recipe to compete the yeast strains. This allowed each group of students to predict how the final beer would change depending on the properties of the yeast, and then test these predictions in the final characterization of the beer.

Overall, student feedback on the course was highly positive. Anonymous online evaluations rated the course very highly on a 1 (very poor) through 5 (excellent) Likert-like scale, with a 2017 rating of 4.80 / 5 (compared to a departmental mean of 3.96) and a 2018 rating of 4.79 / 5 (compared to a departmental mean of 3.83). Student comments pointed to a particular appreciation of connecting molecular biology to real-world applications.

Examples include:

- *I was able to learn about genetics through real life situations, and to apply what I learned, in a way that made much more sense than my general genetics course ever did.*
- *This class is a great example of helping students to understand complex concepts by utilizing an interesting life-application.*
- *Great reminder of some biology concepts that did not seem applicable to real life when taught in another course.*

### Evidence of student learning

Student learning was assessed using a variety of methods (Table 1). Take-home problem sets (Appendix 4) were used to assess understanding of the assigned readings. Lab notebook entries were used to assess students’ abilities to understand the rationale for their experiments as well as their design, analyses, and interpretations. Students were evaluated on their ability to present short (10-15 min) mini-lectures on their choice of topics related to microbial fermentation. Last final written report in the format of a primary research article was used to assess students’ abilities to synthesize what they learned. Rubrics can be found in Appendix 4.

**Table 1:**
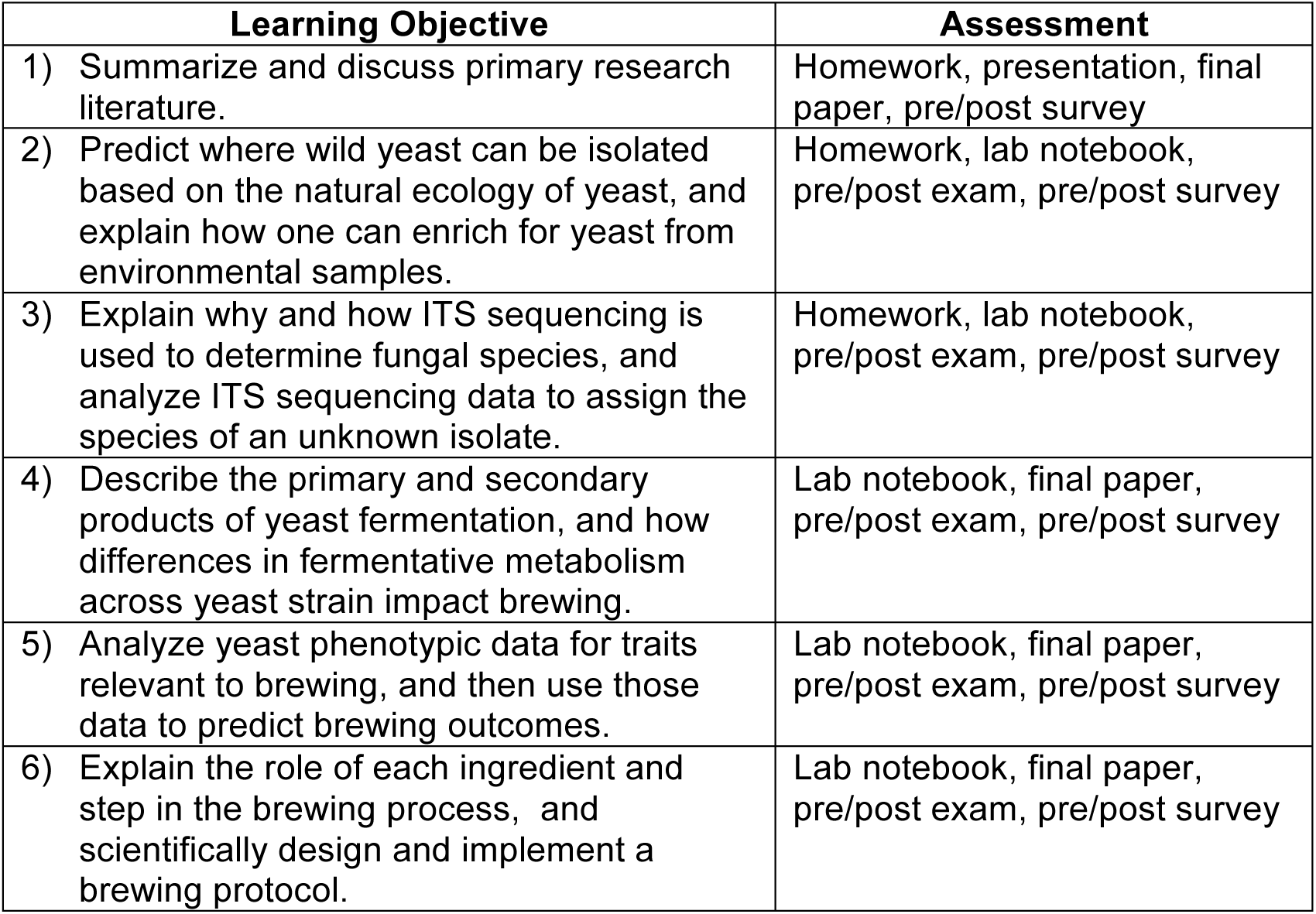
Learning objectives and their corresponding methods of assessment.

We measured changes in student learning with pre- and post-tests, and we assessed changes in student perceptions of their skills and knowledge with pre- and post-surveys (see Appendix 5 for exam and survey questions). The average pre-test score was 26% correct, which rose to 67% following participation in the course (Figure 2). This was statistically significant (*p* = 6 x 10^-14^, two-tailed unpaired Mann-Whitney U test), and of large effect (Cliff’s delta = 1). Additionally, students showed significant increases in learning for the majority of the questions (Figure 3). We should note that formal assessment in the course did not include any exams, so these gains are more likely to reflect long-term understanding instead of short-term memorization. Students also self-reported their perceptions of competency on pre- and post-surveys. Following the course, students showed significantly higher confidence in their abilities to isolate wild yeast from nature, use molecular biology and phylogenetics to identify yeast species, describe the major steps in brewing, and brew beer on their own (Figure 4). Coming into the class, students felt confident with reading scientific articles, though they may have still showed a small gain in confidence following the course (*p* = 0.08, two-way ANOVA, Fisher’s LSD).

**Figure 2:**
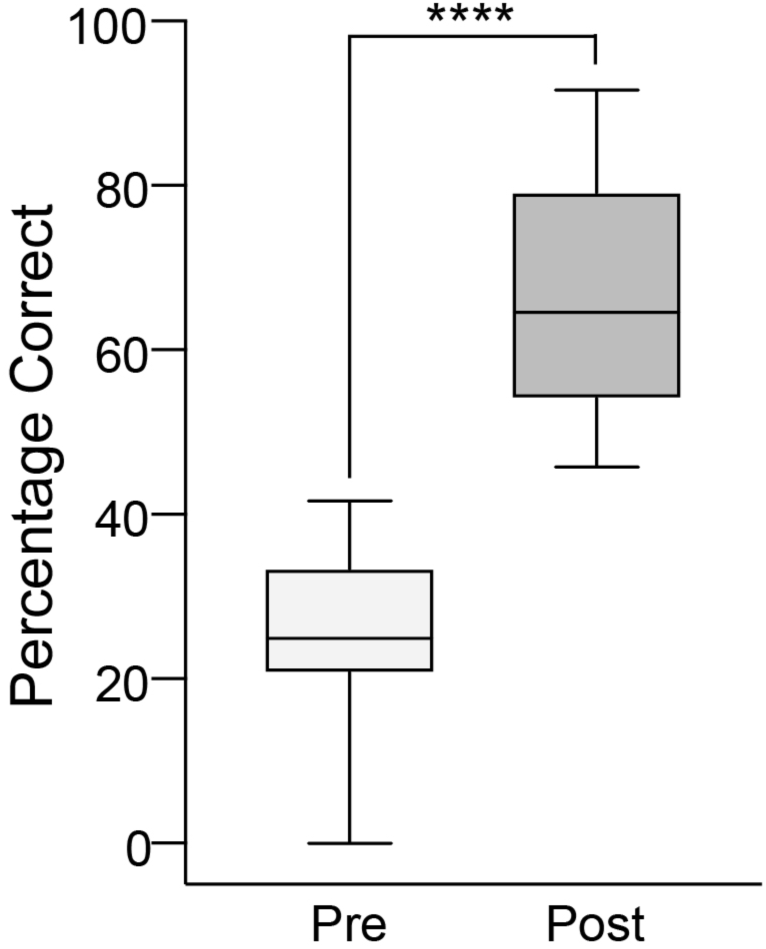
Pre- and post-exams show significant gains in student learning. The boxplot depicts the median and interquartile range, and the whiskers depict the range. **** *P* = 6 x 10^-14^, two-tailed unpaired Mann-Whitney U test.

**Figure 3:**
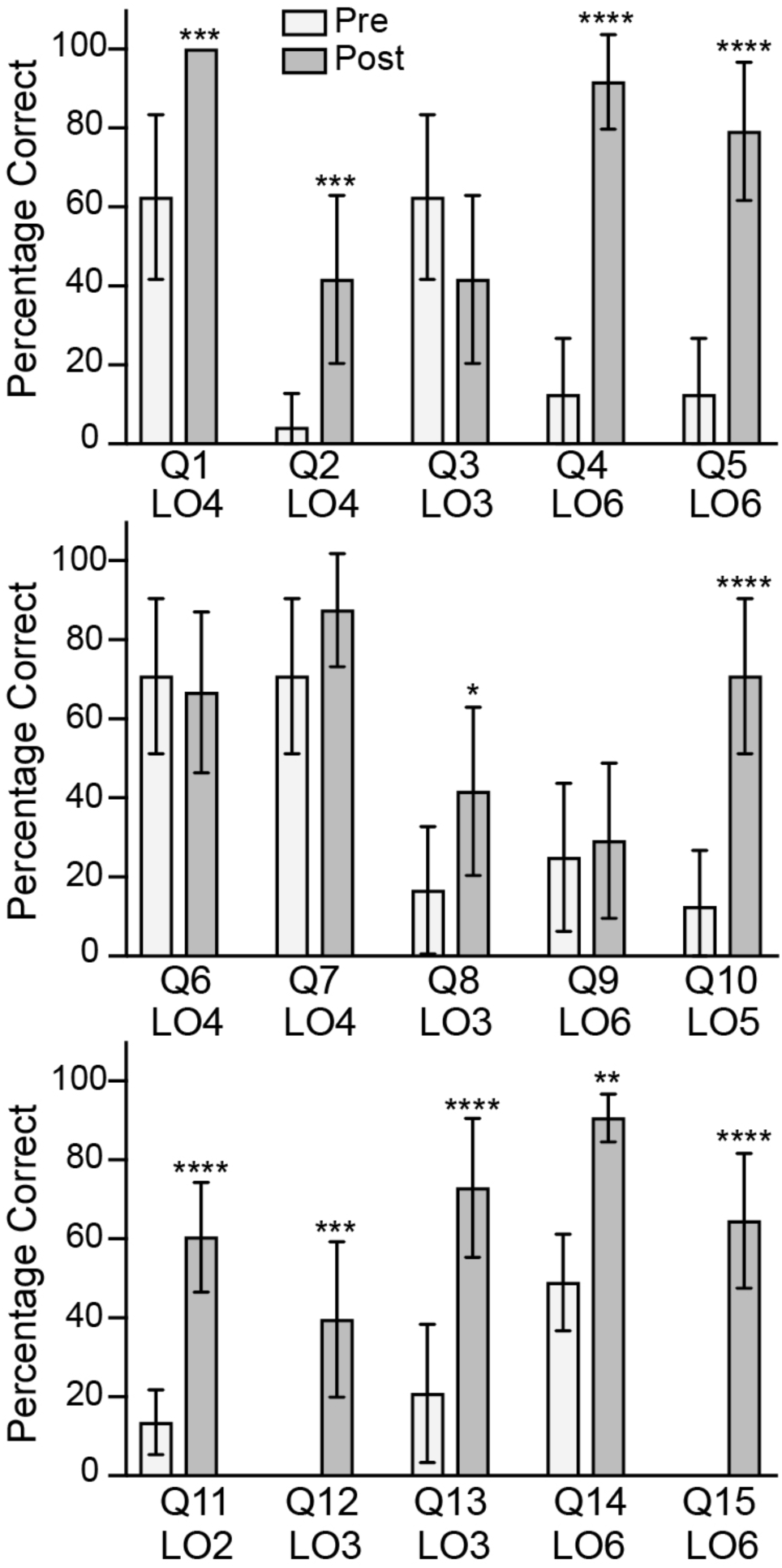
Individual item responses for pre- and post-exam scores. Exam questions (Q1 – Q15) can be found in Appendix 5. LO denotes the learning objectives. * *P* < 0.05; ** *P* < 0.01;*** *P* < 0.001; **** *P* < 0.0001, two-way ANOVA, Fisher’s LSD test.

**Figure 4:**
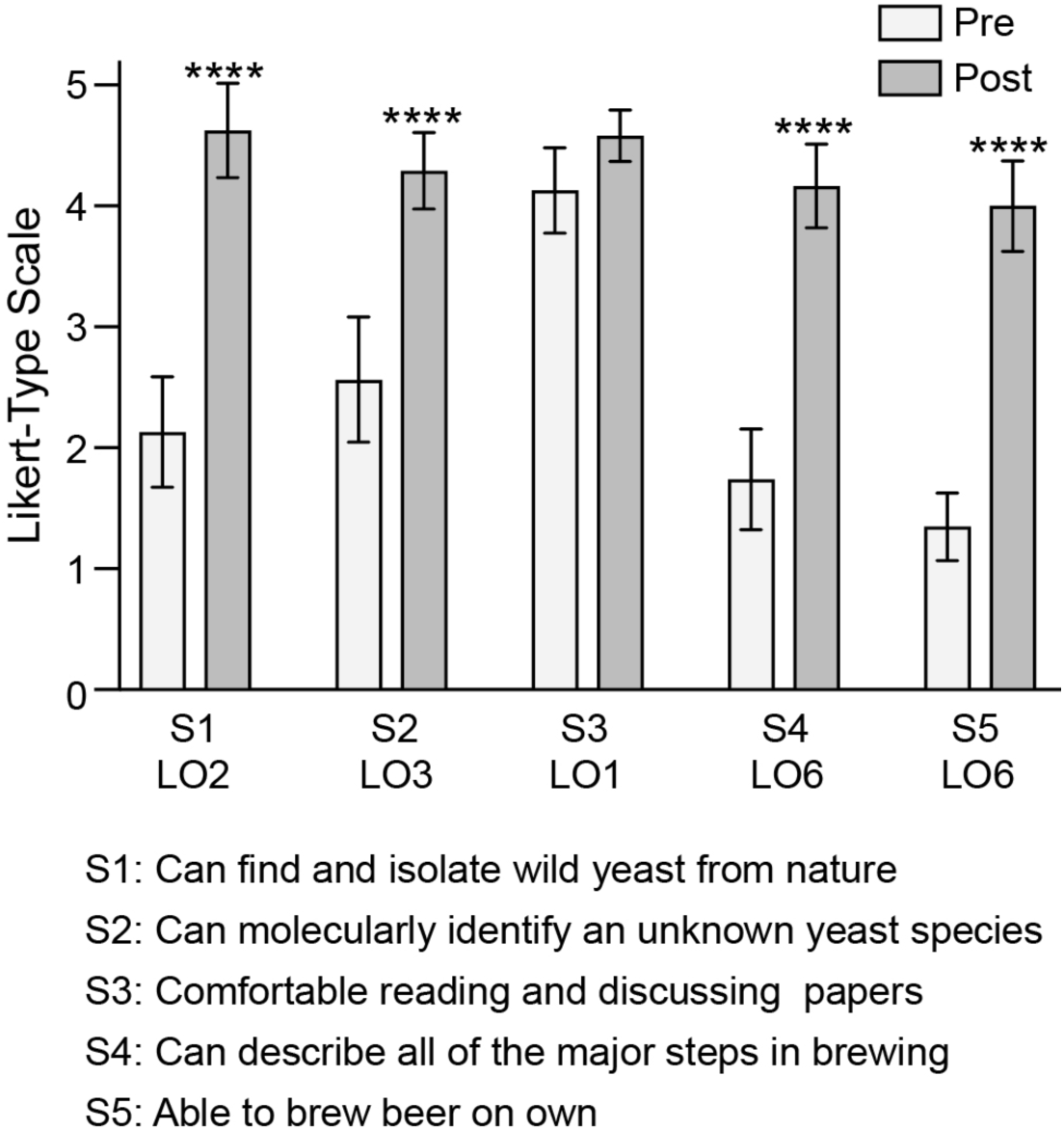
Pre- and post-survey shows increase in self-reported perceptions in student ability. Survey questions (S1 – S5) can be found in Appendix 5. LO denotes the learning objectives. **** *P* < 0.0001, two-way ANOVA, Fisher’s LSD test.

### Possible modifications

This course was designed and offered twice as an upper-level course that met twice a week for one semester. There are several modifications that could be included for a shorter course. For example, the yeast isolation can be shortened by the instructor plate or streak colonies from fermentation-positive cultures. Additionally, the brewing module can be shortened by using commercial malt extracts instead of mashing whole grains. Optional activities that could be omitted include a guest lecture from a local craft brewer, and tours of both a local craft brewery and homebrew store.

One of the optional modules we included was strain characterization of flavor compound formation (e.g. volatile esters and fusel alcohols), which we did both at the gene expression level via quantitative real-time PCR (qPCR) and directly via gas chromatography-mass spectrometry (GC-MS). We understand that some instructors may not have access or funds to include these modules. One cheaper alternative to qPCR would be semi-quantitative PCR (17). An alternative to GC-MS is sensory analysis, where students can be trained to identify esters and fusel alcohols by taste (individual flavor standards may also be purchased from FlavorActiV to facilitate compound identification).

## Supplemental Material

Appendix 1 – Instructor Lab Lectures

Appendix 2 – Instructor Lab Manual

Appendix 3 – Student Manual

Appendix 4 – Assignments and Rubrics

Appendix 5 – Pre- and Post-Exam and Survey

## Acknowledgements

Funding for this project was supported in part by National Science Foundation grant no. IOS-1656602. We are thankful to Ron Schmidt of Core Brewing and Distilling, LLC. for providing a guest lecture and a guided brewery tour. We also thank Steve Wilkes for giving a guided tour of Steve’s Brew Shop. The authors declare that there are no conflicts of interest.

