## Appendices for "A Wild Yeast Laboratory Activity: From Isolation to Brewing"

#### Supplementary Material

Appendix 1: Instructor Lab Lectures

Appendix 2: Instructor Lab Manual

Appendix 3: Student Manual

Appendix 4: Assignments and Rubrics

Appendix 5: Pre and Post-Exam and Survey

**Lab Lectures: Table of Contents**

| Week | Topics | Page Number |
|---|---|---|
| | Readings | 1 |
| | Schedule for Assignments and Presentations | 2 |
| Week 2 | Introduction to Fermentation and Yeast Life History | 3-5 |
| Week 2 | How to Read a Scientific Paper and Wild Yeast Ecology | 6-10 |
| Week 3 | Yeast Enrichment and Isolation | 11 |
| Week 4 | Phylogenetics, PCR and Sequencing | 12-13 |
| Week 5 | BLAST, Malting, Tips on How to Give a Scientific Presentation | 14-19 |
| Week 6 | Yeast Pitching and Fermentation Traits | 20-22 |
| Week 7 | Yeast Fermentation (By)Products and Gene Expression | 23-26 |
| Week 8 | Quantitative PCR and Gas Chromatography-Mass Spectrometry (GC-MS) | 27 |
| Week 9 | Malts and Hops, qPCR Analysis | 28-29 |
| Week 10 | Beer Styles and GC-MS Analysis | 30-36 |
| Week 11 | Designing a Brewing Recipe (Placing Ingredient Orders) | 37 |
| Week 12 | Brewing Day and Brewery Tour | 38 |
| Week 13 | Homebrew Store Tour | 39 |
| Week 14 | Hop Microbiology | 40 |
| Week 15 | Beer Characterization | 41 |
| Week 16 | Student Presentations | 42 |
| Week 17 | Taste Test and Final Paper Due | N/A |

**Note: For each lab lecture, slides are numbered with the header/title of the slide and bullet points for the slide text and/or citations for images used. Red text indicates active learning exercises (e.g. clicker questions, small (4-6 person) group discussions that are then shared with the larger class). We are happy to send PowerPoint slides of the lecture materials. Please email the corresponding author (Jeff Lewis) at.**

#### Readings

**Week 1:**
Piskur J, Rozpedowska E, Polakova S, Merico A, Compagno C. 2006. *How did Saccharomyces evolve to become a good brewer?* Trends in Genetics. 22(4): 183–186

**Week 2:**
Sniegowski PD, Dombrowski PG, Fingerman E. 2002. *Saccharomyces cerevisiae and Saccharomyces paradoxus coexist in a natural woodland site in North America and display different levels of reproductive isolation from European conspecifics.* FEMS Yeast Research. 1(4): 299–306

**Weeks 3 – 4:**
Sylvester K, Wang QM, James B, Mendez R, Hulfachor AB, Hittinger CT. 2015. *Temperature and host preferences drive the diversification of Saccharomyces and other yeasts: a survey and the discovery of eight new yeast species.* FEMS Yeast Research. 15(3): pii: fov002

**Weeks 5 – 6:**
Bamforth CW. 2009. *Current perspectives on the role of enzymes in brewing.* Journal of Cereal Science. 50: 353–357

**Week 7:**
Pires EJ, Teixeira JA, Brányik T, Vicente AA. 2014. *Yeast: the soul of beer's aroma--a review of flavour-active esters and higher alcohols produced by the brewing yeast.* Applied Microbiology and Biotechnology. 98(5): 1937–1949

**Weeks 8 – 9:**
Saerens SM, Verbelen PJ, Vanbeneden N, Thevelein JM, Delvaux FR. 2008. *Monitoring the influence of high-gravity brewing and fermentation temperature on flavour formation by analysis of gene expression levels in brewing yeast.* Applied Microbiology and Biotechnology. 80(6): 1039–1051

**Weeks 10 – 14:**
Pires E and Brányik T. 2015 *Biochemistry of Beer Fermentation, Chapter 1: An Overview of the Brewing Process.*

Schönberger C and Kostelecky T. 2011. *125th Anniversary Review: The Role of Hops in Brewing*. Journal of the Institute of Brewing. 117(3): 259–267

#### Schedule for Student Assignments

**Week 2:**
Homework #1 Due

**Week 4:**
Homework #2 Due

**Week 5:**
Notebook #1 Due

**Week 6:**
Homework #3 Due

**Week 8:**
Homework #4 Due

**Week 9:**
Oral Presentations (2)

**Week 10:**
Oral Presentations (4)

**Week 11:**
Homework #5 Due
Oral Presentations (2)

**Week 12:**
Notebook #2 Due

**Week 13:**
Paper Draft Due
Oral Presentations (2)

**Week 14:**
Oral Presentations (2)

**Week 15:**
Oral Presentations (6)

**Week 16:**
Notebook #3 Due
Oral Presentations (6)

**Week 17 (Final Day)**
Final Paper Due

**Week 1 Lab Lecture Slides**

**Day 1:**

1) Lab safety Discussion

2) Fermentation
    - Louis Pasteur: "the consequence of life without air"
    - broad definition: a form of metabolism in which the end products can be further oxidized

3) Fermentation is something microbes can do…
    - 1789: Antoine-Laurent de Lavoisier shows that sugar gets fermented into ethanol and carbon dioxide
    - 1837-1838: Charles Cagniard de la Tour, Theodor Schwann, and Friedrich Kützing independently show that yeast is a living organism that reproduces by budding
    - 1857: Louis Pasteur shows that fermentation depends upon microorganisms (lactic acid bacteria and milk souring)

4) Why do cells ferment?
    - Image: pathways of lactate and ethanol fermentation highlighting re-oxidation of NADH

5) Why not ferment all the time?
    - **Small groups discuss and report: answers included fermentation having a lower ATP yield, and that fermentation products may be mildly toxic and thus reduce fitness compared to respiration**
    - Table: substrates, products, and energy yield of fermentation vs. respiration

6) Examples of Fermentation products...
    - Image from Biology, Seventh Edition Neil Campbell and Jane Reece. Figure 9.27 (includes propionate->Swiss cheese, lactate->yogurt, ethanol->wine/beer, acetone->nail polish remover, and acetate->vinegar

7) Fermentation can be remarkably complex
    - Figures 1 and 2 from Clark DP. FEMS Micro. Rev. 1989. 63: 223-234

8) What does fermentation do to food?
    - Preservative (ethanol, pH)
    - Changes the composition

9) Example: Making yogurt from milk
    - Lactose -> lactic acid via
    - What happens to proteins when they are acidified?
    - Lactic acid causes milk proteins (e.g. casein) to denature

10) Example: Making yogurt from milk
    - Image of casein bundles in milk that become a gel in yogurt
        - Acidification causes casein bundles denature and separate
        - These rebind to each other as a "gel" matrix that traps water.

**Week 1 Lab Lecture Slides**

**Day 2:**

11) Life History and Biology
    - ....small animal which sips sugar through its snout, and excretes alcohol from its gut and carbonic acid from its urinary organ...
        - Friederich Wohler and Justus Liebig, 1839 Annalen der Pharmacie
        - Satirical images from Enari. J. Inst. Brewing. 1995. 101: 3-33 and Majno. J. Inf. Dis. 1991. 163: 937-945
    - 1878: The term "enzyme" was proposed by Wilhelm Kuhne, which is Greek for "in yeast"
    - 1938: Franz Meyen names yeast *Saccharomyces*, which is Greek for "sugar fungus"

12) Brewing Yeast Species
    - Ale strains are *Saccharomyces cerevisiae*
        - polyploid (or aneuploid) genome structures
        - poor sporulators, low spore viability, poor mating
        - relatively high genetic diversity
        - grow well at higher temperatures, high ethanol production
        - ferment at 18°C – 22°C

    - Lager strains are *Saccharomyces pastorianus*
        - hybrid between *S. cerevisiae* and *S. eubayanus*
        - extremely poor sporulators, low spore viability, poor mating
        - 15th century Bavaria
        - ferment at lower temperatures (6°C – 11°C)

13) *Saccharomyces eubayanus*
    - first discovered in Patagonia by Chris Hittinger and colleagues
    - was found in the sugar-rich fruiting bodies of a fungal parasite of Southern Beech trees
    - Did yeast make a trek from the Southern Andes to Bavaria?
    - has been isolated from other locations and is hypothesized to have first originated in Asia
    - Images from: https://news.wisc.edu/500-years-ago-yeasts-epic-journey-gave-rise-to-lager-beer/

14) Yeast Life History and Biology
    - Image of Tree of Life Wikimedia Commons (NASA Astrobiology Institute): Phylogenetic tree.svg

15) Yeast Cellular Anatomy
    - Image of Yeast Cell Structure from Fig 1.1 of Stewart and Russell. 1998. An Introduction to Brewing Science & Technology - Brewer's Yeast.

16) Yeast Life Cycle
    - Image: Figure 1 from Hanson and Wolfe. 2017. Genetics. 206: 9-32

17) **Clicker Question: Image of growth curve showing lag, log phase, diauxic shift, post- diauxic shift, and stationary phase, with question: What are yeast using for carbon and energy during exponential phase?**
    A. **glucose**
    B. ethanol
    C. amino acids
    D. the bodies of their fallen brethren

18) **Clicker Question: Image of growth curve showing lag, log phase, diauxic shift, post- diauxic shift, and stationary phase, with question: What are yeast using for carbon and energy during the post-diauxic phase?**
    A. glucose

**B. ethanol**
C. amino acids
D. the bodies of their fallen brethren

**19) Small group discussion: What is the cell doing during the diauxic shift?**

- **Remodeling gene expression (e.g. they switch ON the respiration genes and switch OFF the fermentation genes)**

**Week 2 Lab Lecture Slides**

**Day 1:**

1) How to Read a Scientific Paper
    - First, how not to read a scientific paper:
        - read from the Abstract to References without reflecting on content
        - read it like a textbook and take the authors' word for it

2) Guiding Principles:

    - Scrutinize everything:
        - the questions being asked
        - the data
        - the interpretations
        - the authors might be wrong about something!
    - Look stuff up:
        - you will likely need to constantly Google terms, and use a textbook or review article to understand the background
        - it will get easier over time and you will be a better scientist for doing it 3)
    - "A month in the laboratory can often save an hour in the library" (Frank Westheimer)

3) Types of Scientific Papers

    - Review articles:
        - summarize a bunch of papers as they relate to a topic
        - can be thought of as specialized textbook chapters
        - caution: everyone has biases, so look at multiple reviews to get a broad perspective

    - Research Articles:
        - aka primary literature or simply "the literature"
        - often the life work of some poor student, so read critically, but be respectful in critiquing

4) Research Article Format
    - Abstract: the authors' summary of everything
    - Introduction: relevant background, rationale, questions
    - Results: actual data (raw or processed), figures, tables
    - Discussion: authors' interpretation of data, context
    - Acknowledgements: funding and people
    - References: make sure they cited your obviously very important paper
    - Supplement: detailed methods, data files, extra figures

5) Abstract:
    - use it to identify whether you want to read a paper – once it is in your pile, skip it

6) Introduction: Ask the following questions…
    - Why is this topic important to study?
    - Why should you care?
    - What are the big picture questions in the field? (Not necessarily what the authors are going to tackle)
    - What previous work as been done? Were there limitations? What do the authors propose as the next step?
    - What are the specific questions that the authors seek to answer? What approaches are they taking?
    - When reading, write ~5 bullet points to summarize these points

7) Methods: often the hardest section to read, focus on the general approach
    - It often helps to draw a cartoon or flowchart for each major experiment
    - Ask if the methods are appropriate for answering the questions being asked, if there may be better methods, or if there are caveats

8) Results: carefully scrutinize here, for each figure/table/result…
    - Write down the authors' question that the experiment is try to answer
    - Is this experiment sufficient to answer the question?
    - Interpret the data yourself (not what the authors say it means)
        - It's okay to change your mind later, but train yourself to interpret first
        - It's okay if you do not understand something—bring these questions to class
        - Sometimes I will have no clue either! Sciencing is hard and sometimes things do not easily fit into our understanding of the world

9) Discussion: bullet points again
    - What do the authors think their results mean? Do you agree? Are there alternative interpretations or hypotheses?
    - Are there any weaknesses? Places where more experimental support is needed?
    - What is the next step? Has this research opened up new questions?

10) Lab Notebook: Overall Organization
    - Table of Contents:
        - should contain experiment titles and page numbers
    - Each Experiment:
        - should be dated
        - should contain page numbers that match to the table of contents

11) Lab Notebook! Organization for Each Experiment:
    - Title:
        - e.g. Isolating Wild Yeast from Arkansas
        - should be informative but concise
    - Rationale/Purpose/Background:
        - what is the motivation for the experiment or the research question
        - any hypotheses regarding the overall research question
        - this does not have to be very long, maybe 3-6 sentences, but should make the overall rationale clear

12) Lab Notebook! Organization for Each Experiment:
    - Materials and Methods
        - these must be very detailed
        - should include a complete list of the materials used
        - should include a step-by-step methodology
        - must be complete enough that you could give this to a scientist and have them get the same results
    - Results
        - record all of your raw data
        - this includes quantitative (numerical data) and observational data written out in sentence form
        - when in doubt, record it

13) Lab Notebook! Organization for Each Experiment:
    - Conclusions
        - include you interpretations of the results
        - did the results support the hypotheses, and if not, what are the alternative explanations?
        - were there mistakes or pitfalls in the methods, and if so, what could be changed?

14) Wild Yeast Isolation
- List of sampling substrates from the students:
  - **Mushroom**
  - **Rotting berries**
  - **Tomatillo, jalapeno**
  - **Sap**
  - **Soil**
  - **Bark**
  - **Tree nut**
  - **Tree-associated, berries, misc. fruit, mushrooms, non-rotting fruit**

- **Small group discussion: hypothesize where we are most likely to isolate wild yeast:**
  - **tree or rotting fruit, berries, berries or rotting fruit, rotting fruit**

15) Yeast Enrichment Media Recipe
- 0.3% yeast extract
- 0.3% malt extract
- 0.5% bacto peptone
- 1% sucrose
- 8% ethanol
- 1 µg / ml chloramphenicol
- dd H2O
- pH brought down between 5-6 with HCl

**16) Discussion: what is each component doing?**
- 0.3% yeast extract: **vitamins, trace nutrients**
- 0.3% malt extract: **carbon source**
- 0.5% bacto peptone: **protein/amino acid source**
- 1% sucrose: **fermentable carbon source**
- 8% ethanol: **inhibits non-tolerant microbes**
- 1 µg / ml chloramphenicol: **inhibits bacteria**
- dd H2O
- pH brought down between 5-6 with HCl: **pH optimum for yeast**

17) What should be the next step?
- **plate for single colonies**
- **colony morphology**
- **verify that they are budding**

**Week 2 Lab Lecture Slides**

**Day 2:**

1) Yeast whole genome duplication (WGD)
   - Image: Figure 1 of Thompson et al. 2013. eLife. 2:e00603, annotating Crabtree-positive (ferments in the presence of $O_2$), intermediate, Crabtree-negative, and obligate respiratory species

2) Many brewing strains are hybrids
   - Image: Figure 1 of Boynton and Greig. 2014. Yeast. 31(12): 449-462

3) Phenotypic variation in diverse yeast strains
   - Image: Figure 2 of Kvitek et al. 2008. PLoS Gen. 4(10): e1000223

4) The Basic Ingredients of Brewing •
   - water
   - malted grain (most commonly barley)
   - hops (flower that adds bitterness, aromas, and antibacterial properties)
   - yeast
   - adjuncts (other grains, fruit, spices)

5) **Small Group Discussion: what stresses or conditions are important for brewing?**
   - **pH**
   - **Pressure**
   - **Light**
   - **Nutrients**
   - **Contamination**
   - **Temperature**
   - **Osmotic (sugars in the wort)**

6) What stresses or conditions are important for brewing?
   - ethanol: yeast need to be able to produce enough and they need to remain viable at the end of the fermentation
   - osmotic: this is especially important if you are using a more concentrated wort (aka "high gravity") •
   - pH: CO2 will acidify the brew
   - temperature: fermentation produces heat, lagering is done at low temperature, yeast are generally stored in the cold
   - oxygen: yeast need some to grow; too little and they will not divide, too much and they will make biomass instead of ethanol
   - Yeast stress will influence secondary metabolite production, which will influence flavor

7) What stresses or conditions are important for brewing?
   - Image: Figure 1 from Bleoanca and Bahrim. 2013. Rom. Biotech. Lett. 18: 8559-8572

8) **Small Group Discussion: methods to assess yeast happiness?**
   - **Measure nutrient levels in media**
   - **Look at pH of the media**
   - **CO2 output (ethanol too)**
   - **Growth of the yeast**
   - **Viability of the yeast**

9) Methods to assess yeast happiness?
   - growth
     - liquid or plates

- viability
    - colony forming units
    - viable stains (colorimetric or fluorescent)
        - microscopy
        - flow cytometry
- gene expression
    - reporters (e.g. LacZ or GFP fusions)
    - real-time quantitative PCR (RT qPCR)
    - Northern blots
    - global (RNA-seq)
- performance
    - directly measuring ethanol and “off” compounds
        - gas chromatography and mass spectrometry

**Week 3 Lab Lecture Slides (no Day 2, holiday)**

**Day 1:**

1) Plate streaking
   - Image of streaked plate from Wikimedia Commons: https://commons.wikimedia.org/wiki/File:Legionella_Plate_01.png
   - Video: https://www.youtube.com/watch?v=mCOtWztCObY

2) **Group discussion: how to identify yeast species**
   - **Species concept and viable offspring (need known tester species)**
   - **Use a known strain/species and PCR**
   - **Colony morphology (characteristics)**

3) What to look for on plates...
   - discrete off-white/cream colored colonies

**Week 4 Lab Lecture Slides**

**Day 1:**

1) Molecular Phylogenetics (Carl Woese)
   - Image: Figure 1 from Petrov et al. 2014. PLoS One. 9(2): e88222
   - Key Step: select a DNA region that is homologous, or similar across species due to common ancestry
   - Ribosomal RNA (rRNA)
   - Ideal gene for phylogenetic studies because it ...
     - is an essential gene that is present in all organisms.
     - is a common target for sequencing studies; large database for comparisons.
     - contains sites that are relatively conserved (stems) and sites that are more free to vary (loops).

2) Polymerase Chain Reaction (PCR) (Kary Mullis)
   - molecular xerox—rapidly amplify (make copies) of DNA
   - key innovation: discovery of heat-stable DNA polymerase from a bacterium isolated from Yellowstone
     - Taq polymerase from the bacteria *Thermus aquaticus*

3) Polymerase Chain Reaction (PCR)
   - components for PCR:
     - buffer (pH optimized for Taq)
     - dNTPs (nucleotides used to synthesize DNA)
     - Mg++ (nucleotides complex with Mg++ in the active site of Taq)
     - template (DNA to be amplified)
     - primers (small, synthesized DNA oligomers, specific to the template)
     - Taq polymerase

   - automated "thermal cycling"
     1) denaturation (high heat renders DNA single stranded)
     2) annealing (primers bind to the template at lower temp.)
     3) extension (Taq extends DNA starting at the primers)

4) PCR: Polymerase Chain Reaction
   - Image: from Wikimedia Commons: https://commons.wikimedia.org/wiki/File:Polymerase_chain_reaction.svg
   - 30 cycles > 230 (> 1 billion) copies from a single DNA molecule

5) Internal transcribed spacer (ITS) region
   - Image: Box 1 from Nature Reviews Immunology 14, 405–416 (2014), point out ITS1 forward (at 3' end of 18s) and ITS4 reverse (at 5' end of 25s) primers

6) Looking at the DNA extraction protocol...
   - What is SDS and why is it necessary? **helps lyse yeast cells and denatures DNA-binding proteins**
   - What is LiOAc and why is it necessary? **helps DNA to precipitate in solvent**
   - What is EtOH and why is it necessary? **precipitates DNA**
   - What is TE and why is it necessary? **solubilizes the DNA; Tris buffers the solution and EDTA chelates divalent metal ions necessary for DNases to work**

**Week 4 Lab Lecture Slides**

**Day 2:**

1) (Sanger) DNA sequencing: chain termination method
   - Basic Principle: DNA extension is stopped by di- deoxynucleotides (ddNTPs)
   - drew a "normal" dNTP and ddNTP, note that that dNTP has a hydroxyl that can attack the next triposhpate, while a ddNTP does not and is "stuck"

2) (Sanger) DNA sequencing: chain termination method
   - components for Sanger sequencing:
     - buffer
     - DNA polymerase (generally a modified Taq)
     - DNA to sequence (e.g. purified PCR product)
     - a single specific primer
     - normal dNTPs
     - small amount of ddNTPs conjugated to a fluorescent dye • aka "dye-terminators" (e.g. A = green, T = red, C = blue, G = yellow)
   - one-tube sequencing reaction (cycling similar to PCR)
   - **Clicker Question: Does the sequencing reaction exponentially amplify the DNA?**
     A. Yes
     **B. No**

3) Automated Sequencing
   - DNA is placed on gel or capillary
   - Fragments move off gel in size order; pass through laser beam
   - Color each fragment fluoresces is recorded on printout
   - Image: from Wikimedia Commons: https://commons.wikimedia.org/wiki/File:Sanger-sequencing.svg

4) Small group discussion: looking at the PCR purification protocol...
   - Why do we need to "clean" our PCR product? What would happen if we used the "unclean" PCR product for sequencing? **primers and dNTPs would still be present and interfere with the sequencing reaction**
   - The "Binding Buffer" contains buffer (pH ~6), and high concentrations of chaotropic salts and isopropanol. The DNA binds to a silica membrane. How does the "Binding Buffer" facilitate binding to the column membrane? **The combination of low pH and chaotropic salts that disrupt hydrogen bonds decreases DNA interaction with water and increases interaction with the silica.**
   - The "Wash Buffer" contains a small concentration of a buffer to keep the pH at ~6 (e.g. Tris or HEPES) and high concentrations of ethanol (70%). What is the purpose of the ethanol in the wash buffer? **Ethanol maintains the DNA interaction with the silica, while washing away the salts**
   - The "Elution Buffer" is simply 10 mM Tris-HCl pH 8. How does this facilitate elution of the DNA from the column? **High pH and an aqueous environment favors DNA interactions with water instead of the silica membrane**

5) After looking at the gel…
   - What could cause the appearance of two bands? **hybrid species, or mixed culture**
   - Why might PCR have failed to yield a band
     - technical reason(s)? **inhibitors in DNA, made master mix incorrectly…**
     - biological reason(s)? **ITS too variable for primers**

**Week 5 Lab Lecture Slides**

**Day 1 (Computer Lab):**

1) Internal transcribed spacer (ITS) region
   - Image: Box 1 from Nature Reviews Immunology 14, 405–416 (2014), point out ITS1 forward (at 3’ end of 18s) and ITS4 reverse (at 5’ end of 25s) primers

2) DNA Sequencing
   - Performed using 300ng PCR product 8μM ITS1 primer (remember that the PCR of ITS was performed using ITS1 and ITS4)
   - Download ITS PCR Nanodrop Readings (record the results in your lab notebook)
   - Open folder “ITS Sequences” and click on the link to your strain to download the sequence file (.ab1 format)
   - Click to open—should automatically open in SnapGene Viewer

3) Interpreting a Sequencing Chromatogram
   - Showed raw chromatogram image from one of the students

4) Interpreting a Sequencing Chromatogram
   - Showed smoothed algorithm with basecalls from same chromatogram

5) Interpreting a Sequencing Chromatogram
   - Qualitative Measures of Quality
     - well-defined, single peaks
     - peaks are evenly spaced
     - lack of background

6) Interpreting a Sequencing Chromatogram
   - Quantitative Measures of Quality (Phred Scores)
     - examines the peaks around each base to assign a quality score
     - quality score is logarithmically related to base-calling error probability
     - Table of Phred scores and base call accuracy (https://en.wikipedia.org/wiki/Phred_quality_score)

7) Interpreting a Sequencing Chromatogram
   - Quantitative Measures of Quality (Phred Scores)
     - Can click on “show quality values” button to quickly scan for low- quality calls
     - This can be important for when you are trying to determine whether a difference in a database match is *real* vs. a sequencing error
     - If you hover over a base, you will get the actual Phred score
     - The beginning of the sequence generally has low quality bases and is not used (trimmed)
     - Depending on stringency, Phred 20 or Phred 30 are used as a cutoff for high-quality base calls

8) Interpreting a Sequencing Chromatogram
   - Click on chromatogram data to extract the DNA sequence:

     NNNNNNNNNANNANNNTTTGANTGGATTTTTTTGTTTTGGCAAGAGCATGAGAGCTTTTACT
     GGGCAAGAAGACAAGAGATGGAGAGTCCAGCCGGGCCTGCGCTTAAGTGCGCGGTCTTGCT
     AGGCTTGTAAGTTTCTTTCTTGCTATTCCAAACGGTGAGAGATTTCTGTGCTTTTGTTATAGG
     ACAATTAAAACCGTTTCAATACAACACACTGTGGAGTTTTCATATCTTTGCAACTTTTTCTTT
     GGGCATTCGAGCAATCGGGGCCCAGAGGTAACAAACACAAACAATTTTATCTATTCATTAA
     ATTTTTGTCAAAAACAAGAATTTTCGTAACTGGAAATTTTAAAATATTAAAAACTTTCAACA
     ACGGATCTCTTGGTTCTCGCATCGATGAAGAACGCAGCGAAATGCGATACGTAATGTGAAT
     TGCAGAATTCCGTGAATCATCGAATCTTTGAACGCACATTGCGCCCCTTGGTATTCCAGGGG
     GCATGCCTGTTTGAGCGTCATTTCCTTCTCAAACATTCTGTTTGGTAGTGAGTGATACTCTTT

GGAGTTAACTTGAAATTGCTGGCCTTTTCATTGGATGTTTTTTTTCCAAAGAGAGGTTTCTCT
GCGTGCTTGAGGTATAATGCAAGTACGGTCGTTTTAGGTTTTACCAACTGCGGCTAATCTTT
TTTTATACTGAGCGTATTGGAACGTTATCGATAAGAAGAGAGCGTCTAGGCGAACAATGTTC
TTAAAGTTTGACCTCAAATCAGGTAGGAGTACCCGCTGAACNANGNNNNANNNNNNTTTTNN
NNTNNNNNNAANCAAGAAANNTTTTNNGGGNNAAAAANNAAAAANGGAAAACCACCCGGG
CNGCGCAAANNGGGCGGCCCCGCGNNNGGNAATTTCTTCCGGCTTCCNAACGGGNAGAATT
NCTGGCTTNGTTAAGGACAAAAANCCTTNCCATANCACCNGGGNNNNNCAATCTTNCACCT
TNCTTGGGGNNNNNNNNNGCGGNCCNAGGNAAANNNNNCAATTTNNNNNCCTTNTTTNGTCA
ACAAGAATTCAANCGAANTTAANNNAANNNNNNCAANGTTNGNNNCNGCCTTAAAANNGN
NNNNNNNANNGNANNGAANCCGNNNNNNNTAANCNNGGCCNNNTNNNGGNNNNNNGGNT
TCNTNNCATNNNNCNNNNTNNNNNANNNATTCNNNGNNNNNGANNNNTGGNAANNNNNN
NNNNNNNNNNNNTTANNNNNGTAGAAANNANNCTNNNNNNNNTNNNNNNNNNNNNNNNNN
NNNNANGNNNNNNGNGANGNNNNNNGANCNNNNNNNNNNNNNNNANNNN

9) BLAST
- Basic Local Alignment Search Tool
- Developed by David Lipman and colleagues in 1990 at the National
- Center for Biotechnology Information (NIH NCBI)
- Uses a heuristic (approximating) algorithm to rapidly identify alignments
- Does not necessarily give the best possible alignment, but is extremely fast
- Speed is incredibly important if you are searching extremely large databases
- BLAST is generally robust enough and sensitive enough to use for most purposes

10) Open BLAST
- https://blast.ncbi.nlm.nih.gov
- BLAST is actually several algorithms
- We will use Nucleotide BLAST
- Play around with inputting your DNA sequence

11) Example BLAST Output
- Screenshot image from one of the student's BLAST outputs
- sorted by E value

12) The EXPECT (E) Value
- Used to control score reporting (BLAST will not return matches below the set E value threshold)
- The default threshold for E is 10, which means that 10 matches with scores this high are expected to be found by chance
- Lower E value thresholds are more stringent and will report fewer matches
  - (e.g. E = 1, only one match with a score this high expected to be found by chance)

13) Example BLAST Output
- Screenshot image from one of the student's BLAST outputs
- Accession = Link to GenBank file
- Max Score = Alignment
- Links = Link to Entrez for gene (Entrez is the NCBI search engine)

14) Example BLAST Alignment
- Screenshot image of the 1st alignment

15) Blast Settings: Low Complexity Sequences Filtered
- unclick that option for your sequences
- The ITS region has low complexity regions that actually might help us to differentiate species
- Screenshot showing alignment showing low complexity region changed to a string of unaligned N's

16) For your ITS sequences...
   - Note the parameters you used for the BLAST search (which BLAST database, any parameters that are not default)
   - Note the top hits, E values, and the % Identity (use those to putatively assign a species)
   - If <97% identity, this could be a new species. In this case look in the chromatogram to see if the base calls might be sequencing errors
   - You can print the output as a PDF and email it yourself for printing and pasting in your lab notebook

**Week 5 Lab Lecture Slides**

**Day 2:**

1) Brief Overview of the Malting Process
   - Malting is the partial germination of cereal grains
   - Why do we need to do this to brew?
   - The purpose of malting is to produce enzymes necessary to break down starch into sugars that the yeast can use

2) Brief Overview of the Malting Process
   - Steeping: increasing the water content
   - Germination: allow the barley grain to partially develop
   - Kilning: dry the barley to stop complete germination

3) Brief Overview of the Malting Process
   - Image of steeping from Miller Brewing Company
   - Steeping: increasing the water content
     - soak in water with gibberellins, aerate for ~2 days, and drain.

4) Brief Overview of the Malting Process
   - Image of germinating barley from Miller Brewing Company
   - Germination: grow the barley grains
     - allows for the development of malt enzymes needed to break down starches
     - agitate the grains to aerate and remove generated heat

5) Brief Overview of the Malting Process
   - Image of germinating barley from Miller Brewing Company
   - Kilning: dry the barley to stop germination and allow for storage
     - flow heated air onto a bed of barley to dry

6) Malt Color is Determined by Kilning
   - Images of different colored malted grains and the associated beer hues
   - color traditionally measured in degrees Lovibond

7) Brief Overview of the Brewing Process
   - Milling the grain: need to expose the starch
   - Mashing: converting starch to sugar by activating enzymes in the malted grains (e.g. amylases)
   - Sparging: separating the sugars from the spent grain
   - Boiling: sterilizes the wort and extracts hop oils

8) The entire purpose of malting is to produce amylases that break down starch
   - Image: Fig. 6.6 from https://www.e-education.psu.edu/egee439/node/662

9) Mashing (Converting Starch to Sugars)
   - alpha-amylase: random 1-4 cutter, can ultimately yield glucose and maltose
   - image showing random cuts of …G-G-G-G… starches and their "random" products

10) Mashing (Converting Starch to Sugars)
   - beta-amylase: cleaves from the end to generate maltose
   - image showing random sequential G-G cuts to shorten the starch
   - image of maltose from Wikipedia: https://commons.wikimedia.org/wiki/File:Amylase_reaction.png

11) Mashing (Converting Starch to Sugars)

- Both amylase types have different temperature optima, so the temperature you *mash* at affects the types of sugars and complex carbohydrates in the wort
- alpha amylase: optimum = 68°C – 75°C
- beta amylase: optimum = 54°C – 65°C

12) **Clicker question: All else being equal, mashing at which temperature would result in in a lower alcohol beer with a fuller body?**
    - **A. high temperature (69°C)**
    - B. low temperature (63°C)
    - C. moderate temperature (66°C)

13) Malt Color is Determined by Kilning
    - **Small group discussion: What malt properties may be affected by kilning at a higher temperature?**
        - **denatures amylases (need "pale" malts to compensate), generates Maillard products that are generally unfermented (increases fullness of beer)**

14) Mashing (Converting Starch to Sugars)
    - We're going to mash at an intermediate temperature to generate a medium-bodied base wort
    - Testing carbohydrate content: specific gravity (density)
        - The density of water at 22°C is 0.998 grams per cubic cm
        - When sugar is dissolved into water, the density increases by a small (but measurable) amount
        - Brewers use specific gravity instead of density: mass of a specific volume of wort divided by mass of that same volume of water
        - The specific gravity before fermentation is the original gravity
        - The specific gravity after fermentation is the final gravity
        - These two numbers can be used to estimate final alcohol content

15) Measuring Specific Gravity
    - Traditionally performed with a device called a hydrometer, which floats in the liquid
    - Works by Archimedes principle: a submerged object will experience a force (buoyancy) equal to the weight of the liquid it displaces
    - Read the bottom of the meniscus and write down the specific gravity of the wort
    - We are aiming for an original gravity of 1.050 (50 gravity units (GU)), which is estimated to give us ~5% alcohol if our final gravity is ~1.012 (12 GU)

16) How much grain to add?
    - Decide the batch size (1.5 gallons)
    - Decide the target original gravity (1.050 or 50 GU)
    - Determine total gravity units (TGU) per gallon (50 GU x 1.5 gallons = 75 TGU)
    - Calculate the percentage of each grain (100% Briess 2-Row)
    - Convert the extract potential of each grain into GU (Extract potential is 80% from the manufacturer, meaning 80% of the sugars are expected to be extractable compared to 100% for table sugar)
        - 1 lb of sugar combined with 1 gallon of water = 46 GU
        - So, 80% of that = 36.8 GU
    - Adjust for mash efficiency (expect 70% conversion to fermentable sugar during mashing, not 100%) = 36.8 GU x 70% = 25.8 GU
    - So, 1 lb of our grain yields 25.8 GU per gallon, 75 TGU / 25.8 GU = 2.9 lbs

**Tips on How to Give a Scientific Presentation**

1) Reasons to Give a Scientific Presentation: 1) You are compelled to
    1. For practice (you may be doing this a lot!)
    2. To get a job
    3. To teach people stuff
    4. To become better known in your field

5. To get people to read your stuff
6. To gain experience

2) Basic Principles:
    1. Know your audience (don't overestimate their knowledge)
    2. Take it seriously (fair or not, you are being judged)
    3. Simplify when possible
    4. Tell a story
    5. Know yourself (adopt another's style at your own peril)
    6. Learn from other people's talk
    7. Images over text

3) General Advice:
    1. Humor is okay if that's your style, but be careful with forced or rehearsed jokes
    2. Avoid jargon and technical detail, always stay with the big picture
    3. Summarize often
    4. Don't over-speculate – it's more than okay to admit ignorance
    5. Don't go over the time limit!

4) Ways to be annoying:
    1. Unlabeled graphs or meaningless labels Lots of words, complete sentences, or better yet whole paragraphs
    2. Read the text verbatim
    3. Skip the rationale for the talk
    4. Mumble
    5. Use gratuitous animations
    6. Flash your pointer everywhere
    7. Use excessive jargon and abbreviations

5) Final Tips:
    1. Practice talks by volunteering at every opportunity
    2. Rehearse, but don't let it kill your enthusiasm
    3. Try to visualize your target audience when preparing
    4. Focus on transitions and tricky parts
    5. Have the opening minutes down – 1st 2-3 slides
    6. Exude enthusiasm and confidence

**Week 6 Lab Lecture Slides**

**Day 1:**

1) Yeast Pitching Rates
   - Pitching is the brewers' term for inoculating yeast into the wort
   - Too much yeast (over pitching)
     - fast fermentations
     - low levels of esters
     - lack of mouthfeel
     - yeasty flavors
     - poor viability for re-pitching in subsequent fermentations
   - Too little yeast (under pitching)
     - slow fermentation (or "stuck" fermentation—incomplete)
     - longer brew time (lag)
     - increases chance of contaminants taking over
     - lots of fruity character, but increase in off flavors

2) Yeast Pitching Rates
   - Both over and under pitching can result in...
     - high levels of diacetyl (buttery off flavor)
     - high levels of acetaldehyde (tart, green apple off flavor)
     - low attenuation (low ethanol)

3) Yeast Pitching Rates
   - Generally reported as millions of cells per ml wort per degree Plato
   - Degrees Plato (°P) is the concentration of dissolved solids in wort (mainly sugars) as a percentage of by weight
     - e.g. 10°P = 10 g extract per 100 g wort
     - 1°P is approximately 4 gravity units
       - e.g. 50 GU (1.050 OG) / 4 = 12.5°P

4) Yeast Pitching Rates
   - Common pitching rates:
     - Ale Yeast:
       - Lower range: ~5 million cells/ml, 0.4 million cells/ml/1°P
       - Upper range: ~12 million cells/ml, 1 million cells/ml/1°P
     - Lager Yeast:
       - Lower range: ~12 million cells/ml, 1 million cells/ml/1°P
       - Upper range: ~20 million cells/ml, 1.65 million cells/ml/1°P

5) Yeast Pitching Rates
   - Factors to consider:
     - wort gravity
       - high gravity = more sugar -> requires higher pitching rates
     - yeast flocculation
       - very high clumping may require higher pitching rates
     - beer style
       - Belgians and Hefeweizen's may produce more esters at low pitching rates
     - hop interactions
       - yeast cells can absorb iso-alpha acids and affect IBUs
     - bottle conditioning (carbonating)
       - too little yeast = little re-fermentation, too much = off flavors (or worse)

6) Calculating Yeast Pitching Rates

- Step 1: Measuring viability
    - methylene blue staining
    - Image from Wikimedia commons: https://commons.wikimedia.org/wiki/File:Methylene_Blue_Redox_Reaction.svg
        - the dye is reduced to a colorless form by cellular reductases, but is readily re-oxidized to a blue form by $O_2$
        - viable cells are clear, dead cells stain dark blue

7) Methylene blue staining
    - Image: Figures 6 and 7 from http://braukaiser.com/wiki/index.php/Microscope_use_in_brewing

8) Calculating Yeast Pitching Rates
    - Step 1: Measuring viability with a hemocytometer (see worksheet)

9) For our ales...
    - we are looking at 0.5 million cells / ml / °P
    - our test batches are 100 ml, and our OG was ~1.050 (50 GU)
    - to convert OG to °P: 50 GU / 4 = 12.5°P
    - so, 0.5 million cells x 100 ml x 12.5 = 625 million cells
    - Determine the number of viable cells per ml in your starter culture, and then determine how many ml of cells to add to the wort

**Week 6 Lab Lecture Slides**

**Day 2:**

1) Ale Yeast Traits Important for Brewing
   - Attenuation: fermentation ability
   - Flocculation: cell aggregation
   - Byproduct Production: fruity esters, fusel alcohols, vicinal diketones, sulfur volatiles...

2) Attenuation
   - ability of yeast to metabolize wort sugars
   - apparent attenuation is measured by the drop in specific gravity following fermentation
   - all brewing yeast can metabolize mono- and di-saccharides in the wort (mainly glucose and maltose), but some strains also can metabolize tri-saccharides (mainly maltotriose).
     - Approx % sugars: 20% glucose, 45% maltose, 20% maltotriose

3) Ways to Measure Fermentation Ability...
   - Image: ethanol fermentation from Wikimedia Commons: https://commons.wikimedia.org/wiki/File:Ethanol_fermentation-1.svg

4) Ways to Measure Fermentation Ability...
   - $CO_2$ production
   - ethanol production
   - change in specific gravity (original gravity minus final gravity)

5) **Group Discussion: hypothesis for which will ferment wort better: commercial vs. wild**

**Week 7 Lab Lecture Slides**

**Day 1:**

1) Yeast Primary Fermentation Products
   - Image: Figure 2 from Dzialo et al. 2017. FEMS Microbiology Reviews. 41: S95-S128 (with boxes to shade out all but sugar -> pyruvate -> -> ethanol)

2) Yeast Primary Fermentation Products
   - Image: Figure 2 from Dzialo et al. 2017. FEMS Microbiology Reviews. 41: S95-S128 (now full figure to show some of the secondary fates of the sugar and pyruvate)
   - glycerol leads to a "slick"

3) Yeast Fermentation Byproducts
   - Image: Figure 1 from Dzialo et al. 2017. FEMS Microbiology Reviews. 41: S95-S128

4) Yeast Fermentation Byproducts
   - Esters:
     - formed when an alcohol combines with an organic acid (e.g. ethyl acetate)
     - Image of ethyl acetate from Wikimedia commons: https://commons.wikimedia.org/wiki/File:Essigs%C3%A4ureethylester.svg
     - correlates with yeast strain, wort gravity, yeast growth, and fermentation temperature
       - high gravity (more sugar) worts, rapid yeast growth, higher fermentation temperatures all increase ester production
       - pitching high amounts of yeast (less growth) reduces ester formation
     - typically impart fruity aroma and flavor

5) Esters:
   - Acetate esters:
     - ethyl acetate: fruity, sweet
     - isoamyl acetate: banana (e.g. German wheat beers)
     - phenylethyl acetate: rose, honey, apple
   - Fatty acid esters:
     - ethyl caproate: apple, aniseed
     - ethyl caprylate: apple
     - isoamyl decanoate: tropical fruits

6) Higher (Fusel Alcohols)
   - longer chain length and more complex than ethyl alcohol
   - Image: Figure 4 from Dzialo et al. 2017. FEMS Microbiology Reviews. 41: S95-S128

7) Higher (Fusel Alcohols)
   - longer chain length and more complex than ethyl alcohol
   - provide initial sweetness, but astringent after taste
   - formed by metabolism of amino acids
   - increase with fermentation temperature, wort amino acid levels, and wort gravity
   - different strains produce varying amounts (e.g. Wheat beer strains produce 4-vinyl-guaiacol, which has a clove-like character)
   - Image: 4-vinyl-guaiacol from Wikimedia commons: https://commons.wikimedia.org/wiki/File:2-methoxy-4-vinylphenol.png

8) vicinal di(Ketones)
   - vicinal = two functional groups bonded together
   - two major ones important for brewing are diacetyl and 2,3-pentadione

  - Image of each from Wikimedia commons: https://commons.wikimedia.org/wiki/File:Diacetyl_structure.svg, https://commons.wikimedia.org/wiki/File:Acetylpropionyl.svg
- diacetyl has a very low flavor threshold (easy to taste) and has a buttery flavor
  - usually not a flavor you want a lot of
  - produced non-enzymatically from acetolactate (which the yeast produces)
  - broken down by yeast late in the fermentation
  - higher fermentation temperatures lead to both increased formation and increased breakdown
- 2,3-pentandione has a honey flavor
  - found in some Belgian styles

9) Sulfur volatiles
- hydrogen sulfide: rotten eggs
  - rarely what you want
- dimethyl sulfide: malty/cooked corn taste
  - some in lagers, undesirable for most ales

10) Sensory thresholds
- ammonia: 47 ppm
- acetic acid (vinegar): 1 ppm
- chlorine: 0.3 ppm
- hydrogen sulfide: 0.0005 ppm
- ethyl acetate (fruity -> solvent (at high levels): 33 ppm
- isoamyl acetate (banana): 1.6 ppm
- phenylethyl acetate (roses/honey): 0.2 ppm
- ethyl hexanoate (apple/pear/anise): 0.2 ppm
- ethyl octanoate (apple): 0.2 ppm

**Week 7 Lab Lecture Slides**

**Day 2:**

1) Class Hypothesis: **Commercial strains will ferment better**
   - **Scatter plot of the class data showing fermentation efficiency ($CO_2$/min/$OD_{600}$) of wild vs. commercial strains**
   - Questions:
     - Is this significant?
     - How would you test this? **t-test**

2) t-test to compare two means
   - Test of the null hypothesis that two means are equal
   - Results of two-tailed assuming equal variance: $p = 0.09$
   - What does that mean?

3) Transcriptional Regulation
   - Repression: Transcription factors and regulatory proteins bind to DNA and block transcription
   - Induction: Transcription factors and regulatory proteins bind to DNA and allow transcription

4) Basic anatomy of a eukaryotic promoter
   - Image: Promoter from Wikimedia Commons https://commons.wikimedia.org/wiki/File:TATA_box_mechanism.png
   - promoter = DNA sequence immediately 5' of transcription unit that binds RNA polymerase and controls transcription
   - different DNA sequences in the promoter and upstream elements affect how well RNA polymerase recognizes the promoter
   - This is one major way the cell controls gene expression!

5) Using gene expression to predict high producers of esters and fusel alcohols
   - example: fusel alcohols (which lead to harsh flavors at high levels)
     - Image: Figure 1: Pires et al. 2014. Applied Microbiology and Biotechnology. 98(5): 1937–1949
     - **Small Group Discussion: Which step would you focus on?**
       - **rate limiting**

6) Using gene expression to predict high producers of esters and fusel alcohols
   - example: esters
     - Image: Figure 3: Pires et al. 2014. Applied Microbiology and Biotechnology. 98(5): 1937–1949

7) How to could you use PCR to determine how much of a particular mRNA is present?
   - make complementary (c)DNA from mRNA and then use PCR to quantitate
   - qPCR primers
     - *BAT1* correlates best with higher alcohols
     - *ATF1* and *ATF2* correlate best with acetate esters
     - *RDN18* (18s rRNA): "housekeeping" control gene

8) The Basis of Quantitative PCR
   - Image: Amplification plots from Sigma-Aldrich: https://www.sigmaaldrich.com/technical-documents/articles/biology/troubleshooting.html
   - because PCR results in exponential amplification, you can quantitate how much starting material was present based upon when DNA concentrations reach a certain threshold.
   - 

9) Real-time quantitative (q)PCR

- What's in your PCR tube at cycle number 20?
  - –mixture of primers, nucleotides, template cDNA, Taq polymerase, and PCR product
  - –assuming 1 template molecule, 1,048,576 amplicon molecules ($2^{20}$)
- How about cycle 19?
  - same, except 524,288 amplicons
- How about cycle 18?
  - same, except 262,144 amplicons

10) How to quantitate DNA from PCR
- run a gel
  - Image: Figure 5 from Melo et al. 2006. Genetics and Molecular Research. 5(4): 664-687.
  - challenges include which cycle to pick, less sensitive quantification

- fluorescent dyes
  - e.g. SYBR green
    - fluoresces when bound to double-stranded DNA
    - Image from BioRad SYBR Green SuperMixes: http://www.bio-rad.com/featured/en/sybr-green-for-qpcr.html

11) Example: measuring expression differences for YFG...

- Table:

| Cycle | Commercial Ale Yeast | Wild Yeast |
|---|---|---|
| 20 | 1,000,000 | 4,000,000 |
| 21 | 2,000,000 | 8,000,000 |
| 22 | 4,000,000 | 16,000,000 |

- **Clicker question: Which strain had more mRNA of that gene?**
  A. Ale
  **B. Wild**

12) Example: measuring expression differences for YFG...
- Table:

| Cycle | Commercial Ale Yeast | Wild Yeast |
|---|---|---|
| 20 | 1,000,000 | 4,000,000 |
| 21 | 2,000,000 | 8,000,000 |
| 22 | 4,000,000 | 16,000,000 |

- **Clicker question: How much more?**
  A. 2-fold
  B. 3-fold
  **C. 4-fold**
  D. 6-fold

**Week 8 Lab Lecture Slides (no Day 2 lecture)**

**Day1:**

1) How to could you use PCR to determine how much of a particular mRNA is present?
   - make complementary (c)DNA from mRNA and then use PCR to quantitate
   - qPCR primers
     - *BAT1* correlates best with higher alcohols
     - *ATF1* and *ATF2* correlate best with acetate esters
     - *RDN18* (18s rRNA): "housekeeping" control gene

2) Problem: PCR works with DNA, and we are starting with RNA! Need an enzyme that can convert RNA to DNA

3) Retrovirus life cycle
   - Image: https://www.britannica.com/science/retrovirus

4) Synthesize cDNA using reverse transcriptase
   - Image: drawn cartoon of mRNA (with polyA tail) -> (RT from Moloney Murine Leukemia Virus) cDNA

5) Amplify cDNA using gene specific primers and PCR
   - Image: drawn cartoon of cDNA with forward and reverse primers -> amplified DNA

6) Two flavors of primers
   - anchored oligo dT (NVTTTTTTTT; with V = dGTP, dATP, or dCTP, and N = any base)
   - random primers (NNNNNN)

**Week 9 Lab Lecture Slides (no Day 2, holiday)**

**Day 1:**

1) Separation and detection
   - GC: Gas Chromatography
     - Separation of compound of interest from each other and the matrix
   - MS: Mass spectrometer
     - Detection of analytes of interest
     - Leading to identification or quantification

2) Chromatography
   - The separation of compounds of interest from a complex mixture.
   - Image: Schematic of liquid separation: https://www.tissuegroup.chem.vt.edu/chem-ed/sep/lc/lc.html

3) Bit of vocabulary
   - The **analyte** is the substance to be separated during chromatography. It is also normally what is needed from the mixture.
   - **Analytical chromatography** is used to determine the existence and possibly also the concentration of analyte(s) in a sample.
   - A **bonded phase** is a stationary phase that is covalently bonded to the support particles or to the inside wall of the column tubing.
   - A **chromatogram** is the visual output of the chromatograph. In the case of an optimal separation, different peaks or patterns on the chromatogram correspond to different components of the separated mixture.
   - The **eluate** is the mobile phase leaving the column. This is also called effluent.
   - The **eluent** is the solvent that carries the analyte.
   - The **eluite** is the analyte, the eluted solute.
   - An **immobilized phase** is a stationary phase that is immobilized on the support particles, or on the inner wall of the column tubing.
   - The **mobile phase** is the phase that moves in a definite direction. It may be a liquid (LC and Capillary Electrochromatography (CEC)), a gas (GC), or a supercritical fluid (supercritical-fluid chromatography, SFC). The mobile phase consists of the sample being separated/analyzed and the solvent that moves the sample through the column. In the case of HPLC the mobile phase consists of a non-polar solvent(s) such as hexane in normal phase or a polar solvent such as methanol in reverse phase chromatography and the sample being separated. The mobile phase moves through the chromatography column (the stationary phase) where the sample interacts with the stationary phase and is separated.
   - The **sample** is the matter analyzed in chromatography. It may consist of a single component or it may be a mixture of components. When the sample is treated in the course of an analysis, the phase or the phases containing the analytes of interest is/are referred to as the sample whereas everything out of interest separated from the sample before or in the course of the analysis is referred to as waste.
   - The **solute** refers to the sample components in partition chromatography.
   - The **solvent** refers to any substance capable of solubilizing another substance, and especially the liquid mobile phase in liquid chromatography.
   - The **stationary phase** is the substance fixed in place for the chromatography procedure. Examples include the silica layer in thin layer chromatography
   - The **detector** refers to the instrument used for qualitative and quantitative detection of analytes after separation.

4) Basic Chromatography
   - Separation is achieved by differential affinity of the analytes between the mobile and stationary phases

5) Gas Chromatography
   - Analytes are separated in the gas phase
   - Long skinny columns 20-60M x 0.25mm ID

- Ramped oven temperature
- Hot injector >100C
- Image from Wikimedia Commons: https://commons.wikimedia.org/wiki/File:Gas_chromatograph.png

6) Electron Impact Ionization
   - Molecules are ionized by a stream of electrons
   - + ions are accelerated out of the source
   - Non selective
   - Very reproducible
   - Image from Wikimedia Commons: https://commons.wikimedia.org/wiki/File:Schematic_Diagram_of_Electron_Ionization_Instrumentation.jpg

7) GC/MS typically uses a Quadrupole mass spectrometer
   - Image from NASA: https://attic.gsfc.nasa.gov/huygensgcms/MS_Analyzer_1.htm

8) Results:
   - Example GC trace

9) Results:
   - Comparison to known spectra in the NIST library
     - Spectra for 267,376 chemical compounds

10) Data Quality:
    - Quads give mass data that is +/- 0.5 AMU
      - Not accurate mass
    - Confirmation should be done with known standards
    - MS responsive is somewhat compound specific
      - Quantitation should be done with known standards
      - Particularly for unknown or unexpected compound

11) Sample Prep
    - Samples need to be clean and volatile
      - Liquid extraction
      - Headspace
      - Derivatization
      - SPE
    - Dirty samples= poor data and high maintenance and down time

**Week 10 Lab Lecture Slides**

**Day 1 (Computer Lab):**

1) Brief Overview of the Brewing Process
    1. Milling the grain: need to expose the starch
    2. Mashing: converting starch to sugar by activating enzymes in the malted grains (e.g. amylases)
    3. Sparging: separating the sugars from the spent grain
    4. Boiling: sterilizes the wort and extracts hop oils
    5. Primary Fermentation: yeast converts the simple wort sugars to beer
    6. Secondary Fermentation: slow conversion of the remaining fermentables and byproducts (can be done in the bottle)

2) Diastatic Power (and what it means for recipe design)
    - the ability of a malt to reduce starch to sugar (i.e. amylase activity)
    - measured in degrees Lintner – 100 °Lintner is defined using a specific assay for malt amylase activity – each malt type will have a reported value for diastatic power
    - total "grain bill" of mash should be at least 30 °Lintner to ensure conversion (some styles recommend 70 °Lintner)
    - most base malts are above 100 °Lintner.

3) Types of Brewing Malts (look at stevesbrewshop.com)
    - Base Malts – majority of your "grain bill"
        - provide high levels of enzymes for mashing
        - generally pale to golden in color
    - Specialty Malts (E.g. Crystal/Caramel Malts)
        - typically associated with stouts and porters (darker beers)
        - have no enzymes for mashing (never use as base!)
        - should rarely exceed 10% of grain bill
        - made by:
            1. heating wet grain at mash temperatures (to convert the sugars in the grain)
            2. kilning at high temperature to caramelize the sugars

4) Base Malts:
    - comprise the majority of mash ingredients (aka the grain bill)
    - generally lighter in color, provide enough malt enzymes for starch conversion
    - some major classes: two row barley and six row barley
    - Image: two row and six row barley from Wikimedia commons: https://commons.wikimedia.org/wiki/File:BarleyEars.JPG
        - two row: grains grow on 2/6 rows of the ear
            - barley has thinner husks (thus less tannins), but is low in proteins and enzymes
        - six row: grains grow on 2/6 rows of the ear
            - high in proteins and enzymes, but thick husk and more tannins Types of Brewing Malt 6-row 2-row

5) Malt Color is Determined by Kilning
    - color traditionally measured in degrees Lovibond
    - Malt color wheel: http://randymosher.com/mastering-homebrew

6) Base Malts (lots of varieties)
    - 2-row (1.8°Lovibond) –
        - clean and sweet, mild malty finish
        - good base for many ales
    - 6-row (1.8°L)
        - mild, grainy, more malty finish

    - more protein and enzymes than 2-row
    - also a good base for many ales
- Pilsen (1°L)
    - sweet and clean
    - lightest base malt, allows specialty grains to show more
- Pale (3.5°L)
    - biscuit/nutty, heavier 2-row
    - often used in pale ales
- Maris Otter (3.5°L)
    - biscuit/nutty, fully body
    - often used in pale ales and English style beers (session brews)

7) Crystal Malts
- Carapils, Carafoam, Dextrine Malts
    - 1.5 – 3°Lovibond (lightest)
    - roasted slowly at low temp.
    - little caramel overtones, used to add body and foam retention
- Light Caramel Malts
    - 10 – 30°L (light straw color)
    - light honey-like sweetness and body
- Medium Caramel Malts
    - 40 – 60°L (moderate color)
    - most commonly used crystal malt (porters, stouts, bitters)
    - add sweetness and caramel flavor to finished beer
    - add significant body and foam retention
- Dark Caramel Malts
    - 70 – 90°L (much darker, reddish)
    - high roasting level leads to some bitterness (bitter and sweet)
    - add body and foam retention to darker porters and stouts
- Very Dark Caramel Malts
    - 100 – 220°L (much much darker)
    - bitter, nutty, roasted flavor
    - used sparingly (1/8th to ½ pound per 5 gallons), can easily overwhelm

8) Adjuncts (unmalted grains)
- Wheat (malted)
    - higher protein content than barley
    - produces hazy beers fuller mouthfeel – can be used as base (50 - 70%) of total grain bill
    - add body and foam retention to darker porters and stouts
- corn, rice
    - lighten color and body, results in drier beers
- wheat (unmalted)
    - gives sharper "wheat" character than malted wheat
    - often used in Belgians and wheat beers to provide more complexity
- rye
    - spicy, rye character
- oats
    - enhance body and foam retention of some stouts
    - bready or doughy flavor

9) Specialty Ingredients
- honey
- fruit
- molasses
- syrup

- herbs and spices
- Note: make sure to account for in grain bill!

10) Hop History
- prior to ~800 AD, beer was bittered with a herb and spice mixture called gruit –
  - rosemary, ginger, spruce, juniper, and others
- ~1100's – 1200's: commercial hop cultivation in Germany
- 1400's: hopped beer brewed in England
- 1710: English parliament banned the use of non-hop bittering agents
- strongly-hopped pale ales were developed for export to India
  - origin of IPAs in the late 1700's
- currently ~150 different commercially available hop varieties with different properties

11) *Humulus lupulus*
- perennial plant in the *Cannabaceae* family
- bine (not vine): shoots grow in a helix (versus vines that use "suckers" to attach)

12) Introduction to hops
- Image from Wikimedia Commons: https://commons.wikimedia.org/wiki/File:Hops-1.jpg
- cones of the female Humulus lupulus plant
- resins (alpha and beta acids) and essential oils are present in unique organs called lupulin glands
  - Image hop cross section: https://www.cannonhillbrewing.com.au/hopswest-kracanup-hop-cross-section/

13) Boiling your wort…
- isomerizes the alpha acids in the resins (generating iso-alpha-acids)
  - e.g. humulone (most abundant alpha acid in hops)
  - Image from Wikimedia commons: https://en.m.wikipedia.org/wiki/File:Reaction-degradation-humulone.png
- evaporates the volatile essential oils, which produce different flavors and aromas
  - e.g. grassy, piney, earthy, citrus, spicy (depending on hop variety)

14) Major Types of Hops
- bittering: from the alpha acids
  - added at the beginning of the boil (at least 60 min)
- flavor: from the volatile essential oils
  - added with 20-40 min left in the boil
- aroma: from the volatile essential oils
  - added to the last minutes of the boil to prevent evaporation
  - can also be added after fermentation (dry hopping)
- Many different types of hop flavors: fruity, floral, citrus, grassy, spicy
  - Different hop varieties have different flavor profiles
  - Hop aroma wheels can be found in the Hop Aroma Compendium

15) qPCR Analysis: Quality Control
- SYBR Green does not discriminate between specific and nonspecific PCR products
- Melt curve analysis –
  - assesses the denaturation of double stranded DNA during heating –
  - as the DNA denatures, SYBR Green fluorescence decreases –
  - step up temperature from 55°C to 95°C in 2 second steps
- **Small Group Discussion: What is your expectation of melt curve analysis for a single PCR product? Two PCR products? More?**

16) qPCR Analysis: Quality Control

- Image of two melt curves from PCR Troubleshooting and Optimization: The Essential Guide, Chapter 7, Figure 1.

17) qPCR Analysis: Quality Control •
  - Critical Threshold (Ct): point at which PCR amplification exceeds a set fluorescent threshold
  - Image of Amplification Curve: https://bitesizebio.com/24581/what-is-a-ct-value/

18) qPCR Analysis: ΔCt Method •
  - Critical Threshold (Ct): point at which PCR amplification exceeds a set fluorescent threshold –
    - reported in number of cycles.
  - **Clicker Question: What do lower values mean?**
    - **A. high template concentration**
    - B. low template concentration
  - Use of a control gene allows us to normalize to Ct value of that gene –
    - e.g. RDN18 is 18s RNA and should be constant between cells
  - ΔΔCt = ΔCt(your strain) - ΔCt(median of all strains) –
    - Why the median of all strains? **We want to know which strains have higher or lower expression than the average**
  - ΔΔCt = ΔCt[Ct YFG – Ct reference] - ΔCt(median [Ct YFG – Ct ref])
  - Note: since PCR is exponential ΔΔCt is in log space

**Week 10 Lab Lecture Slides**

**Day 2 (Computer Lab):**

1) Beer Styles
    - Image: Beer Styles http://www.xtbrewing.com/beer-chart-full.jpg
2) History of Beer Styles
    - beer "styles" are simply groups of beers with similar characteristics – there can also be "sub-styles" within larger categories
    - yeast fermentation by-products (e.g. esters) make up a significant part of a style's profile
    - historic brewers did not set out to develop a specific "style"
        - e.g. water properties in Europe
            - soft water of Plzen made ales inconsistent, but the soft water made light (Pilsner) lagers taste crisp
            - water in Burton on Trent has high calcium and sulfate content that accentuates the bitterness of well-hopped ales
            - IPAs originated in England to help preserve beer longer
    - German immigration to the US in mid-1800's shape American brewing (many settled in the Midwest) – American lager: 30% corn or rice (~75% of US beer consumption)

3) Note on Beer Styles
    - The Beer Judge Certification Program (BJCP) is a non-profit organization formed in 1985 to promote beer literacy and appreciation
    - The BJCP provides standards for brewing competitions, including a comprehensive style guide
    - The website for BJCP is www.bjcp.org

4) Beer Style Components

| Style Component | Comments |
|---|---|
| Aroma | Malt aroma, hop aroma, yeast character, diacetyl or ester presence, DMS, etc. |
| Appearance | Color, clarity, head retention |
| Flavor | Sweet & malty, hop character/bitterness, carbonation-bite, acidity, diacetyl presence, fruity, clean, estery, etc. |
| Mouthfeel | Body, carbonation, astringency, etc. |
| Overall Impression | General impression that one should have regarding the style |
| **Vital Statistics** | **Comments** |
| IBUs | Bitterness |
| SRM | Color |
| OG | Original Gravity |
| FG | Final Gravity |
| ABV | Alcohol by Volume |

5) Meilgaard Beer Flavor Wheel
    - developed by Morten Meilgaard in 1979 as part of a joint effort of major brewing groups to enable brewers to:
        i. communicate with each other about beer flavor
        ii. name and define each identifiable flavor note in beer
    - Meilgaard Beer Flavor Wheel: http://www.xtbrewing.com/wheel.png

6) Meilgaard Thresholds
- Primary Flavor Constituents (>2 FU)
  - All Beers
    - Ethanol
    - Hop bittering compounds
    - Carbon dioxide
  - Specialty Beers
    - Hop aroma compounds
    - Caramel and roasted flavor compounds
    - Esters and alcohols (high gravity beers)
  - Defective Beers
    - 2-trans-nonenal (oxidation)
    - Vicinal diketones (diacetyl)
    - Sulfur compounds ($H_2S$, DMS)
    - Acetic or lactic acid (contamination)
    - 3-Methyl-2-butene-1-thiol (skunk/lightstruck)
- Secondary Flavor Constituents (0.5-2 FU)
  - Volatiles
    - Banana esters (e.g., isoamyl acetate)
    - Apple esters (e.g., ethyl hexanoate)
    - Fusel alcohols (e.g., isoamyl alcohol)
    - C6, C8, C10 aliphatic acids
    - Ethyl acetate
    - Butyric and isovaleric acids
    - Phenylacetic acid
  - Nonvolatiles
    - Polyphenols
    - Various acids, sugars, and hop compounds
- Tertiary flavor constituents (0.1-0.5 FU)
  - 2-Penethyl acetate, o-amino acetophenone Isovaleraldehyde, methional, acetoin 4-Ethylguaiacol, g-valerolactone

7) Some “Common” Ale styles
- Witbier (Belgian Ale)
  - very pale to light gold, ABV: 4.5 – 5.5%, IBU: 8 – 20
  - moderate wheat-based malty sweetness, complex spicy notes (light honey, vanilla, citrus/fruity), low hop aroma
  - medium-light to medium body
  - e.g. Allagash White
- Pale American (Blonde) Ale
  - light yellow to deep gold, ABV: 3.8 – 5.5%, IBU: 15 – 28
  - light malt flavor, no to low caramel notes, light to moderate hop flavor
  - medium-light to medium body
  - e.g. New Belgium Somersault, Victory Summer Love
- India Pale Ale
  - medium gold to light amber, ABV: 5.5 – 7.5%, IBU: 40 – 70
  - hoppy and bitter, intense hop aromas (citrus, floral, pine, etc.), medium to low malt flavor (clean)
  - medium-light to medium body
  - e.g. Bell’s Two-Hearted Ale, Founder’s All Day IPA
- Amber Ale
  - amber to copper, ABV: 4.5 – 6.2%, IBU: 25 - 40
  - hoppy, moderate-strength, caramel malty flavor
  - medium to medium-full body
  - e.g. New Belgium Fat Tire, Rogue American Amber

- Irish Red Ale •
  - medium amber to reddish-copper, ABV: 3.8 – 5.0%, IBU: 18 – 28
  - subtle flavors, slightly malty with initial caramel/toffee sweetness
  - medium-light to medium body
  - lightly hopped, malty and slightly sweet, slightly nutty taste, relatively full body
  - e.g. Caffrey's Irish Ale, Samuel Adams Irish Red
- American Brown Ale
  - reddish brown to dark, ABV: 4.3 – 6.2%, IBU: 20 - 30
  - malty but hoppy, with chocolate and caramel flavors, often slightly nutty taste
  - medium to medium-full body
  - e.g. Founders Sumatra Mountain Brown, Bell's Best Brown
- American Porter
  - medium to very dark brown, ABV: 4.8 – 6.5%, IBU: 25 – 50
  - dark malt aroma, lightly burnt with coffee or chocolate notes, variable hop aroma (low to high)
  - medium to medium-full body
  - e.g. Anchor Porter, Boulevard Bully
- Imperial Stout
  - very dark brown to black, ABV: 8 – 12%, IBU: 50 – 90
  - rich roasted malt flavors and medium to high hop bitterness, burnt flavor of coffee or chocolate, can have low to intense esters
  - full to very full
  - e.g. Bell's Expedition Stout

8) GC-MS Analysis
   - Image: Figure 2 of Johnson et al. 2017. Beverages. 3:21.
   - GC separates the compounds and GC + MS is used to assign likely identities (based on a libraries of thousands of known compounds)
   - The area under the peak is proportional to the relative amount of a given compound
   - Thus, the $\log_2$ ratio of peak area versus the median will tell you relative abundance of each compound
   - =log(Your Value / median value, 2)
     - Log(number, base) GC-MS Analysis

**Week 11 Lab Lecture Slides (no Day 2 lectures—checking student recipes and placing brewing order)**

**Day 1: Discuss ideas for brewing recipes with each group, have them re-design if necessary**

1) Brewing Protocol
    1. Post-boil volume is 2.5 gallons
    2. Fermentation volume is 1.0 gallon (remember, we are setting up two fermenters)
    3. Determine the mash conditions for your target original gravity:
        - amount of each grain (based on expected conversion efficiency)
        - mash temperature(s) and time(s)
    4. Determine boil conditions (and hop bittering)
    5. Determine amount of yeast to pitch

2) How much grain to add?
    1. Decide the batch size (2.5 gallons)
    2. Decide the target original gravity (1.050 or 50 GU)
    3. Determine total gravity units (TGU) per gallon (50 GU x 2.5 gallons = 125 TGU)
    4. Calculate the percentage of each grain (e.g. 100% Best Ale Malt)
    5. Convert the extract potential of each grain into GU (Extract potential is 82% from the manufacturer, meaning 82% of the sugars are expected to be extractable compared to 100% for table sugar)
        - 1 lb of sugar combined with 1 gallon of water = 46 GU
        - So, 82% of that = ~38 GU 6)
    6. Adjust for mash efficiency (expect 70% conversion to fermentable sugar during mashing, not 100%) = 38 GU x 70% = 26 GU 7)
    7. So, 1 lb of our grain yields 26 GU per gallon, 125 TGU / 26 GU = 4.8 lbs

3) Calculating Expected IBU = alpha acid units (AAU) x utilization (U) 75 / V (volume of the recipe in gallons)
    - Image: Hop Utilization Table: http://www.valhallabrewing.com/dboard/dbnewsl/t9509d.htm
    - AAU = weight (oz) x % alpha acids
    - U = efficiency of isomerization as a factor of wort gravity and time (Table)
    - 75 / V = conversion factor for English to metric (oz/gallon -> mg/L)

4) Yeast Pitching Rates

| Style | Gravity | Pitch Rate (million cells / ml) |
|---|---|---|
| Ale | < 1.060 (15°P) | 6 |
| Ale | < 1.061 – 1.076 (15 – 19°P) | 12 |
| Ale | > 1.076 (19°P) | 18 |

- lower pitch rate increases ester formation as yeast grow
- higher pitch rates have "cleaner" fermentations
- higher pitch rates necessary for "high gravity" fermentations
- target pitch rate is usually 0.5 – 1 million cells per ml per °P
- E.g. target OG is~1.050 (50 GU) or 12.5°P (50 GU / 4 = 12.5°P)
- so, 0.5 million cells x 3,800 ml x 12.5 = 23.75 billion cells Yeast Pitching Rates

**Week 12 Lab Lecture Slides (no lab lectures)**

**Day 1: Brewing Day**

**Day 2: Brewery Tour**

**Week 13 Lab Lecture Slides (no lab lectures)**

**Day 1: Homebrew Store Tour**

**Week 14 Lab Lecture Slides (no Day 2, Holiday)**

**Day 1: Hop Microbiology**

1) Re-Introduction to hops •
   - cones of the female Humulus lupulus plant •
   - resins (alpha and beta acids) and essential oils are present in unique organs called lupulin glands
   - Boiling your wort isomerizes the alpha acids in the resins (generating iso-alpha-acids)

2) Public Service Announcement •
   - humulone (alpha acid) reacts with UV light to generate a free-radical •
   - free radical reacts with cysteine to form 3-meythlbut-2-ene-1-thiol •
   - 3-meythlbut-2-ene-1-thiol is responsible for “skunk” flavor

3) Measuring Hop Bitterness
   - International Bittering Units (IBUs) •
   - 1 IBU = 1 mg of iso-alpha-acids per liter – measured via absorbance at 275 nm
   - General IBU Guide:
     - American Amber Lager: 30-12 IBU
     - English Bitter: 30-40 IBU
     - India Pale Ale (IPA): 60-80 IBU
     - Double or Imperial IPA: 80-100 IBU
     - Barleywine: 70-100 IBU
     - Stout: 30-50 IBU
     - Scottish Ale: 30-50 IBU
     - Porter: 20-40 IBU

4) There are two major morphological classes of bacteria: gram-positive and gram-negative
   - Image of cell walls from Wikimedia Commons: https://commons.wikimedia.org/wiki/File:Figure_22_02_08f.png
   - Hops are biostatic against gram-positive bacteria once they at IPA concentrations

**Week 15 Lab Lecture Slides**

**Day 1:**

1) Bottle Conditioning
    - Basic Principle: add a "priming" sugar, which residual yeast will ferment
    - Impact on the finished product:
        - adds $CO_2$ (carbonates), which is important for mouthfeel and foam
        - removes oxygen (increases shelf life, flavor stability)
        - can add some additional complexity depending on yeast strain (increased ester formation)
        - yeast can consume any residual diacetyl

2) Carbonation Notes on $CO_2$:
    - dissolved $CO_2$ is reported as "volumes"
        - one volume is 1 L of $CO_2$ at 1 atmosphere of pressure in 1 L of beer

3) Table of $CO_2$ ranges for different styles: https://www.homebrewsupply.com/learn/carbonation-the-final-ingredient.html

4) Carbonation Notes on $CO_2$:
    - dissolved $CO_2$ is reported as "volumes"
        - one volume is 1 L of $CO_2$ at 1 atmosphere of pressure in 1 L of beer
        - if force carbonating, can use a pressure gauge to convert psi to volumes –
    - gas solubility increases at lower temperatures –
    - there is going to be some residual $CO_2$ after our primary fermentation
        - at 20°C, we can expect there to already be 0.850 volumes of $CO_2$ in our primary fermentation

5) Carbonation: The Math Part 1
    - How do you calculate the amount of sugar needed per volume $CO_2$?
        1. Equation for fermentation: 1 mol glucose -> 2 mol ethanol + 2 mol $CO_2$
        2. molecular weight of glucose = 180.16; molecular weight of $CO_2$ = 44.01
        3. So, one 180.16 grams of glucose (1 mole) yields 88.02 grams of $CO_2$ (2 moles)
        4. One mole (44.01g) of $CO_2$ at standard pressure occupies 22.4 liters (ideal gas law)
        5. So, 1.96 g of CO2 occupy 1 L (44.01g / 22.4L), which is produced by 4.01 g glucose
        6. Simplified: add 4.01 g glucose per L per volume

6) Carbonation: The Math Part 2
    - Remember: 4.01 g glucose per L per volume
    - Let's imagine the example of a Belgian Witbier (2.9 volumes CO2):
        1. Let's assume we fermented at 20°C (68°F), so we are starting with 0.85 volumes
        2. 2.9 volumes – 0.85 volumes = 2.05 volumes (the amount we want to add)
        3. Assume bottles are 16 ounces or 0.47 liters
        4. So, 2.05 volumes x 0.47 liters x 4.01g glucose = 3.86 g glucose
        5. Our glucose stock is 50% w/v (or 0.5 g glucose per ml), so 3.86 g = 7.72 ml

7) Please, let's not over carbonate!

**Week 16 Lab Lecture Slides (no lab lectures, final student presentations)**

**Table of Contents**

| Title of the Lab | Page Number |
|---|---|
| Instructor Notes | 1 |
| Wild Yeast Collection Cards and Instructions | 2-6 |
| Wild Yeast Enrichment and Isolation | 7-9 |
| Yeast genomic DNA Extraction | 10 |
| ITS PCR | 11 |
| DNA Electrophoresis of ITS PCR | 12 |
| PCR Cleanup | 13 |
| ITS PCR Sequencing | 14 |
| ITS Sequence Analysis and BLAST (computer lab) | 15 |
| Mashing | 16 |
| Starch Iodide Test | 17 |
| Calculating Original Gravity | 18-19 |
| Measuring Fermentation Rate | 20-21 |
| Yeast Viability and Counting | 22-23 |
| Small Scale Fermentations | 24 |
| Yeast Collections for qPCR and GC-MS | 25 |
| RNA Extraction | 26-28 |
| RNA Electrophoresis | 29 |
| cDNA Synthesis | 30 |
| RNA Hydrolysis and cDNA cleanup | 31-32 |
| qPCR | 33-34 |
| GC-MS | 35 |
| qPCR Analysis (computer lab) | 36 |
| GC-MS Analysis (computer lab) | 37 |
| Brewing Protocol | 38-39 |
| Yeast Viability and Counting | 40-41 |
| Hydrometer – Original Gravity | 42 |
| Hydrometer – Final Gravity | 43 |
| Bottling | 44 |
| ABV and Percent Attenuation | 45 |
| Standard Reference Method for Beer Color | 46 |
| Hop Bitterness | 47-48 |
| References | 49 |

**Notes:**

1) BSL2-level protocols must be used when working with unknown environmental samples. Students must show mastery of BSL1-level techniques before performing BSL2-level activities.

2) Gloves, goggles, and lab coats must be worn for every experiment.

3) To avoid contamination, instructors are recommended to aliquot most reagents in single use aliquots. Enzymes should be dispensed by the instructor to ensure correct volume and avoid contamination.

4) All plates should be stored at 4°C until used. Allow plates to warm up for 1 hr at room temperature before use. Plates with colonies should be stored at 4°C until used. Purified DNA, RNA, and PCR products should be stored at -20°C until used.

5) Yeast require overnight growth in liquid culture at 30°C shaking 270 RPM, while taking 2 days to grow on plates.

6) Students will prepare wort to generate a normalized "beer" media for their groups during the fifth week. This shall be stored in 1 L aliquots until further needed.

INSTRUCTIONS for Wild Yeast Collection Kit

Print pages 1-4
-2 pages per sheet
-two sided with short edge binding
-prefer cardstock if possible

### Wild Yeast Collection Kit

*Where to look for wild yeast*

- Try to go off the beaten path→ **away from areas that are likely to be contaminated by human activities (or spilled beer)**
- Some good placed to look:
  - Under wild fruit or berry bushes or trees – especially where berries have fallen into the soil and are starting to rot.
  - Soil under trees – especially if they are injured and leaking sap into the soil or even the tree bark itself

*How to collect sample*

1. Put on gloves.
2. Open the sterile tube—make sure you don't touch the inside of the lid or tube with your hands
3. Scoop up a little soil into the tube (about to the 5 ml mark).
   **Do NOT fill tube above 10 ml mark!**
   If there are rotten berries, sap or rotting leaves in the soil, you can scoop a little of them up too.
4. Seal tube and store at room temperature until you bring the sample back to lab.
5. **Record as much information about the sample collection location as possible on the back of this sheet.**

*Examples of useful information to record*

- Off rock creek trail
- In Fayetteville, AR. Under oak tree in UA arboretum.
- In Ozark National Forest, near West Cobb
- Soil from under oak tree
- Rotten elderberries were present in soil

### Wild Yeast Collection Kit

*Where to look for wild yeast*

- Try to go off the beaten path→ **away from areas that are likely to be contaminated by human activities (or spilled beer)**
- Some good placed to look:
  - Under wild fruit or berry bushes or trees – especially where berries have fallen into the soil and are starting to rot.
  - Soil under trees – especially if they are injured and leaking sap into the soil or even the tree bark itself

*How to collect sample*

1. Put on gloves.
2. Open the sterile tube—make sure you don't touch the inside of the lid or tube with your hands
3. Scoop up a little soil into the tube (about to the 5 ml mark).
   **Do NOT fill tube above 10 ml mark!**
   If there are rotten berries, sap or rotting leaves in the soil, you can scoop a little of them up too.
4. Seal tube and store at room temperature until you bring the sample back to lab.
5. **Record as much information about the sample collection location as possible on the back of this sheet.**

*Examples of useful information to record*

- Off rock creek trail
- In Fayetteville, AR. Under oak tree in UA arboretum.
- In Ozark National Forest, near West Cobb
- Soil from under oak tree
- Rotten elderberries were present in soil

### SAMPLE INFORMATION

Collected by ____________________________________________

Date collected __________________________________________

Description of location sample was collected from:

____________________________________________________________

____________________________________________________________

____________________________________________________________

____________________________________________________________

____________________________________________________________

____________________________________________________________

____________________________________________________________

____________________________________________________________

____________________________________________________________

____________________________________________________________

### SAMPLE INFORMATION

Collected by ___________________________________________

Date collected _________________________________________

Description of location sample was collected from:

___________________________________________________

___________________________________________________

___________________________________________________

___________________________________________________

___________________________________________________

___________________________________________________

___________________________________________________

___________________________________________________

___________________________________________________

___________________________________________________

**Wild Yeast Enrichment and Isolation** (1)

**Purpose:** Collect wild yeast for identification, further trait characterizing, and finally to compare brewing performance.

**Materials**

**Yeast Enrichment Media 1 (YEM1)**

3 g/L yeast extract (0.3% final)
3 g/L malt extract (0.3% final)
5 g/L bacto peptone (0.5% final)
10 g/L sucrose (1% final)

Bring volume to 915.5 ml with $ddH_2O$

---

autoclave 20 min liquid cycle or filter sterilize

---

once cool, add:

84 ml/L 95% ethanol (8% final)
50 µl of 20 mg/ml chloramphenicol (1 µg/ml final)
415 µl 6N HCl

Check pH using pH paper—should be pH 5-6.

**Yeast Enrichment Media 2 (YEM2)**

5 g/L Yeast Nitrogen Base (YNB) without amino acids (Sigma # Y0626)
2 g/L Yeast Synthetic Drop Out Supplements (YSDO) (see below)
20 g/L alpha-D-methylglucoside

Bring volume to 1000 ml with $ddH_2O$ and transfer to a 2 L flask

20 g bacto agar

---

autoclave 20 min liquid cycle

---

add 1.3 ml 6N HCl

Check pH using pH paper—should be pH 3-4
Pour plates (~25 ml per plate)

Sterile spreaders (or glass beads) (3/student)
Sterile petri plates
P200 tips
Gallon size zip lock bags (1/student)
Quart size zip lock bags (3/student)
Sterile 50ml conical tubes (3/student)
Wild yeast collection card (3/student)
Bunsen burners/ethanol burners
YEM1 (150ml/student)
YEM2 plates (6/student)
Gloves (3 pairs/student)

**Before the lab:**

Must assemble Wild Yeast Collection Packets before first day of class.

- Assemble a sterile 50ml conical tube x1, Wild Yeast Collection Card x1, and 1 pair of gloves and place in a quart size zip lock bag
- Place 3 assembled quart bags inside a gallon sized zip lock
- Repeat for each student

**YNB Components** (µg/L unless indicated)

Nitrogen Sources:
Ammonium sulfate, 5.0 g/L

Vitamins:
Biotin, 2.0
Calcium pantothenate, 400
Folic acid, 2.0
Inositol, 2.0 mg/L
Nicotinic acid, 400
p-Aminobenzoic acid, 200
Pyridoxine HCl, 400
Riboflavin, 200
Thiamine HCL, 400

Trace Elements:
Boric acid, 500
Copper sulfate, 40
Potassium iodide, 100
Ferric chloride, 200
Manganese sulfate, 400
Sodium molybdate, 200
Zinc sulfate, 400

Salts:
Potassium phosphate monobasic, 1.0 g/L
Magnesium sulfate, 0.5 g/L
Sodium chloride, 0.1 g/L
Calcium chloride, 0.1 g/L

***YSDO Components** (mg/L)

Adenine, 18
p-Aminobenzoic acid,8
Leucine, 380
Alanine, 76
Arginine, 76
Asparagine, 76
Aspartic acid, 76
Cysteine, 76
Glutamic acid, 76
Glutamine, 76
Glycine, 76
Histidine, 76
myo-Inositol, 76
Isoleucine, 76
Lysine, 76
Methionine, 76
Phenylalanine, 76
Proline, 76
Serine, 76
Threonine, 76
Tryptophan, 76
Tyrosine, 76
Uracil, 76
Valine, 76
* YSDO lacking a single nutrient can also be ordered from Sigma (e.g. # Y1501 = YSDO without uracil); if you use premixed YSDO make sure to add back the missing nutrient (e.g. 76 mg/L uracil)

**Methods:**

**Day 1 (~10 minutes)**

1) Give students Wild Yeast Collection packets

**Day 2 (~10 minutes)**

1) Add 50 ml YEM1 to sterile 50 ml conical tube containing collection material and tighten cap.
2) Vortex well, tighten caps until just closed, and then loosen by ¼ turn.
3) Place at room temperature in the dark. (may wrap tubes in aluminum foil)

**Day 4 and Day 8 (~15 minutes)**

1) Observe samples for fermentation by looking for the production of bubbles.
    a. If bubbles are absent on Day 4, continue to incubate the plate until Day 8.

2) For samples containing bubbles, spread 50 µl onto YEM2 plates.
3) Incubate at 30°C for 2 days.

**Day 6 and/or Day 10 (~30 minutes)**

1) Add 10 µl sterile water to a glass slide.
2) Stab a putative yeast colony with 200 µl pipet tip and mix with the water on the glass slide.
    a. Yeast colonies are generally smooth and cream colored.
3) Use a microscope to observe whether the colony contained budding cells.
4) For colonies containing budding yeast, streak the colony to a fresh YEM2 plate to isolate single colonies.
5) Incubate at 30°C for 2-3 days.

**Yeast genomic DNA Extraction (~30 minutes)** (2)

**Purpose:** To identify of our yeast species, we must first extract DNA for subsequent PCR and sequencing. DNA extraction and PCR (page B11) can be done in the same lab period.

**Materials:**

LiOAc Lysis Buffer:

1 ml 2 M Lithium Acetate (200 mM final)
1 ml 10% SDS (1% final)
8 ml sterile water
10 ml total (filter sterilize or use filter sterilized reagents)

TE:

10 mM Tris-HCl, pH 8.0,
1 mM EDTA

Sterile 1.7 ml microfuge tube (2/student)
Sterile P200 tips
Bunsen burners/ethanol burners
100% Ethanol
Microcentrifuge
Heat block/thermomixer

**Before the lab:**

- Pipette LiOAc Lysis Buffer, 100% Ethanol, and TE pipetted out into single use aliquots for each student
- Make PCR Master Mix (n+0.5 reactions) and pipette into single use aliquots for each student
- Have heat block/thermomixer incubating at 70°C
- Keep Taq polymerase at -20°C or freezer ice block until ready to use

**Methods:**

1) Add 100 µl LiOAc Lysis buffer to a sterile 1.7 ml microfuge tube.
2) Stab a single colony 8x with sterile P200 tip, add to the 100 µl LiOAc Lysis buffer.
3) Incubate at 70°C for 15 min.
4) Add 300 µl of 100% EtOH and vortex well.
5) Spin at 15000 x g or max speed in microcentrifuge for 3 min.
6) Decant supernatant. **Note: if pellet becomes dislodged, do not pour, use a pipet**
7) Resuspend pellet in 100 µl TE
8) Spin at 15000 x g or max speed in microfuge for 1 min,
9) Carefully pipet supernatant to new sterile 1.7 ml microfuge tube.
10) Use 1 µl per 50 µl PCR.
11) Store samples at -20°C

**ITS PCR for Identification of Yeast Species (~10 minutes)** (3)

**Purpose:** To determine the yeast species, we need to amplify and sequence a region of the DNA that is conserved enough to be found in our species, but has enough variability to differentiate species. The internal transcribed spacer (ITS) regions have been traditionally used to differentiate yeast species.

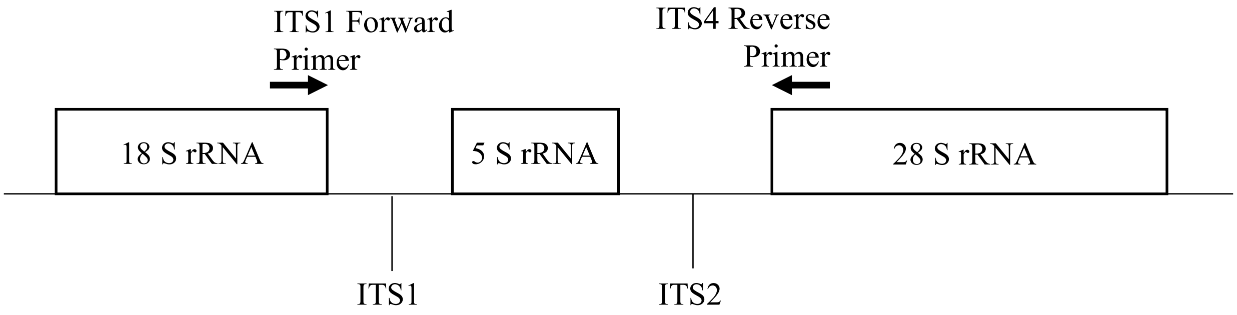

Image made by Amanda Scholes

**Materials:**

PCR Master Mix:

8 µl of 50 µM ITS1 primer (5'-TCCGTAGGTGAACCTGCGG-3'; 0.5 µM final)
8 µl of 50 µM ITS4 primer (5'-TCCTCCGCTTATTGATATGC -3'; 0.5 µM final)
8 µl of 25 mM dNTP mixture (250 µM final)
80 µl Taq Reaction Buffer (10x stock, 1x final concentration, preferably green)
664 µl nuclease-free water
768 µl total

Sterile PCR tubes (1/student)
Sterile P10 tips
Bunsen burners/ethanol burners
RNase-Free filter P10 tips (Instructor only)

**NOTE: Keep Taq polymerase in freezer/freezer block until ready for use.**

**Methods:**

1) Add 96 µl of PCR Master Mix to a PCR tube.
2) Add 2 µl of yeast genomic DNA.
3) Add 2 µl of Taq polymerase (obtain from the TA)
4) Mix well by pipetting gently. **Note: Do not vortex enzymes, they can be mechanically denatured by air bubbles.**

**PCR Cycling Conditions:**

94°C ----> 5 min
94°C ----> 30 sec. \
50°C ----> 30 min x 40 cycles
65°C ----> 2 min. /
65°C ----> 10 min

**DNA Electrophoresis of ITS PCR (~30-55 minutes)**

**Purpose:** To verify we have extracted and PCR the ITS region from a single yeast species (hybrids will show multiple bands). Electrophoresis and, PCR cleanup (page B13), and sequencing (page B14) can be performed in one lab period.

**Materials:**
Gel apparatus with combs
Gel visualizer (e.g. transilluminator)
1X TBE Running buffer
Agarose
Smart-Glow DNA Gel Dye
DNA ladder
*optional loading dye
Sterile P10 tips

**Before the lab:**
- Have 1% agarose gel ready
- Have Binding Buffer (B2), Wash Buffer (W1), and Elution Buffer (E1) pipetted out into single use aliquots.
- Note: If using a new kit make sure to add the isopropanol to the Binding Buffer and add 100% Ethanol to the Wash Buffer.

**Methods:**

1) Pipette gently to mix sample
2) Load 5 µl into well
    a. Be careful agarose gels can tear easily with tips
    b. If samples are clear will need to add loading dye
3) Run gel
    a. 120V for 45 min or 150V for 20 min
4) Visualize gel with transilluminator

**PCR Cleanup (~30 minutes)**

**Purpose:** We need to remove any leftover PCR reagents (dNTPs, primers, salts, enzyme) that may interfere with the sequencing.

**Materials:**
PureLink Quick gel extraction and PCR cleanup combo kit, 50 preps (ThermoFisher #K220001)
Sterile 1.7 ml microfuge tube (2/student)
Sterile P200 tips
Sterile P1000 tips
Bunsen burners/ethanol burners
Microcentrifuge
Nanodrop Spectrophotometer (or standard spectrophotometer with UV-transparent micro-cuvettes)

**Methods:**

1) Transfer PCR product to sterile 1.7 ml microfuge tube
2) Add 360uL B2 (Binding buffer) to tube containing PCR reaction (50–100 µl) and mix well by pipetting.
3) Add sample in Binding Buffer from step 2, to the PureLink® Spin Column.
4) Centrifuge the PureLink® Spin Column at room temperature at 10,000 x g for 1 min.
5) Discard the flow through
6) Replace the PureLink® Spin Column into the Wash Tube.
7) Add 650 µl W1 to the PureLink® Spin Column.
8) Centrifuge the PureLink® Spin Column at room temperature at 10,000 x g for 1 min.
9) Discard the flow-through from the Wash Tube and replace the PureLink® Spin Column into the tube.
10) Centrifuge the PureLink® Spin Column at maximum speed at room temperature for 3 min to remove any residual W1.
11) Discard the Collection Tube.
12) Place the PureLink® Spin Column in a new sterile 1.7 ml microfuge tube
13) Add 15 µl E1 (Elution Buffer) sterile, distilled water to the center of the PureLink® Spin Column.
14) Incubate the PureLink® Spin Column at room temperature for 1 min.
15) Centrifuge the PureLink® Spin Column at maximum speed for 1 min.
16) The elution tube contains the purified PCR product.
17) Discard the PureLink® Spin Column.
18) Quantify the samples with spectrophotometer.
19) DNA can be stored at -20°C for long-term storage.

**ITS Sequencing (Eurofins Genomics) – Instructor Only**

**Purpose:** To determine the species of our budding yeast samples, we will need to sequence the ITS region.

**Materials:**
Nuclease-Free Water
Eurofins Genomics account (other sequencing company)
Nanodrop Spectrometer (or standard spectrophotometer with UV-transparent micro-cuvettes)

**Before the lab:**

- Quantify ITS1 Forward Primer (need 8 µM stock) with spectrophotometer

**Methods:**

1) Prepare 8 µM stock of ITS1 Forward primer.
    a. *Start by making a 1:10 dilution of the stock in the primer collection using Nuclease-Free water. In a Nanodrop spectrophotometer, you can measure the concentration of this dilution using the oligo setting – you will need the primer sequence from the original primer tube*
    b. *Use the primer dilution calculator on IDT's website (https://www.idtdna.com/site/account/login?returnurl=%2FCalc%2FDilution%2F) for dilution calculations – you will need to get the molecular weight of your primer from the original primer tube*
2) Mix the following 12 µl reactions in Eurofins barcoded sequencing tubes:

| | |
|---|---|
| x µl | DNA to be sequenced (320 ng) |
| 1 µl | ITS1 Forward Primer (8 µM)—**Only ONE primer per tube!** |
| y µl | Nuclease-Free water (Final volume 12 µl) |

3) Record barcodes in lab notebook. The sequencing result files will be named only by the barcodes.
4) Make sure lids are pushed all the way down on tubes and place tubes in blue sequencing bags
5) Place blue bag in prepaid UPS shipping envelope.
6) Drop off envelope at the shipping center/UPS store.
    a. If you won't make the 4 pm cutoff, wait to prepare your samples until the next day—do **NOT** store premixed samples in the fridge.

**ITS Sequencing computer lab (~40 minutes)**

**<u>Purpose:</u>** Analyze the ITS sequencing results to determine the species of the budding yeast samples.

**<u>Materials:</u>**
SnapGene Viewer (another sequence viewer)
Computer with internet connection

**<u>Before the lab:</u>**
- Download and rename the sequences to strain names (instead of barcodes)
- Upload those sequences to blackboard (for the students to have access)

**<u>Methods:</u>**

1) Download your sequencing file (.scf) and open in SnapGene Viewer
2) In SnapGene click the box that shows quality score
3) Looking at the chromatogram, copy a region of the sequencing with high quality (>300 bp)
4) Open NCBI Nucleotide Blast
5) Paste sequence, uncheck filter "Low Complexity regions" and click "BLAST"
    a. The run might take a few minutes
6) Students will take a screenshot and note the top several species by lowest E value and high percent identity.
    a. If < 97% sequence identity, could be a new species
    b. Note: For later analyses and brewing, we only moved forward with *Saccharomyces cerevisiae.*

We had six wild *S. cerevisiae* strains total (3 per year). To avoid students designing recipes based upon the identities and known brewing properties of the commercial strains, we randomly assigned them an alias (C1-6). The students were broken up into groups of four to design recipes. 1 pair had a wild yeast and the other had a randomly assigned commercial yeast. These were the strains they used for the remainder of the course.

**Mashing (~90 minutes)**

**Purpose:** Mashing converts the starches into sugars that can be utilized by yeast. The progress of mashing was followed with an iodine test (page B17), and the original gravity of the resulting wort is also measured (page B18). We pool the resulting wort and autoclave to make standardized "beer" media for subsequent yeast characterization.

**Materials:**
20-Quart Stainless Steel Basic Stockpot with lid
The Brew Bag for Kettles (brew in a bag)
Binding clips (3/group)
Digital thermometer
Wort chiller
UltraPure Water (water with minimal ions)
Electric single burner hot plate
Extra-long spoons (>24 inches)
Thick kitchen gloves
1.7 ml microcentrifuge tubes (3/group)

**Before the lab:**

- Measure out 9.5 L (2.5 gallons) ultrapure water and pour into pots (1/group)
- Add brew in a bag to pot, Secure bag with binding clips
- Secure thermometer with binding clips, making sure thermometer isn't touching the bottom
- Get the water up to 67°C (takes approx. 1 hr)
- Have wort chiller connected to sink
- Make 50% glucose (corn sugar – positive control), 50% corn starch (negative control), and 0.025 N iodine solution aliquoted out into 1 mL samples in 1.7 ml microcentrifuge tubes
- Each group will need 7 small weigh boats, P1000 tips and P1000 pipet
- 20 L plastic beaker

**Methods:**

1) Add 3 lb of 2 row grain to brew-in-a-bag in pots
2) Mash for 1 hr at 67°C with the lid on, stirring occasionally (approximately every 10 min)
    a. Pulling 500 µl samples at 0 min, 15 min, 30 min, 45 min, and 60 min (and move onto the starch conversion assay – protocol below)
3) Pull brew in a bag and squeeze excess water
4) Turn off and remove pot from hot plate
5) Add wort chiller to pot and turn on cold water
    a. Will take ~20 minutes to cool down
6) Add ~4 L of wort to a 20 L plastic beaker
    a. Mixing wort form all tables together for homogenous beer media
7) Fill 4-1 L bottles and autoclave to sterilize wort per table
    a. Autoclave on a liquid 20 min cycle
    b. There will be some protein that crashes out (this is normal)

**Starch Iodine Test (~10 minutes)** (4)

**Purpose:** Starch conversion assay allows for visualization of how mashing works.

**Information:** Starch and the triiodide anion ($I_3^-$) interact to form a complex starch helix around the iodide resulting in a blue-black color. In the absence of starch, iodine solutions have a brown color

**Hazard:** Iodine can cause eye, skin, and respiratory tract irritation or an allergic reaction. Wear eye protection and gloves.

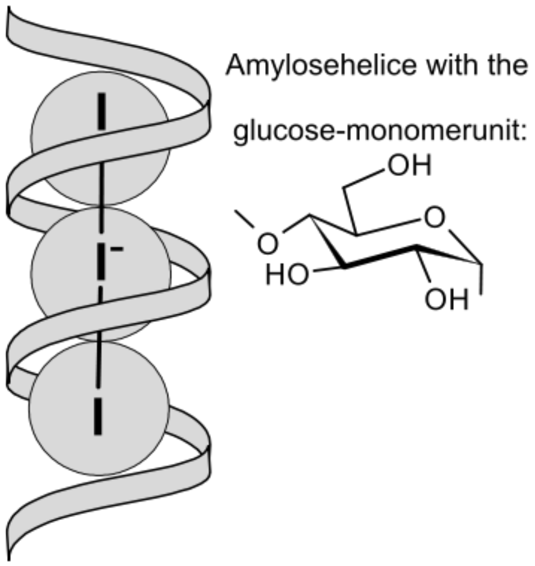

Image taken from Wikipedia

**Materials:**
50% glucose (corn sugar)
50% corn starch
0.025N iodine
Small weigh boats (7/group)
P1000 tips
P1000 pipet

**Methods:**

1) Add 500 µl of 50% glucose into a small weigh boat (enough to fill the bottom).
2) Repeat for cornstarch and every timepoint sample.
3) Add 3-4 drops 0.025 N iodine mix with pipet tip.
4) Record the color

**Calculating Original Gravity (~10 minutes)** (5)

**Purpose:** Hydrometers are used to determine the specific gravity (available carbohydrates) of the wort. This gives information that we will use to calculate the alcohol by volume and percent attenuation at the end of the semester.

**Information:** While not a perfect measure, brewers use specfic gravity (SG) to estimate the carbohydrate content of wort. Carbohydrates are the predominant dissolved solids in the wort, and similarly affect density, so SG is a reasonable approximation. SG is often measured using a hydrometer. The hydrometer is properly read at the bottom of the **meniscus**. Note that the numbers increase going down. In the example below, the hydrometer has a SG reading of 1.047, or 47 points (aka called gravity units or GU).

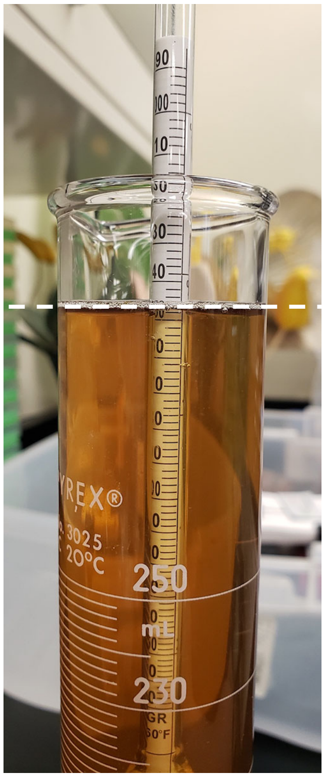

Figure 1: Hydrometer measuring SG of wort.
Image taken by Amanda Scholes

Table 1:

| SG | Percent | SG | Percent | SG | Percent | SG | Percent |
|---|---|---|---|---|---|---|---|
| 1 | 0.25 | 21 | 5.33 | 41 | 10.24 | 61 | 14.98 |
| 2 | 0.51 | 22 | 5.58 | 42 | 10.48 | 62 | 15.21 |
| 3 | 0.77 | 23 | 5.83 | 43 | 10.72 | 63 | 15.45 |
| 4 | 1.03 | 24 | 6.08 | 44 | 10.96 | 64 | 15.68 |
| 5 | 1.28 | 25 | 6.33 | 45 | 11.20 | 65 | 15.91 |
| 6 | 1.54 | 26 | 6.58 | 46 | 11.44 | 66 | 16.14 |
| 7 | 1.80 | 27 | 6.82 | 47 | 11.68 | 67 | 16.37 |
| 8 | 2.05 | 28 | 7.07 | 48 | 11.92 | 68 | 16.60 |
| 9 | 2.31 | 29 | 7.32 | 49 | 12.16 | 69 | 16.83 |
| 10 | 2.56 | 30 | 7.56 | 50 | 12.40 | 70 | 17.06 |
| 11 | 2.81 | 31 | 7.81 | 51 | 12.63 | 71 | 17.29 |
| 12 | 3.07 | 32 | 8.05 | 52 | 12.87 | 72 | 17.51 |
| 13 | 3.32 | 33 | 8.30 | 53 | 13.11 | 73 | 17.74 |
| 14 | 3.57 | 34 | 8.54 | 54 | 13.34 | 74 | 17.97 |
| 15 | 3.82 | 35 | 8.79 | 55 | 13.58 | 75 | 18.20 |
| 16 | 4.07 | 36 | 9.03 | 56 | 13.81 | 76 | 18.43 |
| 17 | 4.33 | 37 | 9.27 | 57 | 14.05 | 77 | 18.65 |
| 18 | 4.58 | 38 | 9.52 | 58 | 14.28 | 78 | 18.88 |
| 19 | 4.83 | 39 | 9.76 | 59 | 14.52 | 79 | 19.10 |
| 20 | 5.08 | 40 | 10.00 | 60 | 14.75 | 80 | 19.33 |

**Temperature Correction**

The hydrometer scale (table 1) above is calibrated to the mass percent of sucrose that has the indicated SG at 20°C. However, water expands as the temperature increases, causing the density to decrease. If the hydrometer is used at a different temperature, a correction must be applied. While it is best to take the reading as close as possible to the temperature for which the hydrometer was designed, the following table gives an approximate correction (in points) that must be added to the reading for 15°C. See table 2, the hydrometer temperature corrections.

Table 2:

| °F | °C | Correction |
|---|---|---|
| 50 | 10 | -1 |
| 59 | 15 | 0 |
| 68 | 20 | +1 |
| 78 | 25 | +2 |
| 86 | 30 | +3 |
| 95 | 35 | +5 |
| 104 | 40 | +7 |

**Materials:**
Bunsen burners/ethanol burners
Hydrometers
Graduated cylinders (250 ml works best for minimizing loss of wort)
Digital thermometers

**Before the lab:**
- Make sure each group has a hydrometer, graduated cylinder, and digital thermometer

**Methods:**

1) Carefully pour wort into graduate cylinder
2) Measure and record the temp of the beer
3) Insert the hydrometer 3/4$^{th}$ way into the filled graduated cylinder
4) Spin and release the hydrometer
5) Read the bottom of the meniscus
6) Make sure to correct for the temperature (according to table 2)
7) Record value in lab notebook

**Calculating Fermentation Rates (~30 minutes)**

**Purpose:** To characterize the fermentation capabilities we determine fermentation rate via $CO_2$ evolution, and set up a small scale fermentation reaction. First, we need to check the viability of our yeast (page B22) before starting the small scale fermentation (page 24). This will give insight into the rate of fermentation, viability, gene expression and ester formation of the wild yeast compared to the commercial strains.

**Information:**
The following equation represents the fate of glucose, energy and reducing equivalents during yeast fermentation:

$C_6H_{12}O_6 + 2ADP + 2Pi + 2NAD^+ \rightarrow 2C_3H_4O_3 + 2ATP + 2NADH \rightarrow 2C_2H_4O + 2CO_2 \rightarrow 2C_2H_5OH + 2NAD^+$
glucose pyruvate acetaldehyde ethanol

This assay measures fermentation efficiency as the volume of $CO_2$ released per minute per cell mass.

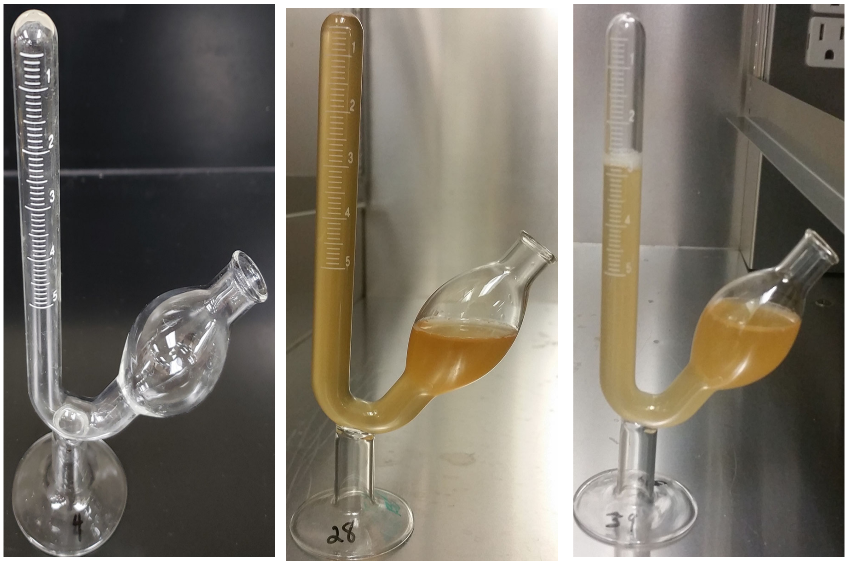

Figure 2: The first panel shows an empty fermentation tube. Second panel shows fermentation tube once filled and about to start the fermentation test. Panel 3 fermentation tube in the middle of the fermentation test. Images taken by Amanda Scholes

**Materials:**
Vortex
Spectrometer
Cuvettes (1/group, Instructor will need one for a media blank)
Wort (made previously)
Fermentation Tube Graduated 5 ml (Science Lab Supplies B2800-2) (2/group)
10 ml glass culture tubes (1/culture)
125 ml Erlenmeyer flask (1/culture)
50 ml conical tubes (1/culture)
Table top centrifuge capable of spinning 50 ml conical tubes
10 ml serological pipets
Pipet bulb/electric pipettor

**Before the lab:**

- **Night before:** Grow a 10 ml overnight culture in wort of each sample
- **Morning of:** Dump 10 ml culture into 30 ml of fresh pre-warmed wort in 125 ml Erlenmeyer flask
- Label tubes and spin down cells 1500 x g for 1 min (students will resuspend in pre-warmed wort)
- Aliquot 10 ml sterile pre-warmed wort/group (keep at 30°C until needed)
- Turn on spectrometer (some take 15 minutes to warm up)

**Methods:**

1) Decant supernatant.
2) Resuspend cell pellet with 10 ml sterile wort via vortexing.
3) Carefully transfer half the cells into fermentation tube
4) Plug opening and invert tube (getting cells into the graduated chamber)
5) Place tube upright and add remaining cells
    a. **Fermentation tubes are VERY fragile. Be careful when handling.**
6) Monitor and record every 2 minutes once it passes the 0.5 ml mark.
7) Make 1:25 dilution of cells into wort (40 µl cells into 960 µl wort) and make blank (1 ml wort)
    a. May need to vary the dilution to get within linear range of spectrometer
8) Spec cells to determine optical density (OD).
9) Plot ml $CO_2$ displaced vs time in minutes.
10) To determine fermentation rate divide slope by the OD.

**Yeast Viability and Counting (~ 20 minutes)**

**Purpose:** To determine the number is live cells to be able to pitch the correct amount of yeast for our small scale fermentation experiment.

**Information:** The dye **methylene blue** can be used to measure yeast viability. Methylene blue is spontaneously oxidized in the presence of oxygen, resulting in a blue color. Cellular reductases catalyze the conversion of methylene blue to the Leuko form, which is colorless (see below). Thus, live yeast cells that contain active reductases will be colorless, while dead cells lacking the enzymes will remain blue.

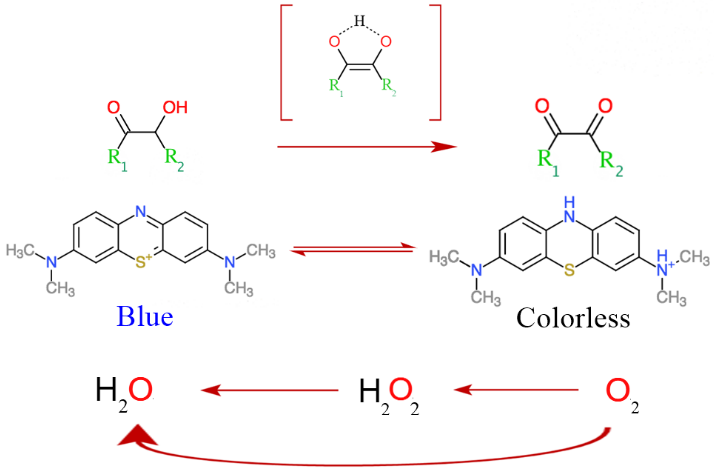

Figure 3: schematic of how the color changing mechanism works. Image from Wikipedia.

The number of viable cells can then be quantified using a cell counting chamber (often referred to as a **hemocytometer**).

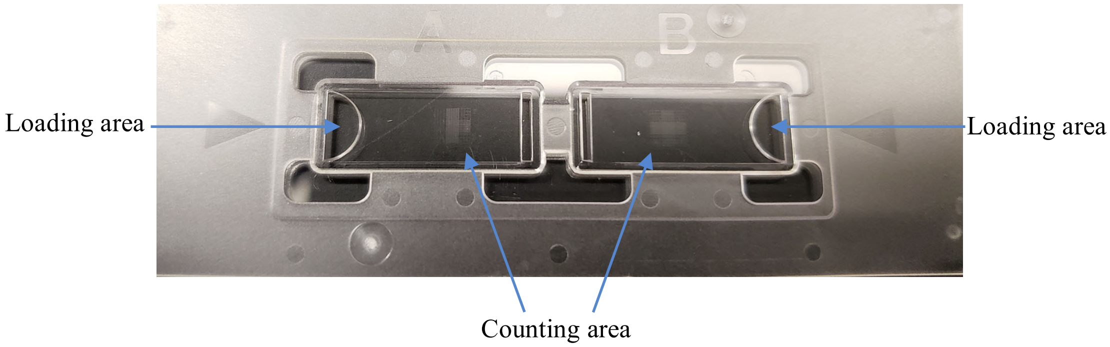

Figure 4: Hemocytometer slide. Image taken by Amanda Scholes

**Materials:**
Hemocytometers
Microscopes
Sterile 50 ml conical tubes
1.7 ml microcentrifuge tubes
Sterile 10 ml glass tubes
Sterile Erlenmeyer flasks (125 ml)
Sterile wort
0.01% Methylene Blue
Optional: Hand tally counter (VWR 23609-102)
**Before the lab:**

- Set up 10 ml cultures (in 10 ml tubes) and grow at 30°C for 1 day
- Next morning set up 30 ml cultures (in 125 ml flasks) and grow at 30°C
- Centrifuge cells in 50 ml conical at 1,500 x g for 3 min
- Carefully, decant media
- Have 20 ml Sterile wort (students made) in 50 ml conical tubes
- Have 0.01% Methylene blue in single use aliquots of 990 µl

**Methods:**

1) Add 10 µl of cells into 990 µl of methylene blue solution (0.01% (w/v)). This constitutes a 1:100 dilution.

2) Vortex briefly and pipette 10 µl of sample into the counting chamber.

3) Place the hemocytometer under a microscope at 10x magnification and find the grid pattern (see below)

4) Count cells using the four corner squares. Count both the total number of cells and the number of live (colorless) cells. For cells falling on a boundary line, only count them if they are on the top or right line (see below). Count budding cells as separate cells if the bud is at least half the size of the mother cell. **Note: try to be consistent in your counting scheme.**

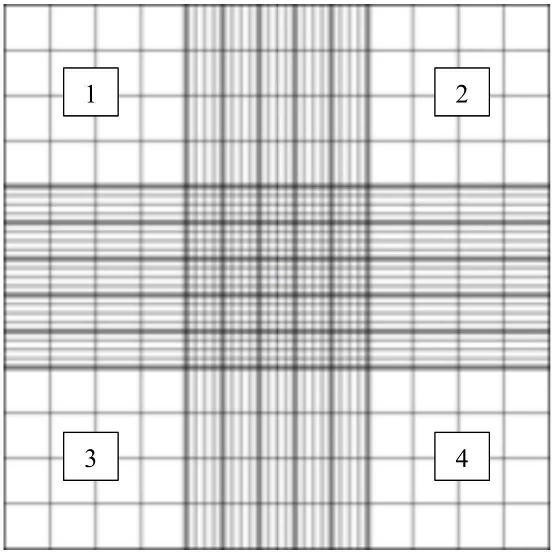

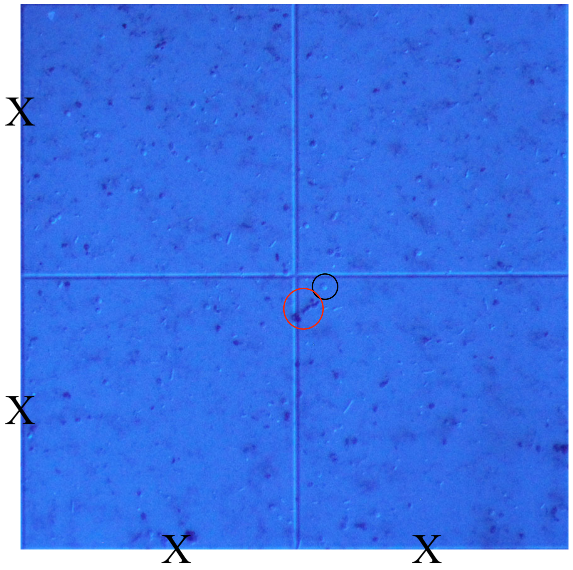

The black circle is showing a live cell (colorless), while the red circle shows dead cells (dark blue).

5) Each of the nine large squares has a volume of 0.1 $mm^3$ ([1 mm x 1 mm] x 0.1 mm chamber depth). There are 1000 $mm^3$ per ml, so each square has a volume of 0.0001 ml or $10^{-4}$ ml.

So, cells per ml = average # of viable cells (per 4 squares) x $10^4$ $ml^{-1}$ x 100 (dilution factor).

**Small Scale Fermentation (~20 minutes)**

**<u>Purpose:</u>** To determine fermentation gene expression and ester formation of these yeast for future recipe development.

**<u>Materials:</u>**
Sterile 125 ml Erlenmeyer flasks (1/group)
Sterile wort (from Mashing protocol pg16)
No. 6 Stopper – Drilled (Northern Brewer Homebrew Supply SKU: SD6) (1/group)
Three-Piece Airlock (Northern Brewer Homebrew Supply SKU: 7010) (1/group)
95% ethanol

**<u>Before the lab:</u>**
- Transfer 100 ml sterile wort into each sterile 125 ml flask
- Autoclave flask stoppers
- Have airlocks soaking in 95% ethanol

**<u>Methods:</u>**

1) Add 625 million cells to flask containing 100 ml sterile wort
2) Add sterile flask stopper onto airlock and carefully plug 125 ml flask
3) Fill airlock with 95% ethanol to fill line
4) After 2 days proceed to Yeast Collections for qPCR and GC-MS (protocol below)

**Yeast Collections for qPCR and GC-MS (~10 minutes)**

**Purpose:** Collect cells to determine fermentation gene expression and ester formation of these yeast for future recipe development.

**Materials:**
50 ml conical tubes (2/group)
Liquid Nitrogen
-80°C freezer

**Before the lab:**

- Give each group 2 sterile 50 ml conical tubes

**Methods:**

1) Label (lid and side) 2-50 ml conical tubes with sample name and:
    a. Tube 1 GC-MS
    b. Tube 2 RNA
2) Carefully **(minimally disturbing the cells)** remove airlock and decant 20 ml of supernatant into 50 ml conical tube labeled GC-MS
3) Swirl flask and decant cells into 50 ml conical tube labeled RNA.
4) Spin down cells at 1500 x g for 3 min
5) Decant supernatant
6) Flash free cells in liquid nitrogen
7) Store cells at -80°C freezer, until ready to be used

**RNA Extraction (~50 minutes)**
modified Zymo Research Protocol

**Purpose:** Extracting RNA to perform qPCR to analyze gene expression of fermentation related genes.

**Materials:**
Quick-RNA MiniPrep Plus kit (Zymo Research R1057)
RNase-Free 1.7 ml microcentrifuge tubes
RNase-Free 2.0 ml microcentrifuge tubes
RNase-Free Filter P100 tips
RNase-Free Filter P1000 tips
RNase-Free PCR tubes
100% RNase-Free Ethanol
Bunsen burners/ethanol burners
Vortex
Multi-Therm incubating vortexer
Microcentrifuge
Nanodrop/Qubit Spectrometer
Ice/ice bin

**Before the lab:**
- Resuspend the students' cell pellet in 400µL with 1X DNA/RNA Shield
- Give students two sets of RNase-free 1.7 ml microcentrifuge tubes, one set of RNase-Free 2.0 ml microcentrifuge tube, and one set of Spin Away Filters (Yellow columns), and Zymo spin columns (Green Zymo Spin III CG columns) for each sample.
- Make D1 (5 µl DNase I and 75 µl DNA Digestion Buffer/sample)
- Aliquot glass beads, RNA Lysis Buffer (L1), 100% EtOH, RNA Wash Buffer (W1), DNase (D1), RNA Prep Buffer (P), RNA Wash Buffer (W2), RNA Wash Buffer (W3), and Elution Buffer (E1) in single use aliquots.
- Turn on and set Multi-Therm to 4°C

**Note: All steps following cell disruption should be performed at room temperature. Work quickly to limit RNA degradation.**

**Note: Do not touch the column resin with tips.**

**Methods:**

1. Resuspend cell pellet in 400 µl with 1X DNA/RNA Shield – Instructor does.
2. Label (lid and side) **two sets** of RNase-free 1.7 ml microcentrifuge tubes, one set of 2.0 ml microcentrifuge tube, one set of Spin Away Filters (Yellow column), and Zymo spin columns (Green Zymo Spin III CG column).
3. Add 400 µl cells (in 1X DNA/RNA Shield) to 400 µl glass beads, and homogenize by placing in MultiTherm set to 4°C and maximum shaking for 10 min.
    a. Vortex samples every two minutes.

4. Add 400 µl L1 (RNA Lysis Buffer) and gently mix by pipetting.
5. Carefully pipette sample into a Yellow Spin Away Filter column (column and collection tube).
6. Centrifuge for 30 sec at 16,000 x g. **Transfer flow through by pipetting to a new RNase-Free 2.0 ml tube**
7. Add 800 µl EtOH (100% EtOH) and gently mix by pipetting.
8. Carefully pipette sample to a Green Zymo Spin III CG mini spin column.
9. Centrifuge for 30 sec at 16,000 x g.
10. Discard flow through by pipetting.
11. Add 400 µl W1 (RNA Wash Buffer) to the column.
12. Centrifuge for 30 sec at 16,000 x g.
13. Discard flow through by pipetting.
14. Add 80 µl D1 (DNase I) directly to the column.
15. Incubate the column at room temperature for **25 min** to digest genomic DNA.
16. Add 400 µl P1 (RNA Prep Buffer) to the column.
17. Centrifuge for 30 sec at 16,000 x g.
18. Discard flow through by pipetting.
19. Add 700 µl W2 (RNA Wash Buffer).
20. Centrifuge for 30 sec at 16,000 x g.
21. Discard flow through by pipetting.
22. Add 400 µl W3 (RNA Wash Buffer).
23. Centrifuge for 2 min at 16,000 x g.
24. Transfer column to a new RNase-Free 1.7 ml microcentrifuge tube **(DO NOT let touch the column the flow through**).
25. Add 50 µl E1 (RNase-free water) and incubate for 1 min at room temp.
    a. **Note: Make sure to pipette water directly onto column resin WITHOUT touching pipette tip to membrane. If water remains on the wall of the column, tap the tube until water absorbs into the membrane.**
26. Centrifuge for 30 sec at 16,000 x g.

27. Carefully pipette the eluted RNA and place it back onto column, incubate for an additional 1 min.
    a. **Note: the second elution increases RNA yield. Note: Make sure to pipette water directly onto column resin WITHOUT touching pipette tip to membrane. If water remains on the wall of the column, tap the tube until water absorbs into the membrane.**
28. Centrifuge for 1 min at 16,000 x g.
29. Transfer Eluted RNA to a fresh RNase-Free 1.7 ml microcentrifuge tube just in case any RNases contaminated the lid of the tube during centrifugation. **Immediately place RNA on ice.**
    a. Long term storage is at -80°C, and RNA concentrations are calculated using a Nanodrop spectrophotometer.
30. Transfer 2 µl of RNA into RNase-Free PCR tube (to be used to run an RNA electrophoresis).

**RNA Electrophoresis (~60 minutes)**

**Purpose:** To verify we have extracted RNA is intact and can be used to make cDNA.

**Materials**
RNase-away
Gel apparatus with combs (size depends on number of samples)
Gel visualizer
RNase-Free 1X TBE Running buffer
Agarose
Erlenmeyer flask (size depends on size of gel – 125 ml for small gel, 250 ml for large gel)
Smart-Glow DNA Gel Dye
DNA ladder
Loading dye
Parafilm
RNase-Free filter P10 tips

**Before the lab:**

- Spray gel apparatus and combs with RNase-away spray and rinse with RNase-Free water
- Make RNase-Free 1% agarose gel

**Methods:**

1) Add 1 µl of loading dye onto a strip of parafilm (1 spot/sample)

2) Pipette 2 µl gently to mix sample

3) Load dyed RNA into well

    a. Be careful agarose gels can tear easily with tips

4) Run gel at 120V for 45 min

5) Visualize gel

**cDNA Synthesis (~25 minutes)**

**Purpose:** Make cDNA from the extracted RNA to be used for qPCR to analyze gene expression of fermentation related genes.

**Materials:**
RNase-Free Filter p10 tips
Anchored oligo-dT (6 µg/µl)
Random hexamer (6 µg/µl)
0.1 M DTT
5X Superscript buffer
10X RNase-Free dNTPs (2.5 mM stock solution, final concentration of 250 µM)
Superscript III Reverse Transcriptase
Bunsen burners/ethanol burners
RNase-Free PCR tubes (1/group)
Thermocycler
Ice/ice bin

**Before the lab:**

- Have anchored oligo-dT, random hexamer, 5X Superscript Buffer, DTT, RNase-Free dNTPs aliquoted out for each table.
- Keep Superscript III RT at -20°C or freezer ice block until ready to use.
- Have thermocycler incubating at 70°C

**Methods:**

1) Prepare RNA/primer mixture in RNase-Free PCR tube:
    - 6 µl total RNA
    - 0.5 µl: anchored oligo-dT
    - 0.5 µl: random hexamer
2) Denature the primer/RNA mixture at 70°C for 10 min in a thermocycler.
3) Immediately chill on ice for >2 min.
4) Prepare reverse transcriptase cocktail in a **separate** RNase-Free PCR tube:
    - 3.75 µl 5x Superscript Buffer
    - 1.88 µl 0.1M DTT
    - 3.75 µl 10x RNase-free dNTPs
5) Add 7.5 µl of reaction cocktail to denatured RNA.
6) Take sample to Instructor to add 0.5 µl Superscript III RT.
7) Incubate on bench (25°C) for 7 min.
    a. This step is critical for allowing random hexamer to anneal to the RNA.
8) Incubate at 50°C for 2 hours in thermocycler.
9) Store samples at -20°C until next class period.

**RNA Hydrolysis and cDNA Cleanup (~25 minutes)**

**Purpose:** Make cDNA from the extracted RNA to be used for qPCR to analyze gene expression of fermentation related genes.

**Materials:**
RNase-Free filter p10 tips
DNA clean and concentrate kit (Zymo Research #D4014)
Bunsen burners/ethanol burners
RNase-Free 1.7 ml microcentrifuge tubes
Thermocycler
Nanodrop/Qubit Spectrometer
Ice/ice bin

**Before the lab:**

- Have 1 M NaOH, 0.5 M EDTA, DNA Binding Buffer (B1), DNA Wash Buffer (W1), DNA Wash Buffer (W2), and Elution Buffer (E1) aliquoted into single use aliquots.
- Give students one RNase-free 1.7 ml microcentrifuge tube and Zymo spin column.
- Have thermocycler incubating at 65°C

**Methods:**

1) Label (lid and side) of both of RNase-free 1.7 ml microcentrifuge tube and Zymo spin column.
2) Add 10 µl 1M NaOH to the cDNA synthesis reaction (PCR tube), mix by pipetting.
3) Add 10 µl 0.5M EDTA to the cDNA synthesis reaction (PCR tube) mix by pipetting.
4) Incubate at 65°C for 15 min.
5) Add cDNA (34.5 µl) sample to a new 1.7 ml tube containing 490 µl B1 (DNA Binding Buffer). Mix thoroughly by pipetting.
6) Add entire sample to Zymo-Spin column.
7) Centrifuge for 30 sec as 16,000 x g.
8) Discard flow through.
9) Add 200 µl W1 (DNA Wash Buffer).
10) Centrifuge at 16,000 x g for 30 sec.
11) Repeat wash with 200 µl W2 (DNA Wash Buffer).
12) Centrifuge at 16,000 x g for 30 sec.
13) Place the Zymo-Spin column in a clean 1.7 ml tube.
14) Add 6 µl E1 (Elution Buffer).

**Note: Make sure to pipette water directly onto column resin WITHOUT touching pipette tip to membrane. If water remains on the wall of the column, tap the tube until water absorbs into the membrane.**

15) Incubate at room temperature for 1 min.

16) Centrifuge at 16,000 x g for 30 sec to elute the cDNA.

17) cDNA concentrations are calculated using a Nanodrop spectrophotometer. Store samples at -20°C.

**qPCR (~20 minutes)**

**Purpose:** To examine fermentation related] gene [*ATF1* (Alcohol acetyltransferase) responsible for the majority of volatile acetate ester, *ATF2* (alcohol acetyltransferase) also forms volatile esters during fermentation. *BAT1* (branched-chain amino acid transferase) makes a precursor that is used to make higher alcohols, *RND18* (18S rRNA) is a housekeeping control gene for normalization.

**Materials:**
Nuclease-Free water
Maxima SYBR Green Supermix (Fisher #FERK0241)
SYBR Green (Fisher #S7563)
RNase-Free filter P10 tips
RNase-Free filterP200 tips
Sterile PCR tube
Sterile 1.7 ml microcentrifuge tube
qPCR plates (Bio-Rad # MLL-9601)
qPCR membranes (Bio-Rad # MSB-1001)
Plate microcentrifuge
Real-Time PCR Machine
Biosafety cabinet
Ice/ice bin

**Before the lab:**

- Have primer master mixes made and stored on ice (n+1 reactions)
- Aliquot Nuclease-Free Water 1 (NFW1) and Nuclease-Free Water 2 (NFW2) into single use aliquots for each student
- Give each student a new PCR tube and 1.7 ml microcentrifuge tube
- Thaw Maxima SYBR Green Supermix and store on top of ice
- Note: do not use the edges of the plate

**Methods:**

1) Label (Lid and side) PCR and 1.7 ml microcentrifuge tube.

2) Dilute the cDNA to 5 ng/µl in 10 µl NFW1 (Nuclease-Free Water) in a new PCR tube.

   **Note:** If your cDNA concentration is <5 ng/µL, just add 5µL NFW1 to your cDNA.

3) Prepare PCR mastermix (this is enough for 4 primer pairs) in a sterile 1.7 ml microcentrifuge tube.

   **Per 4 Reactions:**
   9 µl of diluted cDNA (10 ng final)
   27 µl NFW2 (Nuclease-Free water)
   45 µl Maxima SYBR Green Supermix (see Instructor for this)

4) Place PCR mastermix on ice

**The next steps are exceptionally sensitive to pipetting error and aerosolized DNA, so the Instructor will perform them in a Biosafety cabinet:**

5) Dilute the primers with Nuclease-Free water to a working stock concentration of 10 µM.

6) Prepare primer master mix:

**<u>Per Reaction:</u>**
0.4 µl Forward Primer (200 nM final)
0.4 µl Reverse Primer (200 nM final)
1.2 µl Nuclease-Free Water

The following are the primer pairs:

RDN18-F: CGGCTACCACATCCAAGGAA
RDN18-R: GCTGGAATTACCGCGGCT

ATF1 F: GTTTGTTGGTTTGTATCTCTTCATAAA
ATF1 R: GGGGTGAAGTCAATAAATCCA

ATF2 F: TTCTCGCAGTCTGCAGGC
ATF2 R: GCAGAACGATTCCCATTCG

BAT1 F: CTCAAGAATGGGACATCAACGA
BAT1 R: CTTGGTCAATGCACCACATTG

7) Add 2 µl of the primer mix to the ***<u>bottom</u>*** of the appropriate wells of a qPCR plate.

8) Add 18 µl of the PCR mix to the ***<u>side</u>*** of the appropriate wells. <u>Do not change tips</u> between samples that use the same cDNA.

    a. Careful to only touch the side of the well so as to not contaminate the pipet tip with primer mix.

    b. Change tips for non-identical samples.

9) Seal PCR plate with clear sealing membrane.

10) Briefly spin the plate in a microplate centrifuge.

11) Reaction conditions assume an optimum primer annealing temperature of ~58°C (6)

95°C -> 10 min
-------------------
95°C -> 15 sec
58°C -> 60 sec   40 cycles
-------------------
melt curve analysis (58 – 95°C in 2 sec steps)

**GC-MS**

**Purpose:** Gas chromatography-mass spectrometry (GC-MS) allows us to determine what esters are being produced by yeast during fermentation.

**Materials:**
Shimadzu 2020 GC/MS single quadrupole system
20 mm headspace vials with silicon PTFE septa (DWK Life Sciences Millville, NJ, USA)
10 ml serological pipet
Pipet bulb/electric pipettor

**Methods:**

1) Transfer 10 ml of supernatant into 20 mm headspace vials with silicon PTFE septa using a sterile serological pipet.
2) 1000 µl of headspace was injected in the GC equipped with a RtX-VMS 30M 0.32ID 1.8 µM column (Restek Bellefonte, PA, US).
3) GC-MS machine operation information
    a. Samples were analyzed using a Shimadzu 2020 GC/MS single quadrupole system.
    b. The injector was set to 120°C, initial oven temperature of 35°C rising at 10°C/min to 230°C hold 5 mins in constant velocity mode.
    c. The MS was operated in full scan mode.
4) Peak identification was done with spectral matching to the NIST 2014 database.

**qPCR Computer Lab**

**Purpose:** To examine fermentation related] gene [*ATF1* (Alcohol acetyltransferase) responsible for the majority of volatile acetate ester, *AFT2* (alcohol acetyltransferase) also forms volatile esters during fermentation. *BAT1* (branched-chain amino acid transferase) makes a precursor that is used to make higher alcohols, RND18 (18 S rRNA) is a control gene] expression of the yeast.

**Information:** Critical Threshold (Ct): point at which PCR amplification exceeds a set fluorescent threshold (the number of cycles gives us this information). Lower Ct means it took less cycles to reach that threshold (higher gene expression). RDN18 (18S rRNA) gene is our control gene, which we will normalize our genes to.

**Methods:**

1) For each strain, calculate: $\Delta Ct_1$ [Ct(test gene e.g. *ATF1*) – Ct(*RDN18*)]

2) Then, calculate the median Ct value for the test gene for all strains (call this $\Delta Ct_2$)

3) $\Delta\Delta Ct = \Delta Ct_1$ [your strain] - $\Delta Ct_2$ [median of all strains]

4) Note: since PCR is exponential $\Delta\Delta Ct$ is in log space: to convert to fold change use the formula $2^{-\Delta\Delta Ct}$

**GC-MS Computer Lab**

**Purpose:** GC-MS allows us to determine what esters are being produced by yeast during fermentation. This information will be used to tailor the beer recipes to emphasize certain esters.

**Information:** Similar concept of comparing your ester production to the median of all the samples. The area under the peak is proportional to the relative amount of a given compound. Thus, the log2 ratio of peak area versus the median will tell you relative abundance of each compound

**Methods:**

1) $Log_2$ transform all of the peak areas for each strain and compound.
2) Calculate the median $log_2$ value for the esters of interest from all the strains.
3) To calculate the relative amount of ester for your strain, subtract (Your Ester of Interest) – Median (Ester of Interest)
4) To convert to fold change use the formula $2^{\text{(Your Ester of Interest) – Median (Ester of Interest)}}$

**Brewing Protocol (~3 hrs)**

**Purpose:** We will go through the entire brewing protocol, from mashing to boiling (with hop additions), to chilling and pitching yeast. Each group will prepare enough wort to fill two fermenters—one that will be pitched with commercial yeast and one that will be pitched with wild yeast. During the boil, students can measure yeast viability of their starter cultures to determine how much to pitch (page B40). Once the wort has been prepared and chilled, we will measure the original gravity (page 42).

**Materials:**
20-Quart Stainless Steel Basic Stockpot with lid
The Brew Bag for Kettles (brew in a bag)
Binding clips (3/group)
Digital thermometer
Wort chiller
UltraPure Water (water with minimal ions)
Electric single burner hot plate
Extra-long spoons (>24 inches)
Thick kitchen gloves
Hop bags
Hops
Three-Piece Airlock (Northern Brewer Homebrew Supply SKU: 7010)
95% ethanol
Small Batch 1 Gallon Fermenter (Northern Brewer Homebrew Supply SKU: 41097)
Hand operated auto siphon with tube
Star San

**Before the lab:**

- Have wort chiller connected to sink
- Measure out 13.25 L (3.5 gallons) ultrapure water and pour into pots
- Add brew in a bag to pot, Secure bag with binding clips
- Secure thermometer with binding clips, making sure thermometer isn't touching the bottom
- Follow recipes and get the water up to the right temperatures (takes approx. 1 hr)
- Start mashing (when class starts you want ~10 minutes left on the mash) with lid on
- Have Hops pre-measured out according to the recipes
- Make sure each group has a hydrometer, graduated cylinder, and digital thermometer
- Have the fermenters and auto siphon sterilized with Star San

**Methods:**

1) Pull brew in a bag and squeeze excess water
2) Turn hot plate up to max temp
3) Once boiling, add hops to hop bags and place in pot
4) Boil for 1 hr (lid off)
5) Using the Yeast Viability and Counting protocol, calculate the appropriate volume of cells to add
6) Add wort chiller to pot, put lid on (to prevent contamination) and turn on cold water
    a. Will take ~20 minutes to cool down
    b. Important that the wort is cooled down enough. If the wort is too warm it will kill the yeast

7) Once cool enough move onto taking hydrometer reading.

8) Add appropriate amount of cells to the 1-gallon fermenter.

9) Using an auto-siphon, fill the 1-gallon fermenter 9/10 the way full

10) Add the three-piece airlocks to 1-gallon fermenters.

11) Add 95% ethanol to airlock fill line.

12) Incubate fermenters at room temp for 3 weeks
    a. Be sure to incubate fermenters in the dark or wrap them in foil
    b. Some samples may foam up, so store all fermenters in a secondary container to catch spills.

**Yeast Viability & Counting (~ 20 minutes)**

**Purpose:** To determine the number is live cells to be able to pitch the correct amount of yeast into the wort.

**Information:** The dye **methylene blue** can be used to measure yeast viability. Methylene blue is spontaneously oxidized in the presence of oxygen, resulting in a blue color. Cellular reductases catalyze the conversion of methylene blue to the Leuko form, which is colorless (see below). Thus, live yeast cells that contain active reductases will be colorless, while dead cells lacking the enzymes will remain blue. The number of viable cells can then be quantified using a cell counting chamber (often referred to as a **hemocytometer**).

**Materials:**
Hemocytometers
Microscopes
1.7 ml microcentrifuge tubes
Sterile 10 ml glass tubes
Sterile Erlenmeyer flasks (500 ml and 1000 ml)
0.01% Methylene Blue
Optional: Hand tally counter (VWR 23609-102)

**Before the lab:**

- Have 0.01% Methylene blue in single use aliquots of 990 µl

For students who have an original SG of <60

- Set up 10 ml cultures (in 10 ml glass tubes) and grow at 30°C for 1 day
- Set up 250 ml of sterile wort in 500 ml flask
- Dump the 10 ml culture into a 500 ml flask containing sterile wort and grow at 30°C for 1 day
- Place all cultures at 4°C to crash the cells out (may take 2-3 days)
    - Be careful when transporting cells as they will mix
- Carefully decant media

For students who have an original SG of >61

- Set up 30 ml cultures (in 125 ml flasks) for students who have a target SG of >61 and grow at 30°C for 1 day
- Set up 500 ml sterile wort in 1000 ml flask for students who have an original SG of >61
- Dump the 30 ml culture into a 1000 m flask containing sterile wort and grow at 30°C for 1 day
- Place all cultures at 4°C to crash the cells out (may take 2-3 days)
    - Be careful when transporting cells as they will mix
- Carefully, decant media

**Methods:**

1) Add 10 µl of cells into 990 µl of methylene blue solution (0.01% (w/v)). This constitutes a 1:100 dilution.

2) Vortex briefly and pipette 10 µl of sample into the counting chamber.

3) Place the hemocytometer under a microscope at 10x magnification and find the grid pattern (see below)

4) Count cells using the four corner squares. Count both the total number of cells and the number of live (colorless) cells. For cells falling on a boundary line, only count them if they are on the top or right line

(see below). Count budding cells as separate cells if the bud is at least half the size of the mother cell. **Note: try to be consistent in your counting scheme. See page B23 for a guide.**

5) Each of the nine large squares has a volume of 0.1 $mm^3$ ([1 mm x 1 mm] x 0.1 mm chamber depth). There are 1000 $mm^3$ per ml, so each square has a volume of 0.0001 ml or $10^{-4}$ ml.

So, cells per ml = average # of viable cells (per 4 squares) x $10^4$ $ml^{-1}$ x 100 (dilution factor).

**Hydrometer – Original Gravity (~10 minutes)** (5)

**Purpose:** Hydrometers are used to determine the original gravity (available carbohydrates) of the wort. This gives information that we will use to calculate the alcohol by volume and percent attenuation at the end of the semester.

**Information:** While not a perfect measure, brewers use specfic gravity (SG) to estimate the carbohydrate content of wort. Carbohydrates are the predominant dissolved solids in the wort, and similarly affect density, so SG is a reasonable approximation. SG is often measured using a hydrometer. The hydrometer is properly read at the bottom of the **meniscus**.

**Temperature Correction**

While it is best to take the reading as close as possible to the temperature for which the hydrometer was designed, the following table gives an approximate correction (in points) that must be added to the reading for 15°C. See table 3, the hydrometer temperature corrections.

Table 3

| °F | °C | Correction |
|---|---|---|
| 50 | 10 | -1 |
| 59 | 15 | 0 |
| 68 | 20 | +1 |
| 78 | 25 | +2 |
| 86 | 30 | +3 |
| 95 | 35 | +5 |
| 104 | 40 | +7 |

**Materials:**
Bunsen burners/ethanol burners
Hydrometers
Graduated cylinders (250 ml works best for minimizing loss of wort)
Digital thermometers

**Methods:**

1) Carefully pour beer into graduate cylinder
2) Measure and record the temp of the beer
3) Insert the hydrometer $3/4^{th}$ way into the filled graduated cylinder
4) Spin and release the hydrometer
5) Read the bottom of the meniscus
6) Make sure to correct for the temperature (according to table 3)

**Hydrometer – Final Gravity (~10 minutes) (5)**

**Purpose:** Hydrometers are used to determine the final gravity (available carbohydrates) of the wort after primary fermentation. To carbonate the beer, we will add sugar and have a second round of fermentation called bottle conditioning (page 44). Now we will characterize the beer using the final gravity to calculate the alcohol by volume and percent attenuation (page 45) and determining the color of the beer (page 46).

**Information:** While not a perfect measure, brewers use specfic gravity (SG) to estimate the carbohydrate content of wort. Carbohydrates are the predominant dissolved solids in the wort, and similarly affect density, so SG is a reasonable approximation. SG is often measured using a hydrometer. The hydrometer is properly read at the bottom of the beer **meniscus**. Note that the numbers increase going down.

**Temperature Correction**

While it is best to take the reading as close as possible to the temperature for which the hydrometer was designed, the following table gives an approximate correction (in points) that must be added to the reading for 15°C. See table 4, the hydrometer temperature corrections.

Table 4:

| °F | °C | Correction |
|---|---|---|
| 50 | 10 | -1 |
| 59 | 15 | 0 |
| 68 | 20 | +1 |
| 78 | 25 | +2 |
| 86 | 30 | +3 |
| 95 | 35 | +5 |
| 104 | 40 | +7 |

**Materials:**
Bunsen burners/ethanol burners
Hydrometers
Graduated cylinders (250 ml works best for minimizing loss of wort)
Digital thermometers

**Before the lab:**
- Make sure each group has a hydrometer, graduated cylinder, and digital thermometer

**Methods:**

1) Carefully pour beer into graduate cylinder
2) Measure and record the temp of the beer
3) Insert the hydrometer 3/4$^{th}$ way into the filled graduated cylinder
4) Spin and release the hydrometer
5) Read the bottom of the meniscus
6) Make sure to correct for the temperature

**Bottling (~15 minutes)**

**Purpose:** We need to add sugar to allow a secondary fermentation for proper carbonation of our beer.

**Information:** one volume is 1 L of CO2 at 1 atmosphere of pressure in 1 L of beer. Gas solubility increases at lower temperatures (We ferment at 20°C and the volumes of $CO_2$ is 0.850.) Equation for fermentation: 1 mol glucose -> 2 mol ethanol + 2 mol $CO_2$. Molecular weight of glucose = 180.16; molecular weight of $CO_2$ = 44.01. One 180.16 g of glucose (1 mole) yields 88.02 g of $CO_2$ (2 moles). One mole (44.01g) of $CO_2$ at standard pressure occupies 22.4 liters (ideal gas law). 1.96 g of CO2 occupy 1 L (44.01 g / 22.4 L), which is produced by 4.01 g glucose. Simplified: add 4.01 g glucose per L per volume.

**Materials**:
Star San (food grade, no rinse sanitizer)
Swing neck amber bottles (clear requires fermenting in the dark – prevent "skunking" of the beer) (2/brewing group)
10 ml serological pipettes (2/brewing group)
Pipette bulbs (or automatic pipettors)
Sterilized 50% corn sugar (50% glucose)
Hand operated auto siphon with tube (1/brewing group)
Ethanol burners/Bunsen burners
95% EtOH
Sterile 50 ml conical tube (1/brewing group)

**Before the lab:**
- Have the bottles and auto siphon sterilized with Star San
- Make 50% corn sugar and aliquot each brewing group (both beers) 25 ml into a 50 ml conical

**Methods:**

1) Look up the recommended volumes of $CO_2$ of your beer style
2) 2.9 volumes (depend on your type of beer) – 0.85 volumes (see information section) = 2.05 volumes (the amount we want to add)
3) Assume bottles are 16 ounces or 0.47 liters
4) 2.05 volumes x 0.47 liters x 4.01 g glucose (see information section) = 3.86 g glucose
5) Our glucose stock is 50% w/v (or 0.5 g glucose/ml), so 3.86 g (must divide by 0.5 to calculate volume to add) = 7.72 ml
6) Under a flame, add the appropriate volume of 50% glucose to bottle
7) Using auto-siphon (be careful to not disturb the cells at the bottom) carefully fill glucose containing swing-neck bottle to 2 inches below the stopper.
8) Incubate at room temp (in the dark) for 2.5 weeks.
9) Using auto-siphon fill a 50 ml conical tube to be used for SRM color assay and Hop bitterness assay.

**ABV & Percent Attenuation (~ 5 minutes)**

**Purpose:** Determine that alcohol by volume and calculate the percent attenuation of our beer.

**Information:** Alcohol by volume (ABV) can be estimated using specific gravity (density of a solution relative to water). Water inherently has a specific gravity of 1.000 (density of water/ density of water). Sugars have a higher density and ethanol has a lower density (specific gravity = 0.794). Thus, a simple formula to estimate ABV is to assume that the change in original gravity (OG) to final gravity (FG) is due to ethanol:

**More Accurate Equation**
**ABV = [76.08 x (OG-FG) / (1.775 - OG)] x (FG / 0.794)**

**where [76.08 x (OG-FG) / 1.775 – OG] represents the alcohol percentage by weight, and 0.794 denotes the specific gravity of ethanol**

Apparent attenuation is the percentage of sugars that the yeast consumed, which can be also be calculated using the change in OG to FG:

**Apparent Attenuation (%) = ((OG-FG) / (OG-1)) x 100**

The following are common yeast attenuation ranges:

Low = 65-70%
Medium = 71-75%
High = 76-85%

**Standard Reference Method for Beer Color (~5 minutes)**

**Purpose**: Determine the color of our beer.

**Information:** The Standard Reference Method (SRM) for measuring beer color was adopted by the American Society of Brewing Chemists in 1950 as an alternative to the Lovibond system. The SRM method relies on visible absorbance at 430 nm.

**Materials:**
P1000 tips
Spectrometer
Cuvettes (2/brewing group)
Sterile water

**Before the lab:**

- Turn on spectrometer (some specs require ~15 minutes to warm up)

**Methods:**

1) Pipette 1 ml of beer into a cuvette (If the beer is dark, you will need to make a 10-fold dilution).

2) Add 1 ml of blank (sterile water) to a cuvette.

3) Measure the absorbance of the sample at 430 nm.

4) SRM = Absorbance at 430 nm x 12.7 x dilution factor.

**Hop Bitterness (~12-90 minutes) (7)**

**Purpose:** Determine how bitter our beer is by quantifying the amount of the iso-alpha acids.

**Information:** The alpha acids in hop flowers are isomerized during wort boiling to form **iso-alpha-acids**. Humulone shown below is the most prevalent alpha acid in hops.

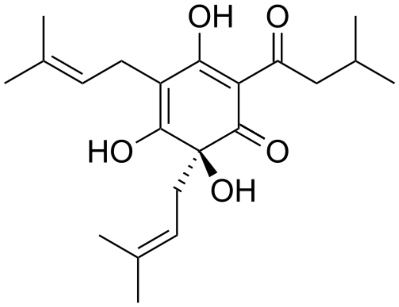

humulone

Image taken from Wikipedia

The hop iso-alpha-acids contribute to the bitterness of the beer. The amount of iso-alpha acids is generally reported on the **International Bitterness Units** (IBU) scale. One IBU equals 1 mg of isomerized alpha acid per liter. Because iso-alpha-acids absorb strongly at 270 nm, a spectrophotometer can be used for quantification.

Hop iso-alpha-acids are slightly hydrophobic, and their hydrophobicity increases at low pH. Acidifying the beer allows iso-alpha acids to be extracted into hydrophobic organic solvents. Iso-octane is the standard solvent used in the brewing industry, since it has a low absorbance at 270 nm.

IBU is calculated as the absorbance multiplied by a factor of 50. This factor is derived from the dilution factor and an assumption that 70% of the absorbance of the iso-octane solution is due to iso-alpha-acids.

**Warning: iso-octane is highly flammable and toxic by inhalation, ingestion, or skin absorption. Hydrochloric Acid is highly corrosive and toxic by inhalation, ingestion, or skin absorption**

**Materials:**
50 ml conical tubes (2/brewing group, 1 for Instructor to make the blank)
Microcentrifuge tubes (for octyl alcohol and 3N HCl)
Octyl alcohol
Iso-octane (2,2,4-trimethylpentane)
3 N hydrochloric acid
P1000 tips
10 ml serological pipettes
Pipette bulbs (or automatic pipettors)
Vortex mixers
UV Spectrometer
Quartz cuvette (Fisher NC9416015) – fragile, only Instructor handles

Optional materials:
Table top centrifuge capable of spinning 50 ml conical tubes

**Before the lab:**
- Turn on spectrometer (some specs require ~15 minutes to warm up)
- Make single use aliquots of octyl alcohol, iso-octane, and hydrochloric acid
- Make blank (20 ml iso-octane plus 50 µl octyl alcohol)

**Methods:**

1) Pipette 10 ml of beer into a 50 ml conical tube containing 20 ml iso-octane.
2) Add 50 µl octyl alcohol and 1 ml 3 N hydrochloric acid.
3) Close the tube (make sure it is well sealed) and vortex for 15 min.
4) Centrifuge at 3000 x g for 5 min to allow phase separation to occur (or allow gravity to separate the phases generally 60 min), **in the dark (iso-alpha-acids are UV sensitive).**
5) Add 1 ml of blank (20 ml iso-octane plus 50 µl octyl alcohol) to a UV transparent cuvette (quartz).
    a. Note: the iso-octane will cloud up plastic cuvettes
6) Transfer 1 ml of the sample (upper organic phase) to a cuvette.
7) Measure the absorbance of the sample at 275 nm.
8) IBU = Absorbance at 275 nm x 50.

**Table of Contents**

| Title of the Lab | Page Number |
|---|---|
| Notes | 1 |
| Wild Yeast Enrichment and Isolation | 2-4 |
| Yeast genomic DNA Extraction | 5 |
| ITS PCR for Identification of Yeast Species | 6 |
| DNA Electrophoresis of ITS PCR | 7 |
| PCR Cleanup | 8 |
| ITS Sequencing | 9 |
| ITS sequencing computer lab | 10 |
| Mashing | 11 |
| Starch Iodine Test | 12 |
| Calculating Original Gravity | 13-14 |
| Calculating Fermentation Rates | 15-16 |
| Yeast Viability and Counting | 17-18 |
| Small Scale Fermentation | 19 |
| Yeast Collections for qPCR and GC-MS | 20 |
| RNA Extraction | 21-23 |
| RNA Electrophoresis | 24 |
| cDNA Synthesis | 25 |
| RNA Hydrolysis and cDNA cleanup | 26-27 |
| qPCR | 28-29 |
| GC-MS | 30 |
| qPCR computer lab | 31 |
| GC-MS computer lab | 32 |
| Brewing Protocol | 33 |
| Yeast Viability and Counting | 34 |
| Hydrometer – Original Gravity | 35 |
| Hydrometer – Final Gravity | 36 |
| Bottling | 37 |
| ABV and Percent Attenuation | 38 |
| Standard Reference Method for Beer Color | 39 |
| Hop Bitterness | 40-41 |
| References | 42 |

**Notes:**

1) BSL2-level protocols must be used when working with unknown environmental samples. Students must show mastery of BSL1-level techniques before performing BSL2-level activities.

2) Gloves, goggles, and lab coats must be worn for every experiment.

**Wild Yeast Enrichment and Isolation** (1)

**Purpose:**

**Materials**

**Yeast Enrichment Media 1 (YEM1)**

3 g/L yeast extract (0.3% final)
3 g/L malt extract (0.3% final)
5 g/L bacto peptone (0.5% final)
10 g/L sucrose (1% final)

Bring volume to 915.5 ml with dd$H_2O$

---

autoclave 20 min liquid cycle or filter sterilize

---

once cool, add:

84 ml/L 95% ethanol (8% final)
50 µl of 20 mg/ml chloramphenicol (1 µg/ml final)
415 µl 6N HCl

Check pH using pH paper—should be pH 5-6.

**Yeast Enrichment Media 2 (YEM2)**

5 g/L Yeast Nitrogen Base (YNB) without amino acids (Sigma # Y0626)
2 g/L Yeast Synthetic Drop Out Supplements (YSDO) (see below)
20 g/L alpha-D-methylglucoside

Bring volume to 1000 ml with dd$H_2O$ and transfer to a 2 L flask

20 g bacto agar

---

autoclave 20 min liquid cycle

---

add 1.3 ml 6N HCl

Check pH using pH paper—should be pH 3-4
Pour plates (~25 ml per plate)

Sterile spreaders (or glass beads) (3/student)
Sterile petri plates
P200 tips
Gallon size zip lock bags (1/student)
Quart size zip lock bags (3/student)
Sterile 50ml conical tubes (3/student)
Wild yeast collection card (3/student)
Bunsen burners/ethanol burners
YEM1 (150ml/student)
YEM2 plates (6/student)
Gloves (3 pairs/student)

**YNB Components**
(μg/L unless indicated)
Nitrogen Sources:
Ammonium sulfate, 5.0 g/L

Vitamins:
Biotin, 2.0
Calcium pantothenate, 400
Folic acid, 2.0
Inositol, 2.0 mg/L
Nicotinic acid, 400
p-Aminobenzoic acid, 200
Pyridoxine HCl, 400
Riboflavin, 200
Thiamine HCL, 400

Trace Elements:
Boric acid, 500
Copper sulfate, 40
Potassium iodide, 100
Ferric chloride, 200
Manganese sulfate, 400
Sodium molybdate, 200
Zinc sulfate, 400

Salts:
Potassium phosphate monobasic, 1.0 g/L
Magnesium sulfate, 0.5 g/L
Sodium chloride, 0.1 g/L
Calcium chloride, 0.1 g/L

**YSDO Components** (mg/L)
Adenine, 18
p-Aminobenzoic acid,8
Leucine, 380
Alanine, 76
Arginine, 76
Asparagine, 76
Aspartic acid, 76
Cysteine, 76
Glutamic acid, 76
Glutamine, 76
Glycine, 76
Histidine, 76
myo-Inositol, 76
Isoleucine, 76
Lysine, 76
Methionine, 76
Phenylalanine, 76
Proline, 76
Serine, 76
Threonine, 76
Tryptophan, 76
Tyrosine, 76
Uracil, 76
Valine, 76

**Methods:**

**Day 1 (~10 minutes)**

1) Give students Wild Yeast Collection packets

**Day 2 (~10 minutes)**

1) Add 50 ml YEM1 to sterile 50 ml conical tube containing collection material and tighten cap.
2) Vortex well, tighten caps until just closed, and then loosen by ¼ turn.
3) Place at room temperature in the dark. (may wrap tubes in aluminum foil)

**Day 4 and Day 8 (~15 minutes)**

1) Observe samples for fermentation by looking for the production of bubbles.
    a. If bubbles are absent on Day 4, continue to incubate the plate until Day 8.

2) For samples containing bubbles, spread 50 µl onto YEM2 plates.

3) Incubate at 30°C for 2 days.

**Day 6 and/or Day 10 (~30 minutes)**

1) Add 10 µl sterile water to a glass slide.

2) Stab a putative yeast colony with 200 µl pipet tip and mix with the water on the glass slide.

    a. Yeast colonies are generally smooth and cream colored.

3) Use a microscope to observe whether the colony contained budding cells.

4) For colonies containing budding yeast, streak the colony to a fresh YEM2 plate to isolate single colonies.

5) Incubate at 30°C for 2-3 days.

**Yeast genomic DNA Extraction (~30 minutes)** (2)

**Note:** This lab will be completed along with ITS PCR for Identification of Yeast Species (page C6)

**Purpose:**

**Materials:**

LiOAc Lysis Buffer:

1 ml 2 M Lithium Acetate (200 mM final)
1 ml 10% SDS (1% final)
8 ml sterile water
10 ml total (filter sterilize or use filter sterilized reagents)

TE:

10 mM Tris-HCl, pH 8.0,
1 mM EDTA

Sterile 1.7 ml microfuge tube (2/student)
Sterile P200 tips
Bunsen burners/ethanol burners
100% Ethanol
Microcentrifuge
Heat block/thermomixer

**Methods:**

1) Add 100 µl LiOAc Lysis buffer to a sterile 1.7 ml microfuge tube.
2) Stab a single colony 8x with sterile P200 tip, add to the 100 µl LiOAc Lysis buffer.
3) Incubate at 70°C for 15 min.
4) Add 300 µl of 100% EtOH and vortex well.
5) Spin at 15000 x g or max speed in microcentrifuge for 3 min.
6) Decant supernatant. **Note: if pellet becomes dislodged, do not pour, use a pipet**
7) Resuspend pellet in 100 µl TE
8) Spin at 15000 x g or max speed in microfuge for 1 min,
9) Carefully pipet supernatant to new sterile 1.7 ml microfuge tube.
10) Use 1 µl per 50 µl PCR.
11) Store samples at -20°C

**ITS PCR for Identification of Yeast Species (~10 minutes)** (3)

**Purpose:**

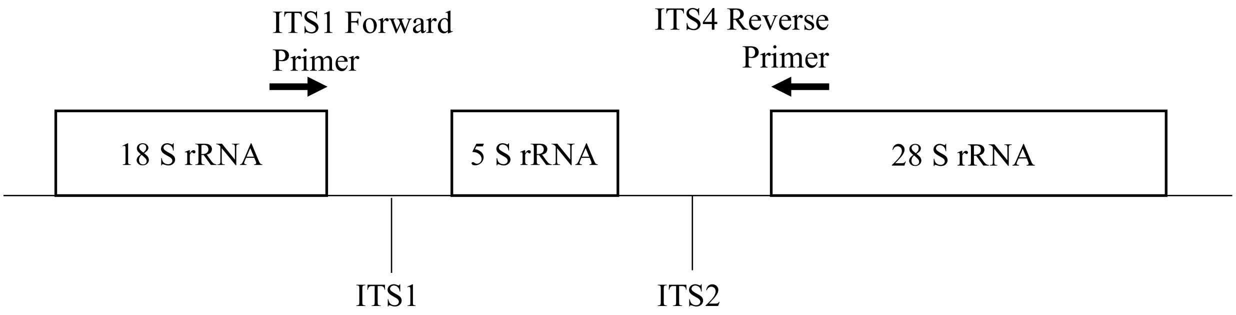

Image made by Amanda Scholes

**Materials:**

PCR Master Mix:

8 µl of 50 µM ITS1 primer (5'-TCCGTAGGTGAACCTGCGG-3'; 0.5 µM final)
8 µl of 50 µM ITS4 primer (5'-TCCTCCGCTTATTGATATGC -3'; 0.5 µM final)
8 µl of 25 mM dNTP mixture (250 µM final)
80 µl Taq Reaction Buffer (10x stock, 1x final concentration, preferably green)
664 µl nuclease-free water
768 µl total

Sterile PCR tubes (1/student)
Sterile P10 tips
Bunsen burners/ethanol burners
RNase-Free filter P10 tips (Instructor only)

**NOTE: Keep Taq polymerase in freezer/freezer block until ready for use.**

**Methods:**

1) Add 96 µl of PCR Master Mix to a PCR tube.
2) Add 2 µl of yeast genomic DNA.
3) Add 2 µl of Taq polymerase (obtain from the TA)
4) Mix well by pipetting gently. **Note: Do not vortex enzymes, they can be mechanically denatured by air bubbles.**

**PCR Cycling Conditions:**

94°C ----> 5 min
94°C ----> 30 sec. \
50°C ----> 30 min x 40 cycles
65°C ----> 2 min. /
65°C ----> 10 min

**DNA Electrophoresis of ITS PCR (~30-55 minutes)**

**Note:** This lab will be completed along with PCR Cleanup (page C8), and ITS Sequencing (page C9).

**Purpose:**

**Materials:**
Gel apparatus with combs
Gel visualizer
1X TBE Running buffer
Agarose
Smart-Glow DNA Gel Dye
DNA ladder
*optional loading dye
Sterile P10 tips

**Methods:**

1) Pipette gently to mix sample
2) Load 5 µl into well
    a. Be careful agarose gels can tear easily with tips
    b. If samples are clear will need to add loading dye
3) Run gel
    a. 120V for 45 min or 150V for 20 min
4) Visualize gel

**PCR Cleanup (~30 minutes)**

**Purpose:**

**Materials:**
PureLink Quick gel extraction and PCR cleanup combo kit, 50 preps (ThermoFisher #K220001)
Sterile 1.7 ml microfuge tube (2/student)
Sterile P200 tips
Sterile P1000 tips
Bunsen burners/ethanol burners
Microcentrifuge
Nanodrop Spectrophotometer (or standard spectrophotometer with UV-transparent micro-cuvettes)

**Methods:**

1) Transfer PCR product to sterile 1.7 ml microfuge tube
2) Add 360uL B2 (Binding buffer) to tube containing PCR reaction (50–100 µl) and mix well by pipetting.
3) Add sample in Binding Buffer from step 2, to the PureLink® Spin Column.
4) Centrifuge the PureLink® Spin Column at room temperature at 10,000 x g for 1 min.
5) Discard the flow through
6) Replace the PureLink® Spin Column into the Wash Tube.
7) Add 650 µl W1 to the PureLink® Spin Column.
8) Centrifuge the PureLink® Spin Column at room temperature at 10,000 x g for 1 min.
9) Discard the flow-through from the Wash Tube and replace the PureLink® Spin Column into the tube.
10) Centrifuge the PureLink® Spin Column at maximum speed at room temperature for 3 min to remove any residual W1.
11) Discard the Collection Tube.
12) Place the PureLink® Spin Column in a new sterile 1.7 ml microfuge tube
13) Add 15 µl E1 (Elution Buffer) sterile, distilled water to the center of the PureLink® Spin Column.
14) Incubate the PureLink® Spin Column at room temperature for 1 min.
15) Centrifuge the PureLink® Spin Column at maximum speed for 1 min.
16) The elution tube contains the purified PCR product.
17) Discard the PureLink® Spin Column.
18) Quantify the samples with spectrophotometer.
19) DNA can be stored at -20°C for long-term storage.

### ITS Sequencing – Instructor Only

#### Purpose:

#### Methods:

1) Sample sent for sequencing.

**ITS Sequencing computer lab (~40 minutes)**

**Purpose:**

**Materials:**
SnapGene Viewer (another sequence viewer)
Computer with internet connection

**Methods:**

1) Download your sequencing file (.scf) and open in SnapGene Viewer
2) In SnapGene click the box that shows quality score
3) Looking at the chromatogram, copy a region of the sequencing with high quality (>300 bp)
4) Open NCBI Nucleotide Blast
5) Paste sequence, uncheck filter "Low Complexity regions" and click "BLAST"
    a. The run might take a few minutes
6) Students will take a screenshot and note the top several species by lowest E value and high percent identity.
    a. If < 97% sequence identity, could be a new species
    b. Note: For later analyses and brewing, we only moved forward with *Saccharomyces cerevisiae.*

**Mashing (~90 minutes)**

**Note:** This lab will be completed along with Starch Iodine Test (page C12), and Calculating the Original Gravity (page C13)

**Purpose:**

**Materials:**
20-Quart Stainless Steel Basic Stockpot with lid
The Brew Bag for Kettles (brew in a bag)
Binding clips (3/group)
Digital thermometer
Wort chiller
UltraPure Water (water with minimal ions)
Electric single burner hot plate
Extra-long spoons (>24 inches)
Thick kitchen gloves
1.7 ml microcentrifuge tubes (3/group)

**Methods:**

1) Add 3 lb of 2 row grain to brew-in-a-bag in pots
2) Mash for 1 hr at 67°C with the lid on, stirring occasionally (approximately every 10 min)
    a. Pulling 500 µl samples at 0 min, 15 min, 30 min, 45 min, and 60 min (and move onto the starch conversion assay – protocol below)
3) Pull brew in a bag and squeeze excess water
4) Turn off and remove pot from hot plate
5) Add wort chiller to pot and turn on cold water
    a. Will take ~20 minutes to cool down
6) Add ~4 L of wort to a 20 L plastic beaker
    a. Mixing wort form all tables together for homogenous beer media
7) Fill 4-1 L bottles and autoclave to sterilize wort per table
    a. Autoclave on a liquid 20 min cycle
    b. There will be some protein that crashes out (this is normal

**Starch Iodine Test (~10 minutes)** (4)

**<u>Purpose:</u>**

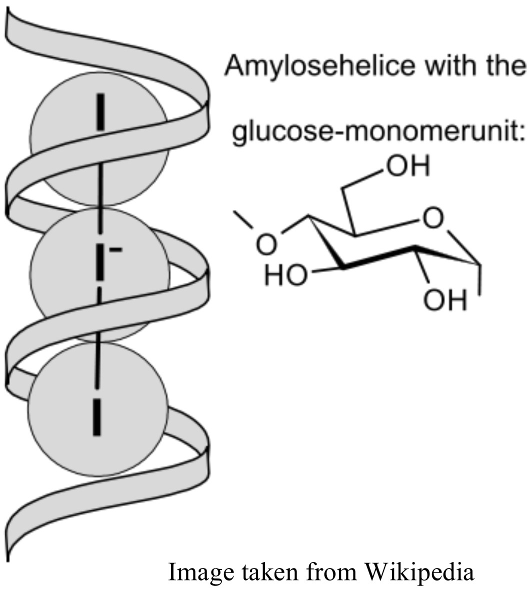

Image taken from Wikipedia

**<u>Information:</u>** Starch and the triiodide anion ($I_3^-$) interact to form a complex starch helix around the iodide resulting in a blue-black color. In the absence of starch, iodine solutions have a brown color

**<u>Hazard:</u>** Iodine can cause eye, skin, and respiratory tract irritation or an allergic reaction. Wear eye protection and gloves.

**<u>Materials:</u>**
50% glucose (corn sugar)
50% corn starch
0.025N iodine
Small weigh boats (7/group)
P1000 tips
P1000 pipet

**<u>Methods:</u>**

1) Add 500 µl of 50% glucose into a small weigh boat (enough to fill the bottom).

2) Repeat for cornstarch and every timepoint sample.

3) Add 3-4 drops 0.025 N iodine mix with pipet tip.

4) Record the color

**Calculating Original Gravity (~10 minutes)** (5)

**<u>Purpose:</u>**

**<u>Information:</u>** While not a perfect measure, brewers use specfic gravity (SG) to estimate the carbohydrate content of wort. Carbohydrates are the predominant dissolved solids in the wort, and similarly affect density, so SG is a reasonable approximation. SG is often measured using a hydrometer. The hydrometer is properly read at the bottom of the **meniscus**. Note that the numbers <u>increase</u> going down. In the example below, the hydrometer has a SG reading of 1.047, or 47 points (aka called gravity units or GU).

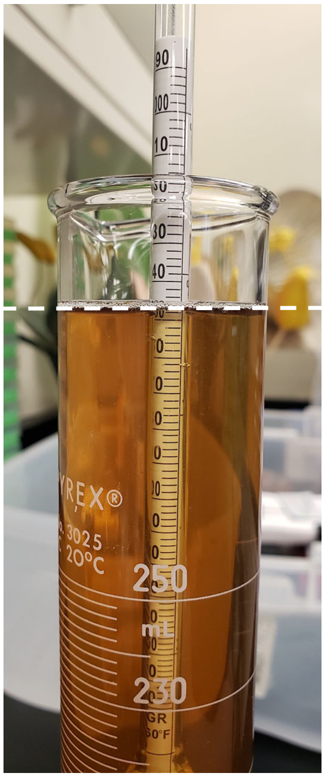

Figure 1: Hydrometer measuring SG of wort. Image taken by Amanda Scholes

Table 1:

| SG | Percent | SG | Percent | SG | Percent | SG | Percent |
|---|---|---|---|---|---|---|---|
| 1 | 0.25 | 21 | 5.33 | 41 | 10.24 | 61 | 14.98 |
| 2 | 0.51 | 22 | 5.58 | 42 | 10.48 | 62 | 15.21 |
| 3 | 0.77 | 23 | 5.83 | 43 | 10.72 | 63 | 15.45 |
| 4 | 1.03 | 24 | 6.08 | 44 | 10.96 | 64 | 15.68 |
| 5 | 1.28 | 25 | 6.33 | 45 | 11.20 | 65 | 15.91 |
| 6 | 1.54 | 26 | 6.58 | 46 | 11.44 | 66 | 16.14 |
| 7 | 1.80 | 27 | 6.82 | 47 | 11.68 | 67 | 16.37 |
| 8 | 2.05 | 28 | 7.07 | 48 | 11.92 | 68 | 16.60 |
| 9 | 2.31 | 29 | 7.32 | 49 | 12.16 | 69 | 16.83 |
| 10 | 2.56 | 30 | 7.56 | 50 | 12.40 | 70 | 17.06 |
| 11 | 2.81 | 31 | 7.81 | 51 | 12.63 | 71 | 17.29 |
| 12 | 3.07 | 32 | 8.05 | 52 | 12.87 | 72 | 17.51 |
| 13 | 3.32 | 33 | 8.30 | 53 | 13.11 | 73 | 17.74 |
| 14 | 3.57 | 34 | 8.54 | 54 | 13.34 | 74 | 17.97 |
| 15 | 3.82 | 35 | 8.79 | 55 | 13.58 | 75 | 18.20 |
| 16 | 4.07 | 36 | 9.03 | 56 | 13.81 | 76 | 18.43 |
| 17 | 4.33 | 37 | 9.27 | 57 | 14.05 | 77 | 18.65 |
| 18 | 4.58 | 38 | 9.52 | 58 | 14.28 | 78 | 18.88 |
| 19 | 4.83 | 39 | 9.76 | 59 | 14.52 | 79 | 19.10 |
| 20 | 5.08 | 40 | 10.00 | 60 | 14.75 | 80 | 19.33 |

**Temperature Correction**

The hydrometer scale (table 1) above is calibrated to the mass percent of sucrose that has the indicated SG at 20°C. However, water expands as the temperature increases, causing the density to decrease. If the hydrometer is used at a different temperature, a correction must be applied. While it is best to take the reading as close as possible to the temperature for which the hydrometer was designed, the following table gives an approximate correction (in points) that must be added to the reading for 15°C. See table 2, the hydrometer temperature corrections.

Table 2:

| °F | °C | Correction |
|---|---|---|
| 50 | 10 | -1 |
| 59 | 15 | 0 |
| 68 | 20 | +1 |
| 78 | 25 | +2 |
| 86 | 30 | +3 |
| 95 | 35 | +5 |
| 104 | 40 | +7 |

**Materials:**

Bunsen burners/ethanol burners
Hydrometers
Graduated cylinders (250 ml works best for minimizing loss of wort)
Digital thermometers

**Methods:**

1) Carefully pour wort into graduate cylinder
2) Measure and record the temp of the beer
3) Insert the hydrometer 3/4$^{th}$ way into the filled graduated cylinder
4) Spin and release the hydrometer
5) Read the bottom of the meniscus
6) Make sure to correct for the temperature (according to table 2)
7) Record value in lab notebook

**Calculating Fermentation Rates (~30 minutes)**

**Note:** This lab will be completed with Yeast Viability and Counting (page C17) and Small Scale Fermentation (page C19).

**Purpose:**

**Information:**
The following equation represents the fate of glucose, energy and reducing equivalents during yeast fermentation:

$$\underset{\text{glucose}}{C_6H_{12}O_6} + 2ADP + 2Pi + 2NAD^+ \rightarrow \underset{\text{pyruvate}}{2C_3H_4O_3} + 2ATP + 2NADH \rightarrow \underset{\text{acetaldehyde}}{2C_2H_4O} + 2CO_2 \rightarrow \underset{\text{ethanol}}{2C_2H_5OH} + 2NAD^+$$

This assay measures fermentation efficiency as the volume of $CO_2$ released per minute per cell mass.

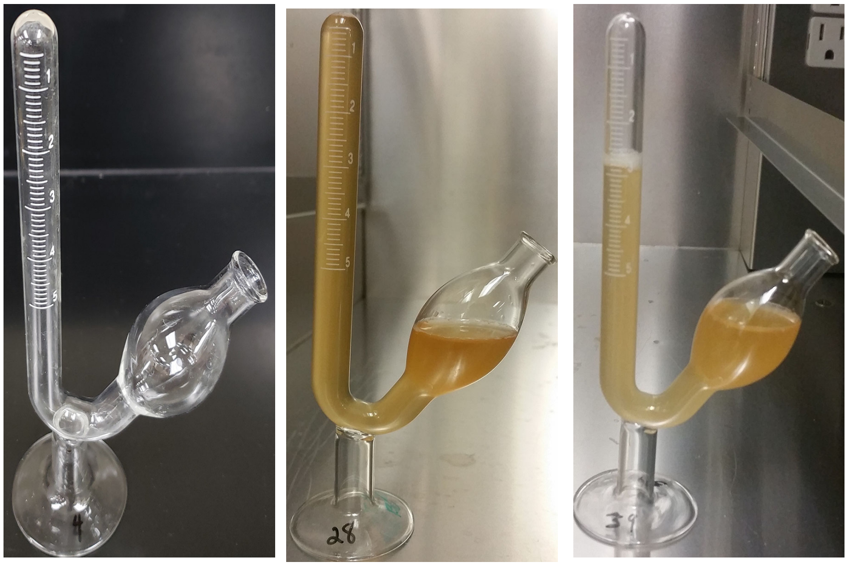

Figure 2: The first panel shows an empty fermentation tube. Second panel shows fermentation tube once filled and about to start the fermentation test. Panel 3 fermentation tube in the middle of the fermentation test. Images taken by Amanda Scholes

**Materials:**
Vortex
Spectrometer
Cuvettes (1/group, Instructor will need one for a media blank)
Wort (made previously)
Fermentation Tube Graduated 5 ml (Science Lab Supplies B2800-2) (2/group)
10 ml glass culture tubes
50 ml conical tubes (1/culture)
Table top centrifuge capable of spinning 50 ml conical tubes
10 ml serological pipets
Pipet bulb/electric pipettor

**Methods:**

1) Decant supernatant.
2) Resuspend cell pellet with 10 ml sterile wort via vortexing.
3) Carefully transfer half the cells into fermentation tube
4) Plug opening and invert tube (getting cells into the graduated chamber)
5) Place tube upright and add remaining cells
    a. **Fermentation tubes are VERY fragile. Be careful when handling.**
6) Monitor and record every 2 minutes once it passes the 0.5 ml mark.
7) Make 1:25 dilution of cells into wort (40 μl cells into 960 μl wort) and make blank (1 ml wort)
8) Spec cells to determine optical density (OD).
9) Plot ml $CO_2$ displaced vs time in minutes.
10) To determine fermentation rate divide slope by the OD.

**Yeast Viability and Counting (~ 20 minutes)**

**Purpose:**

**Information:** The dye **methylene blue** can be used to measure yeast viability. Methylene blue is spontaneously oxidized in the presence of oxygen, resulting in a blue color. Cellular reductases catalyze the conversion of methylene blue to the Leuko form, which is colorless (see below). Thus, live yeast cells that contain active reductases will be colorless, while dead cells lacking the enzymes will remain blue.

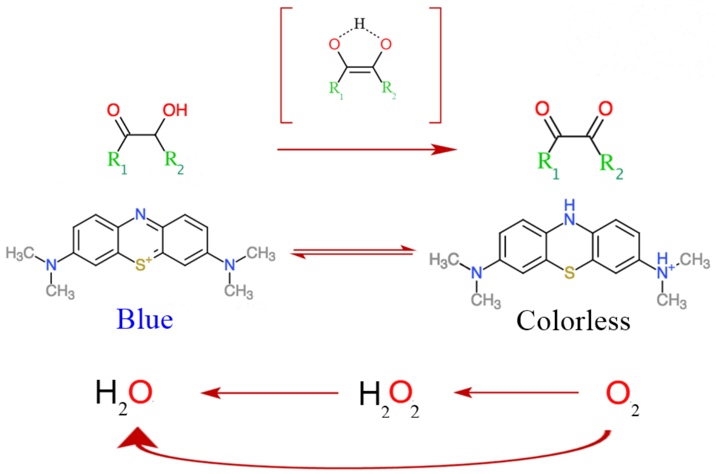

Figure 3: schematic of how the color changing mechanism works. Image from Wikipedia.

The number of viable cells can then be quantified using a cell counting chamber (often referred to as a **hemocytometer**).

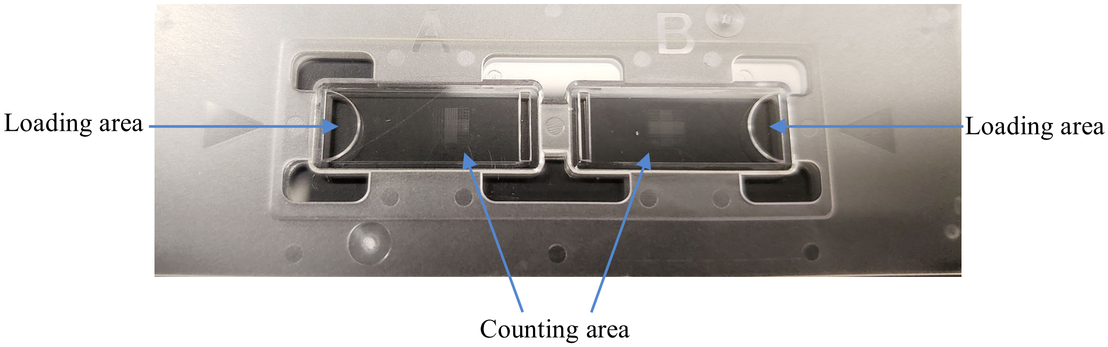

Figure 4: Hemocytometer slide. Image taken by Amanda Scholes

**Materials:**
Hemocytometers
Microscopes
Sterile 50 ml conical tubes
1.7 ml microcentrifuge tubes
Sterile 10 ml glass tubes
Sterile Erlenmeyer flasks (125 ml)
Sterile wort
0.01% Methylene Blue
Optional: Hand tally counter (VWR 23609-102)

**Methods:**

1) Add 10 µl of cells into 990 µl of methylene blue solution (0.01% (w/v)). This constitutes a 1:100 dilution.

2) Vortex briefly and pipette 10 µl of sample into the counting chamber.

3) Place the hemocytometer under a microscope at 10x magnification and find the grid pattern (see below)

4) Count cells using the four corner squares. Count both the total number of cells and the number of live (colorless) cells. For cells falling on a boundary line, only count them if they are on the top or right line (see below). Count budding cells as separate cells if the bud is at least half the size of the mother cell. **Note: try to be consistent in your counting scheme.**

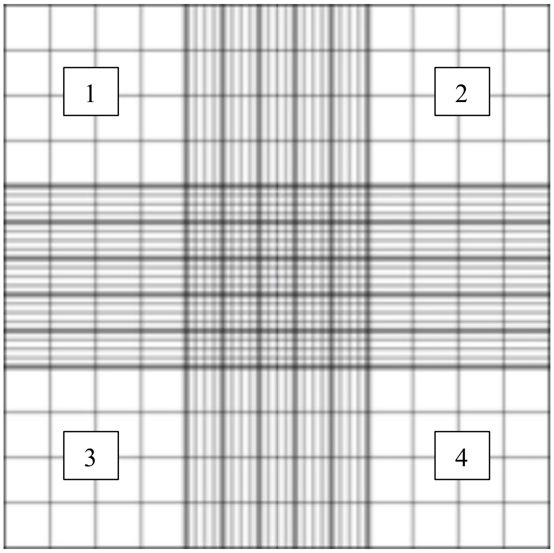

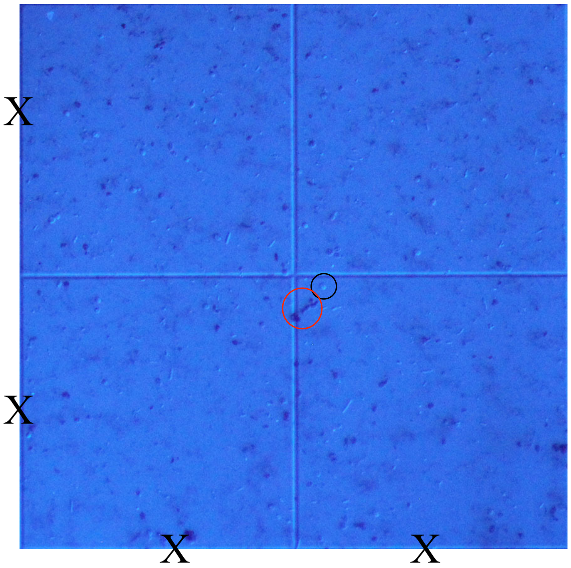

The black circle is showing a live cell (colorless), while the red circle shows dead cells (dark blue).

5) Each of the nine large squares has a volume of 0.1 $mm^3$ ([1 mm x 1 mm] x 0.1 mm chamber depth). There are 1000 $mm^3$ per ml, so each square has a volume of 0.0001 ml or $10^{-4}$ ml.

So, cells per ml = average # of viable cells (per 4 squares) x $10^4$ $ml^{-1}$ x 100 (dilution factor).

**Small Scale Fermentation (~20 minutes)**

**Purpose:**

**Materials:**
Sterile 125 ml Erlenmeyer flasks (1/group)
Sterile wort (from Mashing protocol pg16)
No. 6 Stopper – Drilled (Northern Brewer Homebrew Supply SKU: SD6) (1/group)
Three-Piece Airlock (Northern Brewer Homebrew Supply SKU: 7010) (1/group)
95% ethanol

**Methods:**

1) Add 625 million cells to flask containing 100 ml sterile wort
2) Add sterile flask stopper onto airlock
3) Plug flask with stopper and airlock
4) Fill airlock with 95% ethanol
5) After 2 days proceed to Yeast Collections for qPCR and GC-MS (protocol below).

**Yeast Collections for qPCR and GC-MS (~10 minutes)**

**Purpose:**

**Materials:**
50 ml conical tubes (2/group)
Liquid Nitrogen
-80°C freezer

**Methods:**

1) Label (lid and side) 2-50 ml conical tubes with sample name and
    a. Tube 1 GC-MS
    b. Tube 2 RNA
2) Carefully **(minimally disturbing the cells)** remove airlock and decant 20 ml of supernatant into 50 ml conical tube labeled GC-MS
3) Swirl flask and decant cells into 50 ml conical tube labeled RNA.
4) Spin down cells at 1500 x g for 3 min
5) Flash free cells in liquid nitrogen
6) Store cells at -80°C freezer, until ready to be used

**RNA Extraction (~50 minutes)**
modified Zymo Research Protocol

**Purpose:**

**Materials:**
Quick-RNA MiniPrep Plus kit (Zymo Research R1057)
RNase-Free 1.7 ml microcentrifuge tubes
RNase-Free 2.0 ml microcentrifuge tubes
RNase-Free Filter P100 tips
RNase-Free Filter P1000 tips
RNase-Free PCR tubes
100% RNase-Free Ethanol
Bunsen burners/ethanol burners
Vortex
MultiTherm
Microcentrifuge
Nanodrop/Qubit Spectrometer
Ice/ice bin

**Note: All steps following cell disruption should be performed at room temperature. Work quickly to limit RNA degradation.**

**Note: Do not touch the column resin with tips.**

**Methods:**

1. Resuspend cell pellet in 400 µl with 1X DNA/RNA Shield – Instructor does.
2. Label (lid and side) **two sets** of RNase-free 1.7 ml microcentrifuge tubes, one set of 2.0 ml microcentrifuge tube, one set of Spin Away Filters (Yellow column), and Zymo spin columns (Green Zymo Spin III CG column).
3. Add 400 µl cells (in 1X DNA/RNA Shield) to 400 µl glass beads, and homogenize by placing in MultiTherm set to 4°C and maximum shaking for 10 min.
    a. Vortex samples every two minutes.
4. Add 400 µl L1 (RNA Lysis Buffer) and gently mix by pipetting.
5. Carefully pipette sample into a Yellow Spin Away Filter column (column and collection tube).
6. Centrifuge for 30 sec at 16,000 x g. **Transfer flow through by pipetting to a new RNase-Free 2.0 ml tube**
7. Add 800 µl EtOH (100% EtOH) and gently mix by pipetting.
8. Carefully pipette sample to a Green Zymo Spin III CG mini spin column.

9. Centrifuge for 30 sec at 16,000 x g.
10. Discard flow through by pipetting.
11. Add 400 µl W1 (RNA Wash Buffer) to the column.
12. Centrifuge for 30 sec at 16,000 x g.
13. Discard flow through by pipetting.
14. Add 80 µl D1 (DNase I) directly to the column.
15. Incubate the column at room temperature for **25 min** to digest genomic DNA.
16. Add 400 µl P1 (RNA Prep Buffer) to the column.
17. Centrifuge for 30 sec at 16,000 x g.
18. Discard flow through by pipetting.
19. Add 700 µl W2 (RNA Wash Buffer).
20. Centrifuge for 30 sec at 16,000 x g.
21. Discard flow through by pipetting.
22. Add 400 µl W3 (RNA Wash Buffer).
23. Centrifuge for 2 min at 16,000 x g.
24. Transfer column to a new RNase-Free 1.7 ml microcentrifuge tube (**DO NOT let touch the column the flow through**).
25. Add 50 µl E1 (RNase-free water) and incubate for 1 min at room temp.
    a. **Note: Make sure to pipette water directly onto column resin WITHOUT touching pipette tip to membrane. If water remains on the wall of the column, tap the tube until water absorbs into the membrane.**
26. Centrifuge for 30 sec at 16,000 x g.
27. Carefully pipette the eluted RNA and place it back onto column, incubate for an additional 1 min.
    a. **Note: the second elution increases RNA yield. Note: Make sure to pipette water directly onto column resin WITHOUT touching pipette tip to membrane. If water remains on the wall of the column, tap the tube until water absorbs into the membrane.**
28. Centrifuge for 1 min at 16,000 x g.

29. Transfer Eluted RNA to a fresh RNase-Free 1.7 ml microcentrifuge tube just in case any RNases contaminated the lid of the tube during centrifugation. **Immediately place RNA on ice.**
    a. Long term storage is at -80°C, and RNA concentrations are calculated using a Nanodrop spectrophotometer.
30. Transfer 2 µl of RNA into RNase-Free PCR tube (for RNA Electrophoresis).

**RNA Electrophoresis (~60 minutes)**

**Purpose:**

**Materials**
RNase-away
RNase-away treated Gel apparatus with combs
Gel visualizer
RNase-Free 1X TBE Running buffer
Agarose
Smart-Glow DNA Gel Dye
DNA ladder
Loading dye
RNase-Free filter P10 tips
Parafilm

**Methods:**

1) Add 1 µl of loading dye onto a strip of parafilm (1 spot/sample)
2) Pipette 2 µl gently to mix sample
3) Load dyed RNA into well
    a. Be careful agarose gels can tear easily with tips
4) Run gel at 120V for 45 min
5) Visualize gel

**cDNA Synthesis (~25 minutes)**

**Purpose:**

**Materials:**
RNase-Free Filter p10 tips
Anchored oligo-dT (6 µg/µl)
Random hexamer (6 µg/µl)
0.1 M DTT
5X Superscript buffer
10X RNase-Free dNTPs (2.5 mM stock solution, final concentration of 250 µM)
Superscript III Reverse Transcriptase
Bunsen burners/ethanol burners
RNase-Free PCR tubes (1/group)
Thermocycler
Ice/ice bin

**Methods:**

1) Prepare RNA/primer mixture in RNase-Free PCR tube:
   - 6 µl total RNA
   - 0.5 µl: anchored oligo-dT
   - 0.5 µl: random hexamer
2) Denature the primer/RNA mixture at 70°C for 10 min in a thermocycler.
3) Immediately chill on ice for >2 min.
4) Prepare reverse transcriptase cocktail in a **separate** RNase-Free PCR tube:
   - 3.75 µl 5x Superscript Buffer
   - 1.88 µl 0.1M DTT
   - 3.75 µl 10x RNase-free dNTPs
5) Add 7.5 µl of reaction cocktail to denatured RNA.
6) Take sample to Instructor to add 0.5 µl Superscript III RT.
7) Incubate on bench (25°C) for 7 min.
   a. This step is critical for allowing random hexamer to anneal to the RNA.
8) Incubate at 50°C for 2 hours in thermocycler.
9) Store samples at -20°C until next class period.

**RNA Hydrolysis and cDNA Cleanup (~25 minutes)**

**Purpose:**

**Materials:**
RNase-Free filter p10 tips
DNA clean and concentrate kit (Zymo Research #D4014)
Bunsen burners/ethanol burners
RNase-Free 1.7 ml microcentrifuge tubes
Thermocycler
Nanodrop/Qubit Spectrometer
Ice/ice bin

**Methods:**

1) Label (lid and side) of both of RNase-free 1.7 ml microcentrifuge tube and Zymo spin column.
2) Add 10 µl 1M NaOH to the cDNA synthesis reaction (PCR tube), mix by pipetting.
3) Add 10 µl 0.5M EDTA to the cDNA synthesis reaction (PCR tube) mix by pipetting.
4) Incubate at 65°C for 15 min.
5) Add cDNA (34.5 µl) sample to a new 1.7 ml tube containing 490 µl B1 (DNA Binding Buffer). Mix thoroughly by pipetting.
6) Add entire sample to Zymo-Spin column.
7) Centrifuge for 30 sec as 16,000 x g.
8) Discard flow through.
9) Add 200 µl W1 (DNA Wash Buffer).
10) Centrifuge at 16,000 x g for 30 sec.
11) Repeat wash with 200 µl W2 (DNA Wash Buffer).
12) Centrifuge at 16,000 x g for 30 sec.
13) Place the Zymo-Spin column in a clean 1.7 ml tube.
14) Add 6 µl E1 (Elution Buffer).
    a. **Note: Make sure to pipette water directly onto column resin WITHOUT touching pipette tip to membrane. If water remains on the wall of the column, tap the tube until water absorbs into the membrane.**
15) Incubate at room temperature for 1 min.

16) Centrifuge at 16,000 x g for 30 sec to elute the cDNA.

17) cDNA concentrations are calculated using a Nanodrop spectrophotometer. Store samples at -20°C.

**qPCR (~20 minutes)**

**Purpose:**

**Materials:**
Nuclease-Free water
Maxima SYBR Green Supermix (Fisher #FERK0241)
SYBR Green (Fisher #S7563)
RNase-Free filter P10 tips
RNase-Free filterP200 tips
Sterile PCR tube
Sterile 1.7 ml microcentrifuge tube
qPCR plates (Bio-Rad # MLL-9601)
qPCR membranes (Bio-Rad # MSB-1001)
Plate microcentrifuge
Real-Time PCR Machine
Biosafety cabinet
Ice/ice bin

**Methods:**

1) Label (Lid and side) PCR and 1.7 ml microcentrifuge tube.

2) Dilute the cDNA to 5 ng/µl in 10 µl NFW1 (Nuclease-Free Water) in a new PCR tube.

    **Note:** If your cDNA concentration is <5 ng/µL, just add 5µL NFW1 to your cDNA.

3) Prepare PCR mastermix (this is enough for 4 primer pairs) in a sterile 1.7 ml microcentrifuge tube.

    **Per 4 Reactions:**
    9 µl of diluted cDNA (10 ng final)
    27 µl NFW2 (Nuclease-Free water)
    45 µl Maxima SYBR Green Supermix (see Instructor for this)

4) Place PCR mastermix on ice

**The next steps are exceptionally sensitive to pipetting error and aerosolized DNA, so the Instructor will perform them in a Biosafety cabinet:**

5) Dilute the primers with Nuclease-Free water to a working stock concentration of 10 µM.

6) Prepare primer master mix:

    **Per Reaction:**
    0.4 µl Forward Primer (200 nM final)
    0.4 µl Reverse Primer (200 nM final)
    1.2 µl Nuclease-Free Water

The following are the primer pairs:

RDN18-F: CGGCTACCACATCCAAGGAA
RDN18-R: GCTGGAATTACCGCGGCT

ATF1 F: GTTTGTTGGTTTGTATCTCTTCATAAA
ATF1 R: GGGGTGAAGTCAATAAATCCA

ATF2 F: TTCTCGCAGTCTGCAGGC
ATF2 R: GCAGAACGATTCCCATTCG

BAT1 F: CTCAAGAATGGGACATCAACGA
BAT1 R: CTTGGTCAATGCACCACATTG

7) Add 2 µl of the primer mix to the ***bottom*** of the appropriate wells of a qPCR plate.

8) Add 18 µl of the PCR mix to the ***side*** of the appropriate wells. Do not change tips between samples that use the same cDNA.

    a. Careful to only touch the side of the well so as to not contaminate the pipet tip with primer mix.

    b. Change tips for non-identical samples.

9) Seal PCR plate with clear sealing membrane.

10) Briefly spin the plate in a microplate centrifuge.

11) Reaction conditions assume an optimum primer annealing temperature of ~58°C (6)

95°C -> 10 min
-------------------
95°C -> 15 sec
58°C -> 60 sec   40 cycles
-------------------
melt curve analysis (58 – 95°C in 2 sec steps)

## GC-MS

**Purpose:**

**Materials:**
Shimadzu 2020 GC/MS single quadrupole system
20 mm headspace vials with silicon PTFE septa (DWK Life Sciences Millville, NJ, USA)
10 ml serological pipet
Pipet bulb/electric pipettor

**Methods:**

1) Transfer 10 ml of supernatant into 20 mm headspace vials with silicon PTFE septa using a sterile serological pipet.
2) 1000 µl of headspace was injected in the GC equipped with a RtX-VMS 30M 0.32ID 1.8 µM column (Restek Bellefonte, PA, US).
3) GC-MS machine operation information
    a. Samples were analyzed using a Shimadzu 2020 GC/MS single quadrupole system.
    b. The injector was set to 120°C, initial oven temperature of 35°C rising at 10°C/min to 230°C hold 5 mins in constant velocity mode.
    c. The MS was operated in full scan mode.
4) Peak identification was done with spectral matching to the NIST 2014 database.

**qPCR Computer Lab**

**Purpose:**

**Information:** Critical Threshold (Ct): point at which PCR amplification exceeds a set fluorescent threshold (the number of cycles gives us this information). Lower Ct means it took less cycles to reach that threshold (higher gene expression). RDN18 (18S rRNA) gene is our control gene, which we will normalize our genes to.

**Methods:**

1) For each strain, calculate: $\Delta Ct_1$ [Ct(test gene e.g. *ATF1*) – Ct(*RDN18*)]

2) Then, calculate the median Ct value for the test gene for all strains (call this $\Delta Ct_2$)

3) $\Delta\Delta Ct = \Delta Ct_1$ [your strain] - $\Delta Ct_2$ [median of all strains]

4) Note: since PCR is exponential $\Delta\Delta Ct$ is in log space: to convert to fold change use the formula $2^{-\Delta\Delta Ct}$

#### GC-MS Computer Lab

**Purpose:**

**Information:** Similar concept of comparing your ester production to the median of all the samples. The area under the peak is proportional to the relative amount of a given compound. Thus, the log2 ratio of peak area versus the median will tell you relative abundance of each compound

**Methods:**

1) $\text{Log}_2$ transform all of the peak areas for each strain and compound.
2) Calculate the median $\log_2$ value for the esters of interest from all the strains.
3) To calculate the relative amount of ester for your strain, subtract (Your Ester of Interest) – Median (Ester of Interest)
4) To convert to fold change use the formula $2^{\text{(Your Ester of Interest) – Median (Ester of Interest)}}$

**Brewing Protocol (~3 hrs)**

**Note:** This lab is will be completed along with Yeast Viability and Counting (page C34), and Hydrometer – Original Gravity (page C35).

**Purpose:**

**Materials:**
20-Quart Stainless Steel Basic Stockpot with lid
The Brew Bag for Kettles (brew in a bag)
Binding clips (3/group)
Digital thermometer
Wort chiller
UltraPure Water (water with minimal ions)
Electric single burner hot plate
Extra-long spoons (>24 inches)
Thick kitchen gloves
Hop bags
Hops
Three-Piece Airlock (Northern Brewer Homebrew Supply SKU: 7010)
95% ethanol
Small Batch 1 Gallon Fermenter (Northern Brewer Homebrew Supply SKU: 41097)
Hand operated auto siphon with tube
Star San

**Methods:**

1) Pull brew in a bag and squeeze excess water
2) Turn hot plate up to max temp
3) Once boiling, add hops to hop bags and place in pot
4) Boil for 1 hr (lid off)
5) Using the Yeast Viability and Counting protocol, calculate the appropriate volume of cells to add
6) Add wort chiller to pot, put lid on (to prevent contamination) and turn on cold water
    a. Will take ~20 minutes to cool down
    b. Important that the wort is cooled down enough. If the wort is too warm it will kill the yeast
7) Once cool enough move onto taking hydrometer reading.
8) Add appropriate amount of cells to the 1-gallon fermenter.
9) Using an auto-siphon, fill the 1-gallon fermenter 9/10 the way full
10) Add the three-piece airlocks to 1-gallon fermenters.
11) Add 95% ethanol to airlock fill line.
12) Incubate fermenters at room temp for 3 weeks
    a. Be sure to incubate fermenters in the dark or wrap them in foil
    b. Some samples may foam up, so store all fermenters in a secondary container to catch spills

**Yeast Viability & Counting (~ 20 minutes)**

**Purpose:**

**Information:** The dye **methylene blue** can be used to measure yeast viability. Methylene blue is spontaneously oxidized in the presence of oxygen, resulting in a blue color. Cellular reductases catalyze the conversion of methylene blue to the Leuko form, which is colorless (see below). Thus, live yeast cells that contain active reductases will be colorless, while dead cells lacking the enzymes will remain blue. The number of viable cells can then be quantified using a cell counting chamber (often referred to as a **hemocytometer**).

**Materials:**
Hemocytometers
Microscopes
1.7 ml microcentrifuge tubes
Sterile 10 ml glass tubes
Sterile Erlenmeyer flasks (500 ml and 1000 ml)
0.01% Methylene Blue
Optional: Hand tally counter (VWR 23609-102)

**Methods:**

1) Add 10 µl of cells into 990 µl of methylene blue solution (0.01% (w/v)). This constitutes a 1:100 dilution.
2) Vortex briefly and pipette 10 µl of sample into the counting chamber.
3) Place the hemocytometer under a microscope at 10x magnification and find the grid pattern (see below)
4) Count cells using the four corner squares. Count both the total number of cells and the number of live (colorless) cells. For cells falling on a boundary line, only count them if they are on the top or right line (see below). Count budding cells as separate cells if the bud is at least half the size of the mother cell. **Note: try to be consistent in your counting scheme. See page B23 for a guide.**
5) Each of the nine large squares has a volume of 0.1 $mm^3$ ([1 mm x 1 mm] x 0.1 mm chamber depth). There are 1000 $mm^3$ per ml, so each square has a volume of 0.0001 ml or $10^{-4}$ ml.

   So, cells per ml = average # of viable cells (per 4 squares) x $10^4$ $ml^{-1}$ x 100 (dilution factor).

**Hydrometer – Original Gravity (~10 minutes)** (5)

**Purpose:**

**Information:** While not a perfect measure, brewers use specfic gravity (SG) to estimate the carbohydrate content of wort. Carbohydrates are the predominant dissolved solids in the wort, and similarly affect density, so SG is a reasonable approximation. SG is often measured using a hydrometer. The hydrometer is properly read at the bottom of the **meniscus**.

**Temperature Correction**

While it is best to take the reading as close as possible to the temperature for which the hydrometer was designed, the following table gives an approximate correction (in points) that must be added to the reading for 15°C. See table 3, the hydrometer temperature corrections.

Table 3

| °F | °C | Correction |
|---|---|---|
| 50 | 10 | -1 |
| 59 | 15 | 0 |
| 68 | 20 | +1 |
| 78 | 25 | +2 |
| 86 | 30 | +3 |
| 95 | 35 | +5 |
| 104 | 40 | +7 |

**Materials:**

Bunsen burners/ethanol burners
Hydrometers
Graduated cylinders (250 ml works best for minimizing loss of wort)
Digital thermometers

**Methods:**

1) Carefully pour beer into graduate cylinder
2) Measure and record the temp of the beer
3) Insert the hydrometer 3/4$^{th}$ way into the filled graduated cylinder
4) Spin and release the hydrometer
5) Read the bottom of the meniscus
6) Make sure to correct for the temperature (according to table 3)

**Hydrometer – Final Gravity (~10 minutes) (5)**

**<u>Note:</u>** This lab is will be completed along with Bottling (page C37), ABV and Percent Attenuation (page C38), and Standard Reference Method for Beer Color (page C39).

**<u>Purpose:</u>**

**<u>Information:</u>** While not a perfect measure, brewers use specfic gravity (SG) to estimate the carbohydrate content of wort. Carbohydrates are the predominant dissolved solids in the wort, and similarly affect density, so SG is a reasonable approximation. SG is often measured using a hydrometer. The hydrometer is properly read at the bottom of the **meniscus**. Note that the numbers <u>increase</u> going down. In the example below, the dashed line shows the level at which the hydrometer is read—in this case a SG of 1.037, or 37 points (aka called gravity units or GU).

**Temperature Correction**
While it is best to take the reading as close as possible to the temperature for which the hydrometer was designed, the following table gives an approximate correction (in points) that must be added to the reading for 15°C. See table 4, the hydrometer temperature corrections.

Table 4:

| °F | °C | Correction |
|---|---|---|
| 50 | 10 | -1 |
| 59 | 15 | 0 |
| 68 | 20 | +1 |
| 78 | 25 | +2 |
| 86 | 30 | +3 |
| 95 | 35 | +5 |
| 104 | 40 | +7 |

**<u>Materials:</u>**
Bunsen burners/ethanol burners
Hydrometers
Graduated cylinders (250 ml works best for minimizing loss of wort)
Digital thermometers

**<u>Methods:</u>**

1) Carefully pour beer into graduate cylinder
2) Measure and record the temp of the beer
3) Insert the hydrometer 3/4$^{th}$ way into the filled graduated cylinder
4) Spin and release the hydrometer
5) Read the bottom of the meniscus
6) Make sure to correct for the temperature

**Bottling (~15 minutes)**

**Purpose:**

**Information:** one volume is 1 L of CO2 at 1 atmosphere of pressure in 1 L of beer. Gas solubility increases at lower temperatures (We ferment at 20°C and the volumes of $CO_2$ is 0.850.) Equation for fermentation: 1 mol glucose -> 2 mol ethanol + 2 mol $CO_2$. Molecular weight of glucose = 180.16; molecular weight of $CO_2$ = 44.01. One 180.16 g of glucose (1 mole) yields 88.02 g of $CO_2$ (2 moles). One mole (44.01g) of $CO_2$ at standard pressure occupies 22.4 liters (ideal gas law). 1.96 g of CO2 occupy 1 L (44.01 g / 22.4 L), which is produced by 4.01 g glucose. Simplified: add 4.01 g glucose per L per volume.

**Materials**:
Star San (food grade, no rinse sanitizer)
Swing neck amber bottles (clear requires fermenting in the dark – prevent "skunking" of the beer) (2/brewing group)
10 ml serological pipettes (2/brewing group)
Pipette bulbs (or automatic pipettors)
Sterilized 50% corn sugar (50% glucose)
Hand operated auto siphon with tube (1/brewing group)
Ethanol burners/Bunsen burners
95% EtOH
Sterile 50 ml conical tube (1/brewing group)

**Methods:**

1) Look up the recommended volumes of $CO_2$ of your beer style
2) 2.9 volumes (depend on your type of beer) – 0.85 volumes (see information section) = 2.05 volumes (the amount we want to add)
3) Assume bottles are 16 ounces or 0.47 liters
4) 2.05 volumes x 0.47 liters x 4.01 g glucose (see information section) = 3.86 g glucose
5) Our glucose stock is 50% w/v (or 0.5 g glucose/ml), so 3.86 g (must divide by 0.5 to calculate volume to add) = 7.72 ml
6) Under a flame, add the appropriate volume of 50% glucose to bottle
7) Using auto-siphon (be careful to not disturb the cells at the bottom) carefully fill glucose containing swing-neck bottle to 2 inches below the stopper.
8) Incubate at room temp (in the dark) for 2.5 weeks.
9) Using auto-siphon fill a 50 ml conical tube to be used for SRM color assay and Hop bitterness assay.

**ABV and Percent Attenuation (~ 5 minutes)**

**Purpose:**

**Information:** Alcohol by volume (ABV) can be estimated using specific gravity (density of a solution relative to water). Water inherently has a specific gravity of 1.000 (density of water/ density of water). Sugars have a higher density and ethanol has a lower density (specific gravity = 0.794). Thus, a simple formula to estimate ABV is to assume that the change in original gravity (OG) to final gravity (FG) is due to ethanol:

**More Accurate Equation**
**ABV = [76.08 x (OG-FG) / (1.775 - OG)] x (FG / 0.794)**

**where [76.08 x (OG-FG) / 1.775 – OG] represents the alcohol percentage by weight, and 0.794 denotes the specific gravity of ethanol**

Apparent attenuation is the percentage of sugars that the yeast consumed, which can be also be calculated using the change in OG to FG:

**Apparent Attenuation (%) = ((OG-FG) / (OG-1)) x 100**

The following are common yeast attenuation ranges:

Low = 65-70%
Medium = 71-75%
High = 76-85%

**Standard Reference Method for Beer Color (~5 minutes)**

**Purpose**:

**Information:** The Standard Reference Method (SRM) for measuring beer color was adopted by the American Society of Brewing Chemists in 1950 as an alternative to the Lovibond system. The SRM method relies on visible absorbance at 430 nm.

**Materials:**
P1000 tips
Spectrometer
Cuvettes (2/brewing group)
Sterile water

**Methods:**

1) Pipette 1 ml of beer into a cuvette (If the beer is dark, you will need to make a 10-fold dilution).

2) Add 1 ml of blank (sterile water) to a cuvette.

3) Measure the absorbance of the sample at 430 nm.

4) SRM = Absorbance at 430 nm x 12.7 x dilution factor.

**Hop Bitterness (~12-90 minutes) (7)**

**Purpose:**

**Information:** The alpha acids in hop flowers are isomerized during wort boiling to form **iso-alpha-acids**. Humulone shown below is the most prevalent alpha acid in hops.

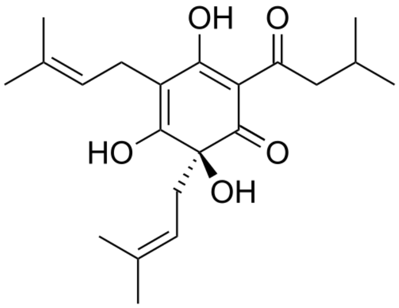

humulone

Image taken from Wikipedia

The hop iso-alpha-acids contribute to the bitterness of the beer. The amount of iso-alpha acids is generally reported on the **International Bitterness Units** (IBU) scale. One IBU equals 1 mg of isomerized alpha acid per liter. Because iso-alpha-acids absorb strongly at 270 nm, a spectrophotometer can be used for quantification.

Hop iso-alpha-acids are slightly hydrophobic, and their hydrophobicity increases at low pH. Acidifying the beer allows iso-alpha acids to be extracted into hydrophobic organic solvents. Iso-octane is the standard solvent used in the brewing industry, since it has a low absorbance at 270 nm.

IBU is calculated as the absorbance multiplied by a factor of 50. This factor is derived from the dilution factor and an assumption that 70% of the absorbance of the iso-octane solution is due to iso-alpha-acids.

**Warning: iso-octane is highly flammable and toxic by inhalation, ingestion, or skin absorption. Hydrochloric Acid is highly corrosive and toxic by inhalation, ingestion, or skin absorption**

**Materials:**
50 ml conical tubes (2/brewing group, 1 for Instructor to make the blank)
Microcentrifuge tubes (for octyl alcohol and 3N HCl)
Octyl alcohol
Iso-octane (2,2,4-trimethylpentane)
3 N hydrochloric acid
P1000 tips
10 ml serological pipettes
Pipette bulbs (or automatic pipettors)
Vortex mixers
UV Spectrometer
Quartz cuvette (Fisher NC9416015) – fragile, only Instructor handles

Optional materials:
Table top centrifuge capable of spinning 50 ml conical tubes

**Methods:**

1) Pipette 10 ml of beer into a 50 ml conical tube containing 20 ml iso-octane.
2) Add 50 µl octyl alcohol and 1 ml 3 N hydrochloric acid.
3) Close the tube (make sure it is well sealed) and vortex for 15 min.
4) Centrifuge at 3000 x g for 5 min to allow phase separation to occur (or allow gravity to separate the phases generally 60 min), **in the dark (iso-alpha-acids are UV sensitive).**
5) Add 1 ml of blank (20 ml iso-octane plus 50 µl octyl alcohol) to a UV transparent cuvette (quartz).
    a. Note: the iso-octane will cloud up plastic cuvettes
6) Transfer 1 ml of the sample (upper organic phase) to a cuvette.
7) Measure the absorbance of the sample at 275 nm.
8) IBU = Absorbance at 275 nm x 50.

Notebook Evaluation Form

**Table of contents (5%)** ____________

List of all experiments and correct page numbers

**Rationale (20%)** ____________

Rationale explaining broad reason as to why we are performing the research and the hypothesis

(i.e. Samples have been collected from the wild. These samples could include wild yeast isolates that could potentially be good at fermenting and produce high levels of ethanol. These wild isolates will be tested to see how they compare to commercial brewing strains. I hypothesized that I will successfully isolate wild yeast.)

**Methods (20%)** ____________

Detailed methods of what exactly was performed

**Results (20%)** ____________

Does every experiment have the correct results?
Every image/table must be present.
Is their sample clearly labeled?
Do not discuss the results (that is for the discussion section)

**Discussion (15%)** ____________

Interpret and discuss the results.
Do your results support your hypothesis?

**Organization (15%)** ____________

Are the experiments in the order they were performed?
Is the notebook legible (if handwritten)?

**Dated (5%)** ____________

Are experiments dated?

Additional comments:

Example Lab Notebook Entry

**Wild Yeast Collection: 08/26/17**

Rationale:

We will collect samples from the environment to isolate wild yeast from. These wild yeasts could potentially be good fermenters. These wild isolates will be tested to see how they compare to commercial brewing strains for brewing a small batch of beer. I hypothesize that I will collect yeast from the wild.

Materials and Methods:

- Where to look for wild yeast
  - Try to go off the beaten path, away from areas that are likely to be contaminated by human activities (or spilled beer)
  - Some good places to look:
    - Under wild fruit or berry bushes or trees – especially where berries have fallen into the soil and are starting to rot
    - Soil under trees – especially if they are injured and leaking sap into the soil

- How to collect sample
  1. Put gloves on
  2. Open the sterile tube – make sure you don't touch the inside of the lid or tube with your hands.
  3. Scoop up a little soil into the tube (about to the 5mL mark). If there are rotten berries in the soil, you can scoop one or two of them up too.
  4. Seal the tube and store at room temperature until you bring the sample back into lab.
  5. Record as much information about the sample collection location as possible.

Results:

BC19 was collected on 08/27/17. Sample was located close to the beach in St. Petersburg, FL.
BC20 was collected on 08/27/17. Sample contains small berry and soil sample. Sample was taken from Skull Creek Trail, Fayetteville, AR
BC21 was collected on 08/28/17. Sample contains berries and soil sample. Sample was taken from campus near Poultry Science building, Fayetteville, AR

Discussion:

Samples were stored at room temp until the next class period. These are the samples I will continue forward with to the wild yeast isolation protocol. They will be referred as BC19, BC20, and BC21.

**Tips on How to Give a Scientific Presentation**

6) Reasons to Give a Scientific Presentation: 1) You are compelled to
   7. For practice (you may be doing this a lot!)
   8. To get a job
   9. To teach people stuff
   10. To become better known in your field
   11. To get people to read your stuff
   12. To gain experience

7) Basic Principles:
   8. Know your audience (don't overestimate their knowledge)
   9. Take it seriously (fair or not, you are being judged)
   10. Simplify when possible
   11. Tell a story
   12. Know yourself (adopt another's style at your own peril)
   13. Learn from other people's talk
   14. Images over text

8) General Advice:
   6. Humor is okay if that's your style, but be careful with forced or rehearsed jokes
   7. Avoid jargon and technical detail, always stay with the big picture
   8. Summarize often
   9. Don't over-speculate – it's more than okay to admit ignorance
   10. Don't go over the time limit!

9) Ways to be annoying:
   8. Unlabeled graphs or meaningless labels Lots of words, complete sentences, or better yet whole paragraphs
   9. Read the text verbatim
   10. Skip the rationale for the talk
   11. Mumble
   12. Use gratuitous animations
   13. Flash your pointer everywhere
   14. Use excessive jargon and abbreviations

10) Final Tips:
   7. Practice talks by volunteering at every opportunity
   8. Rehearse, but don't let it kill your enthusiasm
   9. Try to visualize your target audience when preparing
   10. Focus on transitions and tricky parts
   11. Have the opening minutes down – 1$^{st}$ 2-3 slides
   12. Exude enthusiasm and confidence

Note: This form was also used by students for peer evaluation.

**Presentation Evaluation Rubric**

**Presenter: ________________________________** **Date: _______________**

Please complete this evaluation by rating the presentation in each category:

**1 = needs improvement; 2 = fair; 3 = good; 4 = excellent**

**Introduction**
4: Clearly explains the topic and provides relevant background
3: Clearly introduces the topic, but some context is lacking
2: Topic is vaguely introduced, and the relevance is not clear
1: No clear topic is presented

**Content**
4: Clearly presents useful information in an organized way
3: Presents useful information, but may be slightly unclear or unorganized
2: Information presented is very unclear
1: Very little information presented

**Conclusions**
4: Clearly conveys the big-picture message and meaning
3: Clearly summarizes information, but main point or importance is unclear
2: Information is poorly summarized
1: No clear conclusion or main message

**Organization**
4: Clear evidence of speaker preparation, visuals and organization enhance presentation
3: Clear evidence of preparation, visuals and organization are adequate
2: Preparation, visuals, and/or organization are lacking
1: Lack of preparation, disorganized

**Presentation**
4: Engages audience, easily understood, fields questions well
3: Easily understood, but may not engage audience or field questions well
2: Difficult to understand, little audience engagement or ability to field questions
1: Very difficult to understand, poor attempt to engage audience or answer questions

***What were the strengths of the presentation?***

***What could be improved?***

### How to Write a Scientific Paper

**Purpose of the Exercise**
The primary purpose is to give you experience crafting a clear and logically compelling research article. Effective writing is an important component of scientific communication. A well-written scientific paper should clearly describe the rationale, procedures, and results for each of the experiments that were performed, and then place those results into the context of our existing knowledge.

**Paper Requirements**
The paper should be at least 10 pages (single-spaced, 11 or 12 point font), with at least 10 references from academic sources, and can be used to fulfill the Fulbright College Senior Writing Requirement.

**Paper Organization and Specifics**

1) Paper Title and Student ID.

2) <u>Abstract (~250 – 300 words):</u>
This should be a single paragraph that summarizes the major aspects of the entire paper. This includes the motivation or rationale for the experiments, the major research outcomes, and interpretations or conclusions.

3) <u>Introduction:</u>
This section should give the reader an understanding of the general background literature, while building a case for why the reader should get excited about the experiments performed. This should include the scientific rationale underlying the main questions and/or hypotheses. Background information (generally from the scientific literature) should have enough detail to make the "Results" and "Discussion" understandable to the reader (clarifying figures and cartoons are often helpful).

4) <u>Materials and Methods:</u>
This section should provide enough information about the experimental procedures to allow another scientist to reproduce them. Certainly include references if the methods are being adapted from another source, but make sure this section provides enough detail that the reader does not have to sift through other papers to figure out what you did. This section should be written in paragraph form and not bullet points like a protocol.

5) <u>Results:</u>
Summarize the findings for each experiment. Use figures (e.g. graphs) and/or tables as is appropriate, and make sure that the text clearly states the main findings. All tables and figures must be referred to in the text. Tables should have a "header" or title. Figures should include clear legends and must also have a caption below. All tables and figures should be understandable without having to refer to the main text body. It is helpful to remind the reader of the rationale for each experiment.

6) <u>Discussion:</u>
This section should provide interpretations of the results, and place those results into a broader context. It is helpful to briefly summarize the main findings from the results, but do not simply repeat what has been written in the "Results" section. In this section, it would be appropriate to re-address the questions/hypotheses from the introduction in light of the data. When interpreting the data, think about whether there are alternative explanations, and how the data fit into the existing literature.

7) <u>References:</u>
The citations should be listed at the end. This can be in any standard journal format that includes the title of the article and journal (or book), and the names of all of the authors (if there are more than five authors, list the first five and then add "*et al.*" for the others). <u>This is not part of the page count.</u> The style of "Smith *et al.* 2008" is preferred for in-text citations, though you can see that numbered citations are used by some journals.

**General Suggestions**

1) It is up to you how you want to organize your paper, and what experiments to include, but it's important to have a cohesive narrative. One suggestion would be to focus on comparing and contrasting wild vs. commercial yeast strains during brewing.

2) Take advantage of the draft. This is ungraded (i.e. credit/no-credit) and a chance to get some feedback on what to improve for your final paper.

3) Write up the "Results" section first. This will allow you identify topics for the "Introduction" and "Discussion" sections that should be provided for understanding the context of your results.

4) "Introduction" sections frequently contain information that is not relevant to the overall theme of the paper. Remember, the point is to provide sufficient background to understand the motivation for the experiments, and not to describe the entirety of a given scientific field.

5) Try to be clear in your writing. Scientific reasoning can be challenging to follow. Make sure to only have one major idea per sentence. Active voice is easier to read than passive voice, and it's okay to use first person ("I" or "we"). Topic sentences from each paragraph should let the reader know where your thoughts are going, and the last section of a paragraph should frequently have some sort of conclusion (Look at the research papers assigned during the semester to get an idea about how paragraphs are structured).

6) Things literally done in the past should be written using the past tense. This will include the Methods and Results sections.

7) Do run spellcheck and make sure the paper looks thoughtfully put together. Typos are not explicitly a criterion for grading, but they necessarily affect the readers' impressions.

**Grading**

The primary criterion for grading is whether you craft a well-researched, well-organized, and logically presented paper. Explicitly, I am looking for clarity of ideas, and thoughtful presentation of the data and its interpretation. This is meant to be a useful exercise, and you will have feedback on the ungraded first draft to give you ample opportunity to improve your final drafts.

**Paper Evaluation Rubric:**

| Sections | Points | Criteria | Sum |
|---|---|---|---|
| **Abstract** | | | 5 |
| background | 1 | *context, existing knowledge, relevant information* | |
| main objectives | 1 | *rationale for study is clear* | |
| approach | 1 | *general approaches / experiments stated* | |
| results and conclusions | 1 | *clearly explained and summarized* | |
| narrative | 1 | *concise, clear logic* | |
| **Background and Significance** | | | 20 |
| rationale | 6 | *motivation for the study, hypotheses, and or objectives* | |
| background | 6 | *existing knowledge, gaps in the research* | |
| narrative | 8 | *concise, clear logic* | |
| **Materials and Methods** | | | 20 |
| protocols and approaches | 10 | *clear descriptions of materials and methods, paragraph form* | |
| narrative | 10 | *concise, clear logic* | |
| **Results** | | | 20 |
| research objectives | 6 | *describes experimental objectives and overall approaches* | |
| results | 7 | *sufficiently detailed and comprehensive, including tables/figures* | |
| narrative | 7 | *concise, clear logic* | |
| **Discussion** | | | 20 |
| summary and interpretation | 6 | *summary of outcomes, what results mean* | |
| conclusions and context | 6 | *fit with other research, possible future directions* | |
| narrative | 8 | *concise, clear logic* | |
| **References** | | | 5 |
| appropriate | 2 | *primary literature, relevance* | |
| comprehensive | 2 | *all aspects/methods referenced?* | |
| format | 1 | *formatted properly (Journal style)* | |
| **Final Version** | | | |
| revisions completed | 90 | | |
| proposal improved | 10 | | |

| | |
|---|---|
| Total = | **100** |

Homework #1

1) Every student will be given a wild yeast collection kit. Take a pen with you and be sure to carefully record on the provided card where you isolated the sample from (e.g. Fayetteville AR, rotting blueberries from Finger Park).

Homework #2

**You can either hand in your typed answers in class, or email them to me. The only identifying information should be your student ID.**

General Rules: 1) I strongly encourage people to work together in groups, but each person must generate their own answers. Directly copying answers is considered plagiarism. 2) Type your answers and try to be concise. Most questions can be answered in 2-3 sentences, unless otherwise stated. 3) I am happy to field any questions or concerns during class or by email.

…………………………………………………………………………………………………………………...

***The following questions refer to the Sniegowski et al. paper (Week 2 reading):***

1. **(6 points)** The authors describe a 2-step enrichment of yeast from natural substrates, with one of the major differences being the choice of carbon source (sucrose for liquid enrichment, followed by methyl-a-D-glucopyranoside on solid media—use PubMed and/or Google Scholar to identify yeast papers that reference this compound)

   a) What is methyl-α-D-glucopyranoside?

   sugar, oligosaccharide, glucose derivative, carbon source

   b) What is a major benefit to using this 2-step procedure to enrich for yeast?

   relatively few non-yeast can grow on methyl-glucopyranoside; 2-step decreases number of contaminants

   c) What is a potential downside for using methyl-α-D-glucopyranoside in the second step?

   some strains unable to grow on methyl-glucopyranoside

2. **(4 points)** The authors tested spore viability in hybrids of their newly isolated wild strains when crossed to a European tester strain (Table 2).

   a) Which species had lower hybrid spore viability?

   Paradoxis

   b) How might long periods of reproductive isolation lead to yeast hybrid infertility (at the genetic and level)?

   changes in chromosome numbers, chromosomal mutations/rearrangements, incompatible mutations/alleles…

3. **(4 points)** The authors determined whether their isolates were *S. cerevisiae* or *S. paradoxus* by separating the chromosomes using a CHEF gel and then detecting probes that hybridize to Ty1 elements.

   a) What is CHEF gel electrophoresis, and how does it work to separate chromosomes on a gel?

   type of pulse-field electrophoresis, where gel is moved to wriggle DNA through, normal electrophoresis cannot separate large fragments

   b) What is the rationale of using Ty1 elements to classify the wild yeast isolates into species?

   transposons can move (mobile DNA), ending up on different chromosomes across species time

4. **(6 points)** The Ty1 hybridization analysis only works if you have known DNA standards to compare to (the authors had known *S. cerevisiae* and *S. paradoxus* isolates). In the case of your wild yeast collections, there may be previously unidentified yeast species or species that we do not have access to. So, we need another strategy that does not require having known DNA standards.

   Suggest an alternative approach to identify the species of your wild yeast isolates. This should not be a detailed protocol (we will work through that in class), but should describe the big picture of how the approach can be applied to identify new yeast species.

   something specific that would work for unknowns (e.g. gene specific PCR, gene/genome sequencing)

Homework #3

**You can either hand in your typed answers in class, or email them to me. The only identifying information should be your student ID.**

General Rules: 1) I strongly encourage people to work together in groups, but each person must generate their own answers. Directly copying answers is considered plagiarism. 2) Type your answers and try to be concise. Most questions can be answered in 2-3 sentences, unless otherwise stated. 3) I am happy to field any questions or concerns during class or by email.

.................................................................................................................................................................

***The following questions refer to the Sylvester et al. paper (Week 3-4 reading)***

1. The authors argue that the main natural habitat of North American *Saccharomyces* is not closely associated with oak trees?

    a) **(2 points)** What data do they use to support this argument?

    failure to identify certain *Saccharomyces* species on oak substrates, despite large sample sizes (statistically significant) and the ability to identify other genera on oak (the lone exception was *S. paradoxus*)

    b) **(2 points)** If *Saccharomyces* is not closely associated with oak trees, what is one hypothesis for why *Saccharomyces* can still be isolated from oak trees at low numbers?

    the yeast end up there incidentally by being vectored (possibly by insects, for example)

    c) **(4 points)** An alternative hypothesis is that *Saccharomyces* is truly associated with oak trees, but not in the Wisconsin/Michigan/ or Washington/Oregon sites that were heavily sampled in this study. What about the ecological preferences of *Saccharomyces* might cause this lack of association with oak trees at these more Northern latitudes? Where would you sample extensively to test this hypothesis?

    higher temperature preferences by *Saccharomyces* yeast, which can be tested by sampling at lower latitudes that have warmer climates (and perhaps milder winters)

2. **(3 points)** The authors were interested in understanding the population genetics of *S. paradoxus* strains, so they used the *POP2* and *RPB2* genes to make population assignments. Why is ITS sequencing sufficient for assigning species, but insufficient for population (strain) level analysis?

    ITS can resolve closely related species because there is enough sequence divergence; however, within a species ITS sequences do not vary enough to differentiate populations, so markers (genes) with more sequence divergence are necessary

3. You are given DNA isolated from an unknown budding yeast sample. You perform PCR on the ITS sequence, and after confirming a single band, PCR purify the DNA for sequencing.

    a) **(3 points)** Before proceeding to sequencing, you measure the absorbance of the DNA in a spectrophotometer. Specifically, you measure the absorbance at 230 nm, 260 nm, and 280 nm. What biological molecules relevant to DNA purification absorb at each of those wavelengths?

    230 nm: common contaminants (chaotropic salts, phenolics…)
    260 nm: DNA (specifically the heterocylic rings of the nucleotides)
    280 nm: protein (specifically aromatic amino acids, especially tryptophan)

b) **(3 points)** You obtain a 260/280 ratio of 1.52. Do you proceed with the sequencing or repeat the PCR and purification? Why?

repeat purification due to protein contamination (which may interfere with downstream sequencing)

4. You obtain the following ITS sequence from the unknown yeast in Question 3:

GATCATTACTGATTTGCTTAATTGCACCACATGTGTTTTTCTTTGAAACAAACTTGCTTTGGCGGTGGG
CCCAGCCTGCCGCCAGAGGTCTAAACTTACAACCAATTTTTTATCAACTTGTCACACCAGATTATTACT
TAATAGTCAAAACTTTCAACAACGGATCTCTTGGTTCTCGCATCGATGAAGAACGCAGCGAAATGCGA
TACGTAATATGAATTGCAGATATTCGTGAATCATCGAATCTTTGAACGCACATTGCGCCCTCTGGTATT
CCGGAGGGCATGCCTGTTTGAGCGTCGTTTCTCCCTCAAACCGCTGGGTTTGGTGTTGAGCAATACGAC
TTGGGTTTGCTTGAAAGACGGTAGTGGTAAGGCGGGATCGCTTTGACAATGGCTTAGGTCTAACCAAA
AACATTGCTTGCGGCGGTAACGTCCACCACGTATATCTTCAAACTTTGACCTCAAATCAGGTAGGACT
ACCCGCTGAACTTAAGCATATCAATAAGCGGAGGAAAAGAAA

a) **(2 points)** Very briefly describe how you would analyze the sequence (what software you are using and what settings).

Blast (nucleotide, default settings, optionally do not filter low complexity regions)

b) **(2 points)** Perform the analysis above and report the closest species "hit."

*Candida sp. (in: Saccharomycetales)* strain LEMI7885

c) **(2 points)** Unfortunately, the person who handed you the yeast DNA cannot remember where the sample came from (alas, they were not given superb training in how to keep a laboratory notebook, unlike you). Because you would like to grow more of the yeast to characterize further, what temperature would you use for optimal growth?

~22°C based on *Candida sp.* in the paper. Those who noted that this was identical to *Candida albicans* and would grow best at 33°C got an extra point

Homework #4

**You can either hand in your typed answers in class, or email them to me. The only identifying information should be your student ID.**

General Rules: 1) Unlike previous problem sets, I expect students to perform the following research independently. 2) I am happy to field any questions or concerns during class or by email.

The lab has done an admirable job in collecting new wild yeast strains, the vast majority of which are not *Saccharomyces cerevisiae* (Brewer's yeast). For this exercise, students should perform a literature search to research a **non-*cerevisiae*** yeast species. The species you choose to perform a literature search can be the one you personally isolated, or a strain isolated by your lab partner of another student. The only rule is that you should research a species other than *S. cerevisiae.*

PubMed and Google Scholar are excellent tools for finding the relevant literature.

Please provide a short review of the relevant literature concerning the species. I expect this to be ~1 page total, single-spaced (11- or 12-pt Arial or Time New Roman—no funny business with font). Please provide references for specific facts, and please refrain for citing word-for-word. This is an exercise in putting the relevant literature into your own words (which will be good practice for your final paper).

Specifically, I'm looking for 1-2 paragraphs on each of the following topics (noting that for some organisms not everything will be known—just make sure to say as much):

**1.** It's history and life history. For example where else has the organism been isolated? What is known about its natural ecology? It's evolutionary history (where does it fit into the fungal tree of life—who are the closest relatives)? And anything else that paints a picture of the background of this organism.

**2.** General characteristics of the organism, including but not necessarily limited to its morphology, physiology, and biochemistry (especially if anything unique is known about its metabolic capabilities).

**3.** Biotechnological applications for the organism. Is anything known about this organism's association with humans? Has it been used for brewing in the past or present? What is this organism potentially good at doing that could be useful for brewing or even other biotechnological applications?

Homework #5

**This is a <u>group assignment</u> (four students per group). Please have one designated student email the completed assignment to me. I'll print out the assignment, which we will use to during our group discussion with Ron Schmidt from Core Brewing.**

The goal of this exercise is to scientifically design a brewing recipe. Specifically, I want you to learn about the properties of different ale styles you may want to brew, how to fit an ale style to the particular properties of your yeast strains, and how to construct a recipe.

**1.** Throughout the course, each group of four students will have characterized one wild yeast strain and one commercial yeast strain. Based upon the entire set of class data for all strains, estimate or predict whether your yeast strains are below average, average, or above average:
   i. producers of esters
   ii. producers of fusel alcohols
   iii. fermenters (fermentation rate as measured by the class)
   iv. attenuators (for commercial strains)

**2.** Identify 4 different **<u>ale</u>** styles as a topic of research. For each style, write a short paragraph about the defining characteristics. (Note: I picked 4 styles so that each individual in the group can write a paragraph, contribute, and share).

**3.** Based on the properties of your yeast (either wild or commercial), choose an appropriate ale style for brewing. Briefly describe why your choice of style fits your yeast (i.e. what yeast characteristics are important for your ale style?)

Based on your choice from Question 3 above, begin to design a brewing recipe by answering the following questions.

**4.** What are the target parameters you will build your style around? Specifically what is your target…
   a. alcohol by volume (ABV)?
   b. color (either degrees lovibond or the standard reference method (SRM))?
   c. bitterness (IBUs)?
   d. final pH?

**5.** Selection of fermentables?
   a. What will be your base malt, and in what percentage of total grains?
   b. What will be your specialty grain(s), and in what percentage of total grains?

**6.** Selection of other specialty ingredients?
   a. What (if any) adjuncts or non-grain specialty ingredients will be included, and in what percentage?

**7.** What are your mashing parameters (temperature(s) and time)?

**8.** Selection of hops?
   c. What hop varieties will you add?
   d. At what time(s) during the brewing process will you be adding hops?

**Pre/Post-Exam Questions**

**1.** What are the primary products of yeast fermentation?
   A. ethanol and carbon dioxide.
   B. ethanol and pyruvate.
   C. pyruvate and carbon dioxide.
   D. lactate and carbon dioxide.
   E. I do not know.

**2.** Which of the following is not a characteristic of lager yeast?
   A. They ferment well at low temperatures.
   B. They exhibit high genetic diversity compared to ale yeast.
   C. They are hybrids of *Saccharomyces cerevisiae* and *Saccharomyces eubayanus.*
   D. They display poor sporulation and mating ability.
   E. I do not know.

**3.** Which of the following is not a component of PCR?
   A. dideoxy-dNTPs (ddNTPs).
   B. a thermostable DNA polymerase.
   C. primers.
   D. template DNA.
   E. I do not know.

**4.** Which step of the brewing process describes the conversion of starches into sugars utilizable by yeast?
   A. milling.
   B. malting.
   C. mashing.
   D. sparging.
   E. I do not know.

**5.** Hop iso-alpha acids are most effective in controlling the growth of…
   A. gram-positive bacteria.
   B. gram-negative bacteria.
   C. yeast.
   D. filamentous fungi.
   E. I do not know

**6.** True or false: Fermentation generates more ATP than respiration.
   A. True.
   B. False.
   C. I do not know.

**7.** True or false: *Saccharomyces cerevisiae* is able to ferment glucose both aerobically and anaerobically.
   A. True.
   B. False.
   C. I do not know

**8.** True or false: When using BLAST, lowering the E value threshold will result in more matches.
   A. True.
   B. False.
   C. I do not know.

**9.** True or false: Mashing at higher temperatures will result in lower levels of utilizable sugars.
   A. True.
   B. False.
   C. I do not know.

**10.** True or false: Attenuation refers to the ability of certain yeast strains to aggregate.
   A. True.
   B. False.
   C. I do not know.

*Short Answer Questions:*

**11.** List two ingredients in media for yeast enrichment and describe their function.

EtOH -> prevents growth of EtOH-sensitive microbes
antibiotic -> prevents bacterial growth
sucrose -> carbon and energy source
yeast extract -> vitamins and other micronutrients
peptone -> nitrogen source
yeast nitrogen base -> nitrogen source, vitamins
amino acid cocktail -> amino acids for biosynthesis
alpha-D- methylglucoside -> stringent carbon and energy source
agar -> solid substrate

**12.** Why is the ITS region used for fungal phylogenetics?

contains enough sequence differences to resolve closely related fungal species

**13.** During PCR of the ITS region from an unknown yeast species, you notice two bands. What is a possible explanation?

sample has a single hybrid species
sample contaminated, has two different species present

**14.** List the four major ingredients necessary for brewing.

water, grain (or malt), hops, yeast

**15.** Describe how beer bitterness (in IBU) can be measured.

extract iso-alpha acids and measure absorbance

**Pre/Post-Survey Questions**

**1.** I know where to find yeast in natural environments, and would be able to isolate wild yeast.

A. strongly disagree
B. disagree
C. neither agree or disagree
D. agree
E. strongly agree

**2.** If given an unknown yeast isolate, I would be able to use molecular biology and phylogenetics to determine which species it is.

A. strongly disagree
B. disagree
C. neither agree or disagree
D. agree
E. strongly agree

**3.** I am comfortable reading and discussing scientific research papers (aka the primary literary).

A. strongly disagree
B. disagree
C. neither agree or disagree
D. agree
E. strongly agree

**4.** I can describe all of the major steps in brewing.

A. strongly disagree
B. disagree
C. neither agree or disagree
D. agree
E. strongly agree

**5.** I am able to brew beer on my own.

A. strongly disagree
B. disagree
C. neither agree or disagree
D. agree
E. strongly agree
